# Integrated QTL mapping and CRISPR screening in pooled iPSC-derived microglia reveals genetic drivers of neurodegenerative risk

**DOI:** 10.1101/2025.08.18.670767

**Authors:** Marta Perez-Alcantara, Sam Washer, Yixi Chen, Juliette Steer, Daianna Gonzalez-Padilla, Joe McWilliam, David Willé, Nikos Panousis, Peep Kolberg, Elena Navarro Guerrero, Kaur Alasoo, Hazel Hall-Roberts, Julie Williams, Sally A. Cowley, Gosia Trynka, Andrew Bassett

**Affiliations:** Wellcome Sanger Institute, Wellcome Genome Campus, Hinxton, CB10 1SA, UK; Open Targets, Wellcome Genome Campus, Hinxton, CB10 1SA, UK; James and Lillian Martin Centre for Stem Cell Research, Sir William Dunn School of Pathology, University of Oxford, South Parks Road, Oxford, OX1 3RE, UK; UK Dementia Research Institute at Cardiff University, Maindy Road, Cardiff CF24 4HQ, UK; GSK Research, GSK Medicines Research Centre, Gunnels Wood Road, Stevenage, SG1 2NY, UK; Institute of Computer Science, University of Tartu, Tartu, 51009, Estonia; Target Discovery Institute, University of Oxford, Oxford, OX3 7FZ, UK

**Author notes:** Joint first authors (Marta Perez-Alcantara, Sam Washer). Joint senior and corresponding authors.

## Abstract

Mounting evidence implicates microglia in neurodegeneration, but linking disease-associated genetic variants to target genes and cellular phenotypes is hindered by the inaccessibility of these cells. We differentiated 261 human iPSC lines into microglia-like cells (iMGL) in pools with phenotypic (differentiation, phagocytosis and migration) and single-cell transcriptomic readouts. Burden analysis of deleterious variants detected 36 genes influencing microglial phenotypes. Expression quantitative trait locus (eQTL) analysis found 7,121 eGenes, and 79 colocalizations across four neurodegenerative disease GWAS, half of which had limited prior evidence of causality. Integration of eQTL and phenotypic associations highlighted the role of disease-relevant variants including *LRRK2* and *TREM2* acting via microglial phagocytosis. A coupled CRISPR screen identified a role of *TREM2* in phagocytosis and highlighted the importance of cellular state in directionality of phenotype. By contextualizing variant effects within disease-relevant microglial states, we provide a comprehensive framework for interpreting the function of risk loci in neurodegenerative disorders.

## Introduction

Genome-wide association studies (GWAS) have identified thousands of genetic variants linked to complex traits and diseases, but translating these into biological insight remains challenging. Most variants are non-coding and likely influence disease by regulating gene expression, often in specific cell types, states, or environments.

While expression quantitative trait locus (eQTL) mapping helps link regulatory variants to their target genes, it does not reveal the downstream cellular consequences. Similarly, population-scale mapping of genetic effects on cellular traits links genetic variation to functional outcomes, however, such studies have been limited by practical challenges: disease-relevant cell types are often difficult to access, and detecting modest genetic effects on complex cellular traits typically requires large sample sizes.

Human induced pluripotent stem cells (hiPSCs) offer a scalable platform to model human biology *in vitro*, enabling access to otherwise inaccessible cell types, like those of the brain. They are also suitable for exploring how natural genetic variation influences gene expression and cellular traits independent of uncontrolled environmental effects, and provide a foundation for targeted functional follow-up using CRISPR perturbations, enabling stronger causal inference for variant-to-gene-to-phenotype relationships^1–6^.

Pools of iPSCs enable the use of natural genetic variation while enhancing experimental scalability and reducing differentiation variability and batch effects. Recent studies have leveraged large numbers of donors to investigate how genetic variation influences iPSC-derived neuronal responses to stress and viral susceptibility^6^, demonstrating links between genetic background, gene regulation, and cellular phenotypes, identifying hundreds of eQTLs that colocalize with disease-associated loci from GWAS.

Despite these advances, pooled iPSC-based approaches present several technical challenges. Both somatic and germline genetic variants can affect differentiation efficiency, resulting in imbalanced representation of donors within a pool. This introduces variability that can confound downstream analyses. Additionally, independent differentiation pools are still prone to batch effects, complicating data integration across experiments^6^. Improving our understanding of the factors that drive efficient and consistent differentiation is essential for scaling these models and maximizing their utility across the research community.

Microglia have been implicated in neurodegeneration through their involvement in neuroinflammation, synaptic pruning, and clearance of pathological proteins. GWAS have identified hundreds of loci altering risk of neurodegenerative disease, and disease-associated variants, especially in Alzheimer’s disease (AD), are enriched in microglial-specific chromatin regulatory regions and genes^7–10^. Rare variants have further implicated microglia as a key cell type, e.g. the R47H mutation in microglia-specific *TREM2* gene increases risk of AD^11^. However, the precise impact of the common variants linked to neurodegenerative disease on gene expression and microglial phenotypes mostly remains poorly understood, in part, due to the inaccessibility of the primary human cellular material. Human iPSC-derived microglia-like cells (iMGL) have emerged as valuable cell models to study molecular mechanisms dysregulated by disease variants.

Here, we develop an experimental framework of pooled iMGL from 261 lines to map the impact of genetic variation on microglial molecular and cellular phenotypes. By combining single-cell transcriptomics in naive and stimulated states with functional assays measuring differentiation, phagocytosis, and migration, we capture a wide spectrum of microglial responses. We identify both common and rare variant effects through integration of genotype information, single-cell QTL mapping, genome-wide and gene-based association testing, and independent functional identification and validation of key regulators using CRISPR screens. Our suite of assays contextualises variant effects within microglial states relevant to disease enabling functional interpretation of risk variants implicated in neurodegeneration.

## Results

### Single-cell eQTL mapping in resting and stimulated iMGL

We differentiated 16 pools totalling 261 iPSC lines (247 HipSci^12^ from healthy donors and 14 IPMAR^13^ lines including patients with early or late onset AD; Supplementary Table 1 and 2), with 16-72 lines per pool, to iMGL in naive (untreated), LPS, and IFN𝛄 treated conditions, to mimic inflammatory immune responses (<u>Methods</u>, Figure 1a)^14^. After quality control (<u>Methods</u>) we retained ∼2 million cells and integrated the pools using donors shared across all pools to remove batch effects (Supplementary Figure 1b). We analysed microglial subtypes present in our population using marker gene expression and label transfer^15^ (Figure 1b-d; Supplementary Figure 1b-d). This showed that untreated cells presented a signature similar to disease-associated microglia (DAM, expressing *CD9*, *LGALS3* and *PLA2G7*, 85% of cells), a small percentage of homeostatic microglia (HM, expressing *P2RY12* and *CX3CR1*, 2%) and a proliferative cluster (proliferating; expressing *MKI67*, *ASPM*, *UBE2C*, *RRM2* and *DLGAP5*^16^, 11%).

**Figure 1:**
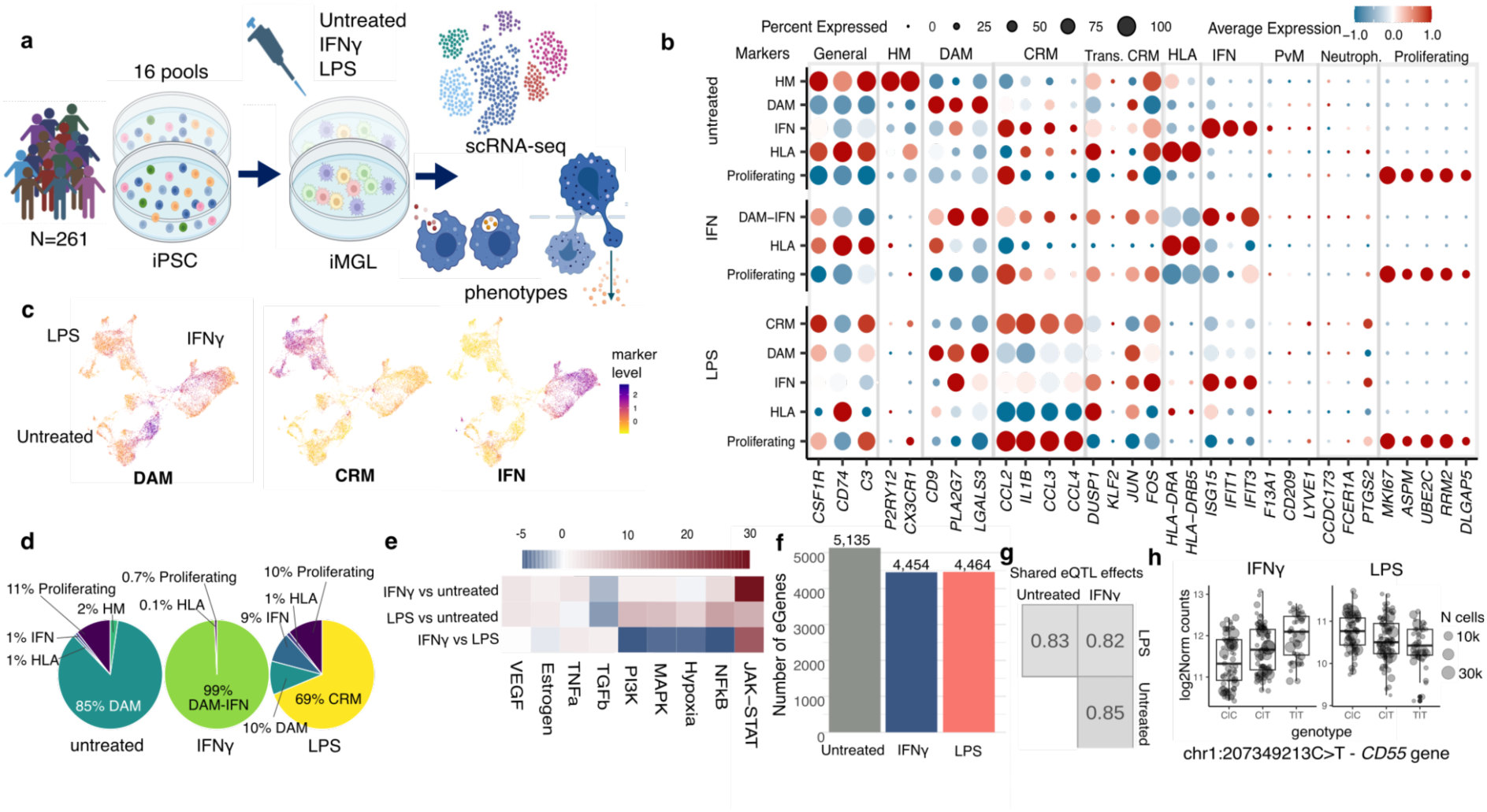
Transcriptomic profile of iPSC-derived microglia. a) Experimental summary. b) Markers of microglial subtypes expressed in the three treatments. c) Aggregated marker scores from *AddModuleScore* of DAM, CRM and IFN markers of microglial subtypes overlaid over UMAP of merged treatments. d) Abundance of each annotated cluster per treatment. e) Pathway activity analysis with PROGENy on the pairwise differential expression results. f) Number of significant eGenes. g) Proportion of pairwise significant shared eQTL effects. g) eQTL in an eGene with diverging effects in IFN𝛄 and LPS. DAM: Disease- associated microglia, CRM: Cytokine-response microglia, HM: Homeostatic microglia, IFN: interferon response microglia.

LPS treatment induced a signature of cytokine-response microglia (expressing *CCL2*, *CCL3*, *CCL4* and *IL1B*) in 69% of cells), with 10% of cells retaining a DAM signature, consistent with the microglial LPS response reported in the literature in mice and human^17^. Conversely, IFN𝛄-treated cells had a smaller proliferative cluster (1%), and overall showed increased expression of HLA genes and interferon response (IFN markers, including *IFIT1*, *IFIT3* and *ISG15*), as expected^18^. To maximize cell number, and thus power in downstream analyses, we focused only on proliferative and non-proliferative cell clusters.

Differential gene expression (DE) analysis (<u>Methods</u>) identified 1726-2053 significant (FDR-adjusted p-value < 0.05 & abs(log2FC)>1) DE genes per comparison (IFN𝛄 or LPS against the untreated baseline, and IFN𝛄 vs LPS, Supplementary Figure 2b, Supplementary Table 3). Gene set enrichment analysis (GSEA) of mSigDb Hallmark pathways (Supplementary Figure 2c) confirmed the increased expression of IFN𝛄 and inflammatory response genes in both the IFN𝛄 and LPS-treated samples. Pathway activity analysis (Figure 1e, <u>Methods</u>) highlighted a significantly increased JAK-STAT pathway activity in IFN𝛄, and to a lesser degree LPS, compared to untreated microglia. *FOXA2* was upregulated in the untreated samples compared to both LPS and IFN𝛄 (Supplementary Figure 2d), consistent with its protective role against inflammatory phenotypes in microglia^19^. This indicated that our iMGL recapitulated the inflammatory transcriptional response.

We next investigated how genetic variation influences gene expression in resting and stimulated iMGL. Following genotype and gene expression quality control (see <u>Methods</u>), we retained 6,405,518 variants and 188-189 lines for analysis. This resulted in 7,121 unique significant eGenes (FDR <5%; Figure 1f, Supplementary Table 4) and 12,230 associated variants across the three treatments, split fairly evenly across conditions.

To assess the consistency of genetic effects across conditions, we compared each significant eQTL-eGene pair across the three treatments (<u>Methods</u>). This revealed a high degree of pairwise eQTL sharing across treatments (83-85%; Figure 1g). When considering only significant eGenes (FDR < 0.05), 74.6% were shared across all three treatments (Supplementary Figure 3b, Supplementary Table 5). Most of these shared eGenes had similar effect sizes (differences in beta < 0.5), while only 0.3% of eGenes showed the same direction of effect but significantly different magnitude, and 0.3% exhibited opposite directions in at least two conditions such as the diverging effects between LPS and IFN𝛄 for *PARP1* and *CD55* (Figure 1h). Only 11.2% of eGenes were significantly shared by any two treatments, while 14.2% were treatment-specific. The treatment-specific eGenes highlighted regulation of genes in processes specifically induced by treatment responses, e.g. *STAT1* and *PPARG,* two genes in this category, are central in the enriched network of nitrogen metabolism in IFN𝛄-stimulation (STRINGdb FDR = 3 x 10^-5^, Supplementary Figure 3c-d), which regulates the production of the nitric oxide compounds and macrophage polarization in response to inflammatory signals^20^.

### A regression-based framework for donor identity estimation in pooled differentiation assays

While pooling iPSCs from multiple donors improves scalability, a major challenge lies in the variable response of individual donor lines to cell culture conditions. Some donors differentiate rapidly or proliferate efficiently, leading to imbalanced representation in the final cell pool. This variation can distort assay readouts and confound downstream association analyses if not properly accounted for.

To address this, it is critical to monitor donor composition during differentiation to accurately measure donor-specific effects. We developed a method, *poodleR* (POoled dOnor Deconvolution by LEast square Regression), to estimate donor proportions from low-depth whole genome sequencing (WGS). This approach leverages known donor genotypes and minor allele frequencies across sequenced variants in the pool, applying constrained least squares regression to deconvolute the contribution of each donor with high accuracy, even at low coverage (Figure 2a).

**Figure 2:**
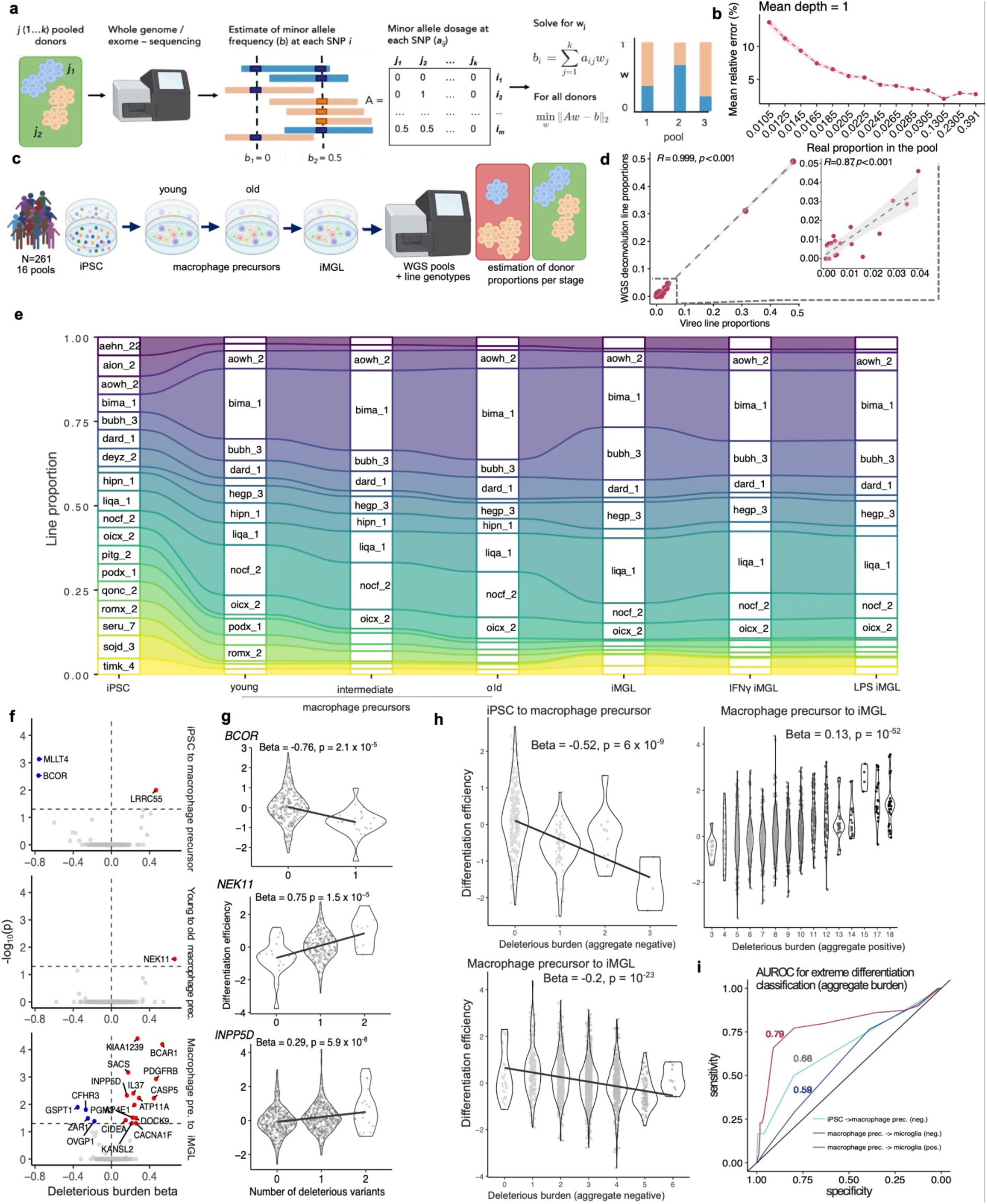
Deconvolution method and statistical analysis of differentiation efficiency associations. a) Schematic representation of the deconvolution method. b) Relative error at representative line proportions. c) Schematic representation of the microglia differentiation steps, including each step where line proportion was measured. d) Correlation between WGS deconvolution method and vireo estimates. e) Line proportions at every differentiation stage for a representative pool. f) Genes with significant (Bonferroni-adjusted p- value < 0.05) burden of deleterious variants on microglial differentiation efficiency for iPSC to macrophage precursor, macrophage precursor aging, and macrophage precursor to microglia stages. g) Violin plots of differentiation efficiency across deleterious variant burden for selected significant genes. h) Effect of aggregate burden within all significant genes (of same directionality) on differentiation efficiency for iPSC to macrophage precursor, and macrophage precursor to microglia. i) Area Under the Receiver Operating Characteristic curve (AUROC) for the three aggregate burden measures from significant results.

To benchmark the accuracy of *poodleR*, we tested it on *in silico* pools with equal donor contributions (ten donors per pool). Even at a low mean sequencing depth (0.25×), the absolute error remained below 0.05 (Supplementary Figure 4a). We then evaluated performance under more realistic conditions using *in silico* pools of 19 donors with unequal proportions ranging from 0.0005 to 0.39. In this setting, absolute errors ranged from 0.005 to 0.03, while relative error at higher donor proportions remained low (∼5%, Figure 2b, Supplementary Figure 4b; see <u>Methods</u>). As expected, relative error increased for donors with smaller proportions, reflecting the reduced number of reads overlapping informative variants.

Despite this, *poodleR* achieved comparable accuracy to existing methods, such as census-seq, with both showing ∼94% accuracy at donor proportions of 2% (Supplementary Figure 5B). We also observed some donor-specific differences in accuracy, with certain donors consistently showing slightly higher or lower estimation precision (Supplementary Figure 5C)^21,22^. This may be explained by the degree of genetic similarity between donors in a pool, which can affect the ability to resolve individual contributions.

By leveraging single-cell RNA sequencing (scRNA-seq) at the microglia stage, we were able to directly benchmark the accuracy of our WGS-based donor deconvolution method, *poodleR*, against established single-cell deconvolution approaches. Specifically, we used Vireo to assign cell line identities from scRNA-seq and then calculated global donor proportions for comparison. The results demonstrated the high accuracy of *poodleR*, with a Pearson’s correlation of *R* = 0.99 across all donor proportions in matched samples (Figure 2d). Even for donors contributing less than 5% to the pool, the correlations remained strong (*R* = 0.87), underscoring the robustness of the method across a wide range of donor abundances.

### Genetic effects underpin donor composition during pooled iMGL differentiation

The iMGL differentiation protocol (∼42 days) has key transitions at day 14 and day 28, marking commitment to the macrophage precursor and microglial stages. We used *poodleR* to monitor donor abundance dynamics at different differentiation stages in 16 pools. DNA was collected for WGS at the iPSC, macrophage precursor (at different days in culture, termed “young”, “intermediate” and “old”), and microglia stages, with the latter also being treated with LPS or IFN𝛄 (Figure 2c).

Despite all pools presenting approximately equal cell line proportions at the iPSC stage, line abundance varied substantially during iMGL differentiation (Figure 2e). Competition differed throughout the process, such that donor lines taking over during iPSC growth (Supplementary Figure 5b) did not correlate with those outcompeting at the later stages (Figure 2e). Competition in culture is common to all pooled iPSC-derived cell studies, reflecting the variation of proliferation, survival and differentiation capacity of the donor cell lines (referred to from now on as “differentiation efficiency”, see <u>Methods</u>). Several lines were shared across pools to control for batch effects: two lines (hegp_3 and aowh_2) were present in all pools, with several others repeated in a variable number of pools. We also included seven pairs of hiPSC clones from the same donor to measure clonal variability (Supplementary Table 1). We observed consistent changes in donor abundance for the same cell lines repeated across different pools (average interquartile ranges for repeated lines < 1, consistently smaller than that of a non-shared line; Supplementary Figure 5c), as well as across replicates of different iPSC clones from the same donor (data not shown). This indicates that genetic effects underpin differentiation efficiency.

We therefore tested the effects of genetic variants on changes in donor proportions across key differentiation transitions: iPSC to macrophage precursor, young to aged precursors, and precursor to microglia, but did not detect any genome-wide significant variants (Supplementary Fig. 6d). However, there were three signals at p-value < 10^-7^. Two variants in the differentiation from precursors to microglia in the IFN𝛄 condition, one at chr4:48871391AT>A within 20 kb of *OCIAD1* and *OCIAD2* (involved in the JAK-STAT and Notch pathway regulating cell cycle progression and differentiation^23^), and another at chr7:138680001G>A within an intron of *SVOPL*. The third signal chr4:135084336C>T was found in the differentiation from iPSC to macrophage precursors within the lncRNA ENSG00000248434.

Given the limited power to detect individual common variant effects we next evaluated the burden of deleterious exonic variants per gene. Using whole exome sequencing (WES), we tested all genes containing at least one deleterious (7,757 genes), missense non-deleterious (1,012 genes), or synonymous variants (9,080 genes) (Figure 2d; <u>Methods</u>). We identified three genes with significant deleterious variant burden during the transition from iPSC to macrophage precursor (Bonferroni corrected p = 2 × 10⁻⁴): *BCOR*, *MLLT4*, and *LRRC55* (beta = –0.76, p = 2 × 10⁻⁵; beta = –0.76, p = 6 × 10⁻⁶; and beta = 0.47, p = 8 × 10⁻⁵, respectively; Figure 2f, Supplementary Table 6). Notably, deleterious *BCOR* variants have previously been linked to decreased dopaminergic neuron differentiation and enhanced iPSC survival^3,24^. This association was also evident when considering missense non-deleterious variants (Supplementary Figure 7a). For the transition from young to aged macrophage precursors, we identified a single significant gene, *NEK11* (beta = 0.66, Figure 2g), which encodes a kinase involved in the G2/M DNA damage checkpoint and is upregulated in cells with an arrested cell cycle^25^. The precursor-to-iMGL transition yielded 18 gene associations (Figure 2f, Supplementary Table 6), most in the direction of enhancing microglial differentiation. One example is *INPP5D* (beta = 0.15, Figure 2g), an AD-associated gene that regulates the inflammasome in microglia, and is also associated with monocyte and eosinophil counts^8,26^.

We found that the cumulative burden of deleterious variants can be used to partially predict lines that may disproportionately expand or disappear in pooled cultures (Figure 2h–i; AUC = 0.59–0.79). This aggregate score was constructed by summing the burdens of deleterious variants across all genes that were significant in the burden tests and showed consistent directionality of effect for each transition. The global burden of deleterious variants across all genes and lines showed a modest yet significant overall effect on precursor to iMGL transition (beta = 0.0008, p = 8 × 10⁻⁵; Supplementary Figure 7b). Finally, aggregating the significant genes across all differentiation transitions revealed significant enrichment for essential genes (<u>Methods</u>) and genes required in macrophage survival^27^ (p = 0.02 and 0.04 for DepMap and Covarrubias *et al.*’s sets, respectively; Supplementary Figure 7c).

### Genetic mediators of phagocytosis and migration in iMGL

Microglia mediate key neurodegenerative processes by sensing environmental cues and clearing protein aggregates, dead cells and synapses. Two essential functions related to these are migration and phagocytosis. Therefore, identifying genetic variants and genes that regulate these functions informs mechanisms underlying cell phenotypes and disease.

We used a dual-fluorescent reporter assay to quantify phagocytic capacity, where iMGLs engulf GFP- and mCherry-labeled dead SH-SY5Y cells (pmChGIP SH-SY5Y). GFP fluorescence is quenched rapidly in the acidic endolysosomal compartment. We sorted two cell populations (mCherry⁺/⁻) by FACS and measured donor abundance by WGS (Figure 3a). This assay measures at the same time phagocytosis, acidification and degradation of reporter cells, which we will describe subsequently as the “phagocytosis” phenotype. Migration was assayed using transwells: microglia were placed on top, and DNA from cells migrating to the bottom or those remaining at the top was sequenced (Fig. 3a). The lower compartment contained chemoattractant (C5a) or media alone, to measure chemotaxis versus random migration.

**Figure 3:**
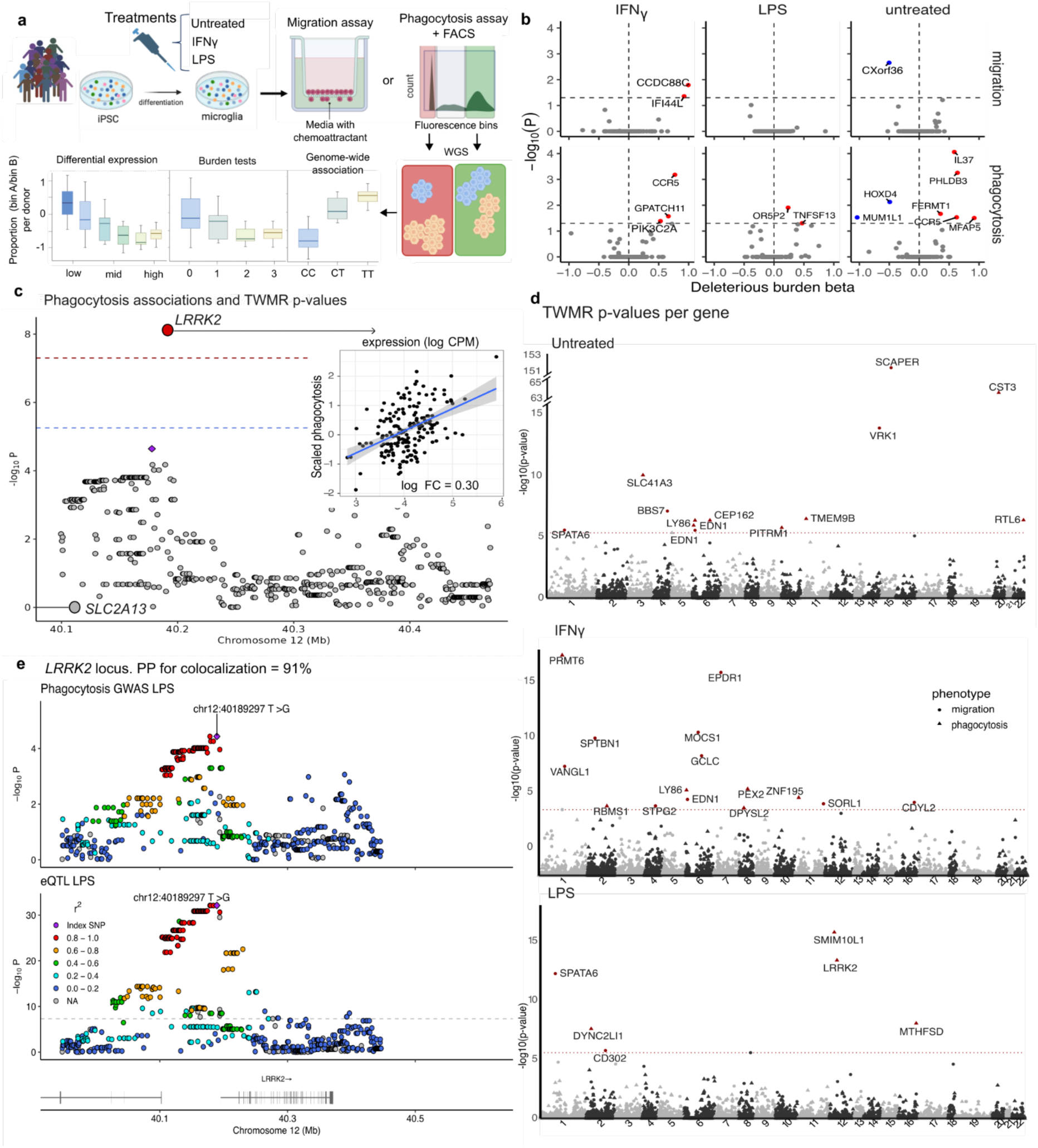
Genetic associations with cellular phenotype. a) Schematics of the assays and statistical testing strategy. b) Significant results for the burden tests of deleterious variants with phagocytosis and migration, per treatment. c) Comparison of the phagocytosis GWAS p-values at *LRRK2* and the p-value for the aggregated *LRRK2* gene signal in the eQTL-informed TWMR. Red line signals genome-wide significance for GWAS (5x10^-8^) and the blue line the significance for TWMR (10^-6^). Gene-annotated horizontal lines with larger dots indicate significance for TWMR. Inserted scatterplot shows phagocytosis phenotype (y axis) vs gene expression, with blue linear regression line. d) Miami plots of TWMR results (p-values per gene) integrating phagocytosis or migration GWAS and eQTL results, per treatment. Red lines signal genome- wide significance for TWMR (∼5x10^-5^, above). Only significant genes with F-statistic above 10 are highlighted. e) Colocalization plot for the phagocytosis GWAS and the eQTLs in the LPS-treated sample at *LRRK2* locus.

No genome-wide significant associations were detected for either phenotype, likely due to limited power from small sample size and phenotypic variability (Supplementary Figs. 9–10), consistent with previous reports on cellular trait GWAS in *in vitro* models^28,29^. However, when testing for gene burden of exonic deleterious variants we identified 11 genes (*IL37*, *PHLDB3*, *CCR5*, *HOXD4*, *OR5P2*, *FERMT1*, *GPATCH11*, *MUM1L1*, *MFAP5*, *PIK3C2A*, *TNFSF13*) associated with phagocytosis and three (*IFI44L*, *CCDC88C*, *CXorf36*) with migration (Figure 3b, Supplementary Fig. 11a, Supplementary Table 7, <u>Methods</u>). For example, *CCR5*, a chemokine receptor highly expressed by microglia, was linked to phagocytosis in IFNγ-treated and untreated iMGLs (Supplementary Fig. 11b), in line with the role of *CCR5* in mediating microglial recruitment and activation at sites of CNS injury or neurodegeneration, in synaptic plasticity, and upregulation in AD- associated microglia^30^. Additionally, *IL37* was associated with enhanced phagocytosis in untreated microglia (Fig. 3b), aligning with prior findings in monocytes and macrophages^31^.

Given that phenotype-associated variants are enriched for expression quantitative trait loci (eQTLs),^32,33,34^ we leveraged our microglia-specific eQTL map to perform a transcriptome-wide Mendelian Randomization (TWMR) analysis^35^ (<u>Methods</u>). Unlike variant-level tests, TWMR models joint causal effects of multiple eGenes at a locus, accounts for correlated regulatory architecture, mitigates horizontal pleiotropy, and is well-suited to underpowered GWAS. Applied to phagocytosis and migration across three treatment conditions, TWMR identified 32 significant gene–trait associations (Bonferroni p < 0.05 and F-statistic above 10; Fig. 3c–d, Supplementary Table 8). Among the 19 phagocytosis genes, we identified *DYNC2LI1* (LPS-treated), a component of the dynein motor complex which is essential for various intracellular transport processes^36^, and *PEX2* (IFNγ-treated) which is involved in peroxisome-mediated lipid and reactive oxygen species modulation, a process dysregulated in disease associated microglia^37,38^. Notably, *LRRK2* showed a strong causal link with phagocytosis in LPS-treated cells (α = 0.67, p = 4×10⁻^14^; Fig. 3c) consistent with the role of the G2019S activating variant on phagocytic activity in iMGL^39^. This association, driven by the lead *LRRK2* eQTL, is markedly stronger than conventional association testing (Figure 3d), which does not leverage gene expression information. We further validated this effect using colocalization (H4=0.91, Figure 3e, <u>Methods</u>). *LRRK2* is implicated in the pathobiology of PD and AD, and these results further establish a role of common genetic variants in regulating the expression of *LRRK2* and phagocytic activity in iMGL.

Amongst the 16 migration-associated genes we observed *VRK1, VANGL1, SPTBN1, EDN1, EPDR1, BBS7,* and *GCLC*, genes involved in cell polarity and motility.^40–48^ Importantly, we also uncovered disease-relevant genes, including *MTHFSD* (phagocytosis) and *SORL1* (migration), implicated in Amyotrophic Lateral Sclerosis (ALS) and AD GWAS, respectively. *SORL1* has been recently implicated in lysosomal regulation, a process closely intertwined with chemotaxis^49^. We also identified a phagocytic role for *CST3*, a cysteine protease that regulates amyloid-beta deposition and is linked to AD risk.^50,51^ We further identified microglial immune regulators such as *PRMT6, EDN1*, and *LY86*, shown to promote proinflammatory phenotypes in microglia and macrophages.^52,53,54,55^ Together, this approach, combining GWAS of cellular phenotypes with eQTLs, reveals new biologically and clinically relevant gene-phenotype relationships.

### iMGL transcriptomic and phenotypic regulation reveals mechanisms of neurodegenerative risk

Building on transcriptomic profiling and genetic analyses of migration and phagocytosis in iMGLs, we examined how these regulatory mechanisms intersect with neurodegenerative disease risk variants. First, using differentially expressed genes between resting and stimulated states revealed significant enrichment of AD^7^ and PD candidate genes across all treatments (GSEA, Benjamini-Hochberg [BH]-adjusted p < 0.05; Fig. 4a), indicating that IFNγ- and LPS-activated iMGLs recapitulate disease-relevant transcriptional signatures.

**Figure 4:**
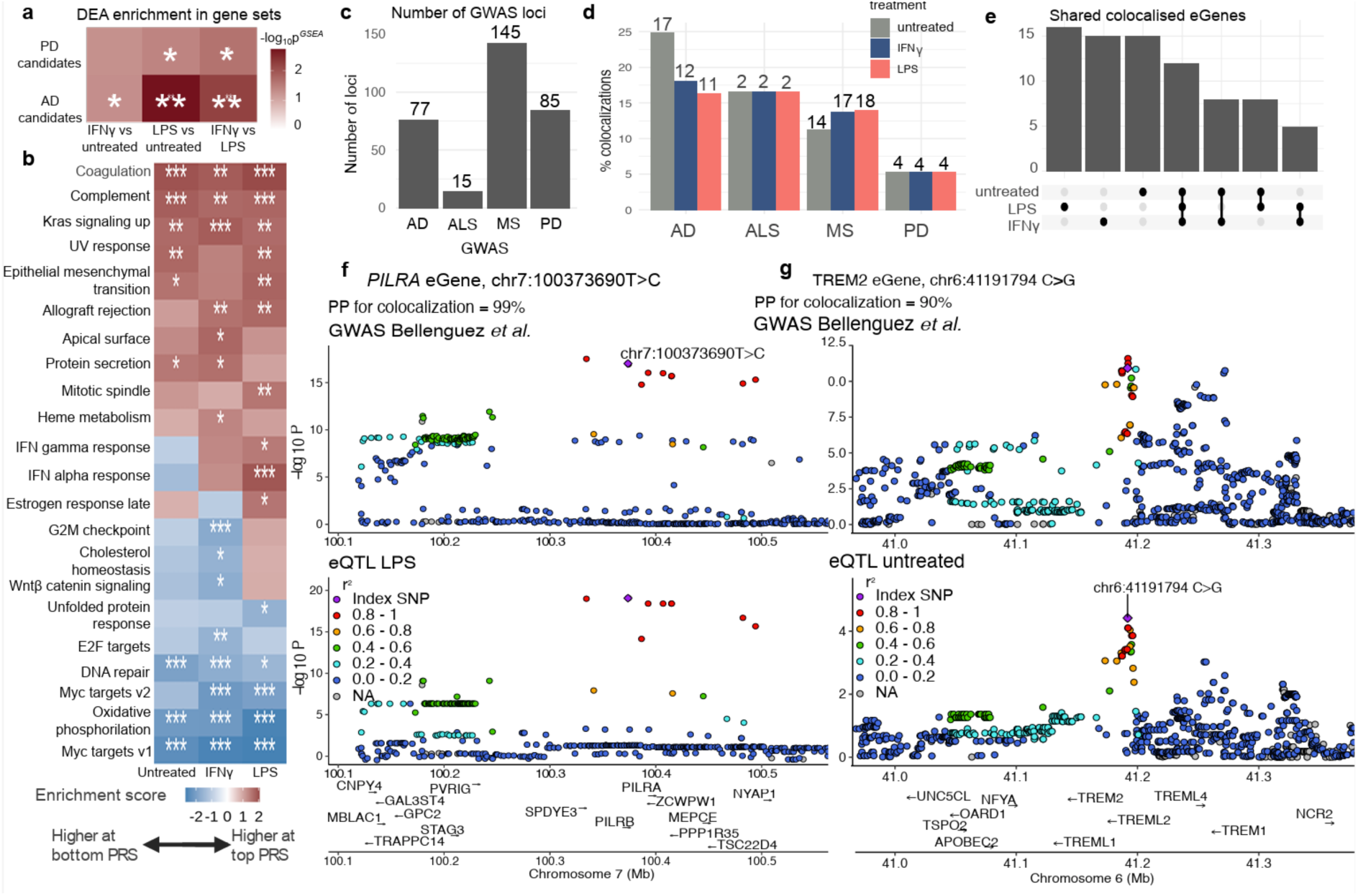
Linking iMGL biology to neurodegenerative disease. a) Enrichment of DEGs across treatments in AD and PD candidate genes. Color denotes adjusted p-value of enrichment using GSEA. Stars denote adjusted significance of * < 0.05, **<0.01,***<0.001 b) Enrichment of DEGs across AD-PRS in hallmark pathways within each treatment. Color denotes normalised enrichment score from GSEA. Stars denote adjusted significance of * < 0.05, **<0.01,***<0.001 c) Number of independent GWAS loci tested for colocalization. d) Number and percentage of colocalizations per GWAS dataset. e) Degree of sharing of colocalized eGenes across treatments. f) Colocalization plot of *PILRA* eGene with AD GWAS in the LPS- treated iMGL. g) Colocalization plot of *TREM2* eGene with AD GWAS in the untreated iMGL. PP = posterior probability.

Next, we explored if we could capture transcriptional changes induced by the cumulative effects of disease risk alleles using polygenic risk scores (PRS). We focused on AD, given its evidence for microglia involvement in disease pathology^7^. We computed PRS for each donor (<u>Methods</u>, Supplementary Fig. 12), including the 14 IPMAR donors with extreme PRS, to assess gene expression differences across treatments. PRS-based differential expression analysis (PRS- DEA, <u>Methods</u>) identified 143 significant genes in untreated, 56 in IFNγ-, and 21 in LPS-treated iMGLs (FDR < 0.05; Supplementary Fig. 13, Supplementary Table 9). Though modest in number, PRS effects were directionally consistent across conditions (Pearson’s R = 0.68–0.69; Supplementary Fig. 13d) and >75% remained significant after permutation testing (Supplementary Fig. 13e).

We observed that genes identified through PRS-DEA converged on specific pathways (Supplementary Fig. 14a; Fig. 4b; <u>Methods</u>). High AD PRS was associated with reduced expression of oxidative phosphorylation genes across all conditions, consistent with AD-linked mitochondrial dysfunction^56^. Inflammatory response pathways were upregulated in both IFNγ- and LPS-treated microglia with high PRS, with a stronger signal in the LPS condition, including significant activation of *STAT1* targets, driven by increased expression of *SPI1, STAT1,* and *RFX* family targets. These results support a model in which increased AD risk involves heightened microglial inflammatory responses^57^.

Furthermore, genes upregulated at high PRS within LPS were significantly enriched for AD heritability using partitioned LDSC (Supplementary Figure 14b, Methods).

We next assessed colocalization between iMGL eQTL and GWAS loci from AD, PD, ALS, and multiple sclerosis (MS) (Figure 4c). Among 77 AD independent loci from Bellenguez *et al.* (2022) we found 27 unique colocalizing eGenes at 22 loci (posterior probability of colocalization > 70%, Supplementary Table 10): *ALKBH5*, *ASPHD1*, *BIN1*, *CASS4*, *CIAO2A*, *CTSH*, *DOC2A*, *FAM131B*, *GPR141, GSTK1*, *IDUA*, *KAT8*, *LLGL1*,, *MS4A6A*, *NSF*, *PILRA, PILRB, PLEKHA1*, *PTK2B, RABEP1*, *RASA1, SORL1*, *TREM2*, *TRIM37, TRPM7, YPEL3*, and *ZYX*. From Nalls *et al.* (2019) PD GWAS (85 loci) we found seven unique colocalizing eGenes (*LRRK2, TMEM163, NUP42, NSF, SPNS1, GPNMB,* and *STX4*) at six loci. Finally, for MS (Patsopoulos *et al.* (2019), 145 signals) there were 41 unique eGene colocalizations at 29 loci and for ALS (van Rheenen *et al.* (2021), 15 signals) four unique eGene colocalizing at three loci (*C9orf72*, *SCFD1*, *MOB3B*, and *ERGIC1*). There were 79 unique eGenes colocalizing within any of the GWAS loci (Figure 4d-e, Supplementary Table 10- 11). AD had the highest colocalization rate (29%), followed by MS and ALS (20%); PD showed ∼4-fold fewer colocalizations (7%) despite more loci being tested (85 PD vs 77 AD loci). The larger number of colocalizations for AD highlights shared causal mechanisms underpinning AD and gene expression regulation in iMGL.

Using Open Targets^58^, we assessed how many of the 79 colocalizing eGenes were previously prioritized as likely causal. We found that 37 (47%) had low prioritization scores (L2G < 0.2), including 15 of 27 AD colocalizations (56%), with limited additional genetic evidence from the literature (Supplementary table 11). This suggests that previous AD gene prioritization efforts may have missed key microglia-specific regulatory contexts. We observed 16 of the 79 colocalized genes were unique to the LPS condition, while 15 were unique to IFNγ (Figure 4e), thus identifying loci with complex, context specific gene regulatory effects, manifesting only upon a specific stimulation. For example, the AD-associated locus 16:30010081, colocalized with a different eGene depending on the condition: *DOC2A* in the untreated state, *ASPHD1* in IFNγ-treated cells, and *YPEL3* in LPS-treated cells. Our results recapitulated eight out of 14 AD colocalizations, three out of four PD colocalizations, and the single ALS colocalization (*SCFD1* eGene) previously reported in primary microglia^59,60^ (Supplementary Table 10), highlighting the similarity between iMGL and primary cells and reinforcing iMGLs as a scalable, disease-relevant model of brain immune function.

Our data enabled nomination of new candidate causal genes at disease loci in stimulated iMGL, e.g. at an AD locus, variants in high LD colocalized with expression of both *PILRA* and *PILRB* (PPH4 > 0.70; Fig. 4f, Supplementary Figure 15). These genes encode transmembrane receptors with distinct roles in inflammatory signaling and phagocytosis. The common AD-protective *PILRA* chr7:100406823 C-allele is in LD with *PILRA* G78R coding variant, which reduces ligand binding, including decreased entry of HSV-1^61^, supporting the hypothesis that recurrent infections may increase AD risk. *PILRA* KO microglia present altered immune signaling, including reduced cytokine production and increased chemotaxis^62^. *PILRB,* in contrast, encodes an activating receptor that signals through TYROBP/DAP12, promoting inflammatory responses, chemotaxis towards amyloid beta and phagocytosis, and increased neuronal damage^63,64^. While *PILRA* is often cited as the causal gene at this locus based on fine-mapping, rare coding variants^61^, and previously reported colocalization, our data provide colocalization evidence for both *PILRA* and *PILRB* (Supplementary Figure 15). In IFNγ-treated cells, the lead eQTL variant for *PILRB* (chr7:100334426C>T, rs7384878, beta = 0.58) is also the lead AD GWAS variant (beta = 0.92, PPH4=0.99). This variant is in high LD (r^2^=0.85) with the lead *PILRA* eQTL variant chr7:100482234A>T (rs6971558, beta = 0.22) in the same condition. In LPS-treated microglia, another eQTL variant (chr7:100373690T>C, rs2405442), also in strong LD with the GWAS lead variant (r^2^=0.97), serves as the lead eQTL variant for *PILRA* (beta = 0.27) and *PILRB* (beta = 0.99), further supporting the involvement of both genes. To our knowledge, this is the first evidence of colocalization for both *PILRA* and *PILRB* in microglia.

In untreated microglia we identified *TREM2,* which encodes a receptor for amyloid beta and apolipoprotein particles, and upon ligand binding, activates microglia via the TYROBP/DAP12 adaptor, similar to PILRB, triggering migration, cytokine production, and phagocytosis. Rare missense mutations in *TREM2* have been implicated in AD and neurodegenerative disorders^65,66^. We found a colocalization (PPH4 = 0.90, Figure 4g) driven by a common eQTL variant, where the disease protective allele increases the gene expression (chr6:41191794 C>G, rs3800342; FDR=0.02, beta = 0.12; AD beta = -0.059; MAF = 0.36). This variant lies in the 3’ of *TREM2* with putative regulatory activity. Fine mapping of the *TREM2* locus in the Bellenguez *et al*. AD GWAS identified three independent rare risk variants, chr6:41181270 A>G (rs60755019, MAF =0.004, OR = 1.55), chr6:41161514 C>T (rs75932628, R47H, MAF = 0.003, OR = 2.39) and chr6:41161469 C>T (rs143332484, R62H, MAF = 0.013, OR = 1.41). The latter two are on the same haplotype (D′ = 1) as our common lead eQTL variant (Supplementary Figure 16a–c). This pattern is consistent such that the rare non- risk alleles are always on the AD-protective eQTL allele background, suggesting that increased *TREM2* expression may reduce AD risk, providing a mechanistic link between common regulatory variation and rare coding risk alleles.

Of the 79 colocalizing eGenes, *SORL1* (AD) and *LRRK2* (PD). were also identified in our TWMR analysis, providing a link from GWAS variant association, through eQTL to cellular phenotype. *LRRK2* is an established PD gene, and our TWMR results indicate its role in regulating phagocytosis.

In LPS-treated iMGLs, the lead colocalizing variant between phagocytosis and *LRRK2* expression (chr12:40189297 T>G; phagocytosis LPS beta = –0.387, MAF = 0.44, Figure 3e) is in perfect LD (D′ = 1, r^2^=1) with the lead *LRRK2* eQTL in the same condition (chr12:40178345 T>C, eQTL in LPS beta = –0.59, MAF=0.44). In contrast, the lead *LRRK2* eQTL in IFNγ-treated cells that colocalizes with PD-associated variants (chr12:40220632 C>T; eQTL-IFNγ beta = 0.39; PD GWAS beta = 0.14, MAF=0.13, Supplementary Figure 17) is only in moderate LD with the aforementioned phagocytosis-associated variant (chr12:40189297 T>G, D′ = 0.87, r^2^=0.15, MAF = 0.44). Despite this, the former variant is also significantly associated with changes in gene expression in LPS despite not being the lead signal in the region (nominal p-value = 6x10^-13^, eQTL-LPS beta = 0.86), following the same direction of increased gene expression associating with increased phagocytosis.

These observations suggest that *LRRK2* expression and its phenotypic consequences are modulated by treatment-specific regulatory variants. The more common haplotype, tagged by the T alleles at chr12:40189297 and chr12:40178345, respectively, is associated with increased *LRRK2* expression and enhanced phagocytosis in LPS-treated cells. However, this haplotype most often also carries the major C allele at chr12:40220632, which reduces *LRRK2* expression in IFNγ-treated cells and is associated with reduced PD risk. Because the lead variants in LPS are relatively common (MAF = 0.44), LD estimates with the rarer IFNγ-specific variant may be less precise. Together, these findings underscore the complexity of regulatory architecture at the *LRRK2* locus and highlight the importance of accounting for context-specific gene regulation when interpreting disease associations. Further functional validation will be needed to fully disentangle the molecular mechanisms linking *LRRK2* to microglial function and PD risk.

Finally, to explore how disease-linked genes might influence microglial behavior, we correlated gene expression with phagocytosis and migration phenotypes for genes within AD and PD GWAS loci (<u>Methods</u>). Expression of 47 AD- and 19 PD- candidate genes from GWAS associate significantly with phagocytosis (FDR < 0.05), and 49 AD- and 18 PD-associated genes with migration (Supplementary Table 12). Among these, *LRRK2* expression increases with phagocytic activity (log₂FC = 0.30), consistent with the direction of its eQTL effect, and also with migration (log₂FC = 0.51). Conversely, *TREM2* expression is negatively associated with both phenotypes (log₂FC = –0.11 and –0.28, respectively; Supplementary Fig. 18a). Looking at global expression patterns, we observed an inverse relationship between several regulatory pathways and TFs compared to those seen along the PRS differential expression axis (Supplementary Fig. 18b-c, Figure 4b), including increased oxidative phosphorylation gene expression associated with higher phagocytic activity. Additionally, to assess the collective phenotypic impact of disease-associated eQTLs, we constructed microglia-specific PRS from regions with strong eQTL-AD GWAS colocalization (PPH4 > 70%), computed per treatment (<u>Methods</u>). We found a significant inverse association between the polygenic component of the AD eQTL-PRS and phagocytosis in LPS-treated microglia (p = 0.04, beta = –0.14; Supplementary Fig. 19), suggesting that increased genetic risk is associated with reduced phagocytic activity under inflammatory conditions.

These findings underscore the utility of pooled iMGL assays in uncovering disease-relevant regulatory mechanisms and suggest that neurodegenerative risk genes modulate microglial gene expression and function. To build on this, we performed targeted CRISPR knockouts of genes implicated in Alzheimer’s disease.

### Targeted KO CRISPR screening identifies modulators of phagocytic uptake of dead neurons in iMGL

To better understand the causal link between AD-associated GWAS genes and microglia function we performed a pooled CRISPR KO screen for phagocytosis in iMGL. Cells were transduced with a dual-guide CRISPR library (NeuroKO) targeting 251 genes consisting of the top 2-3 candidate genes at each AD GWAS locus and controls (Supplementary Table 13). Following 14 days of differentiation, iMGL were incubated with dead/fixed pmChGIP SH-SY5Y to assess phagocytosis. iMGL were then fixed and sorted into four bins based on mCherry intensity, reflecting different levels of phagocytosis, rates of degradation or export^14^ (Figure 6a and 6b).

**Figure 6:**
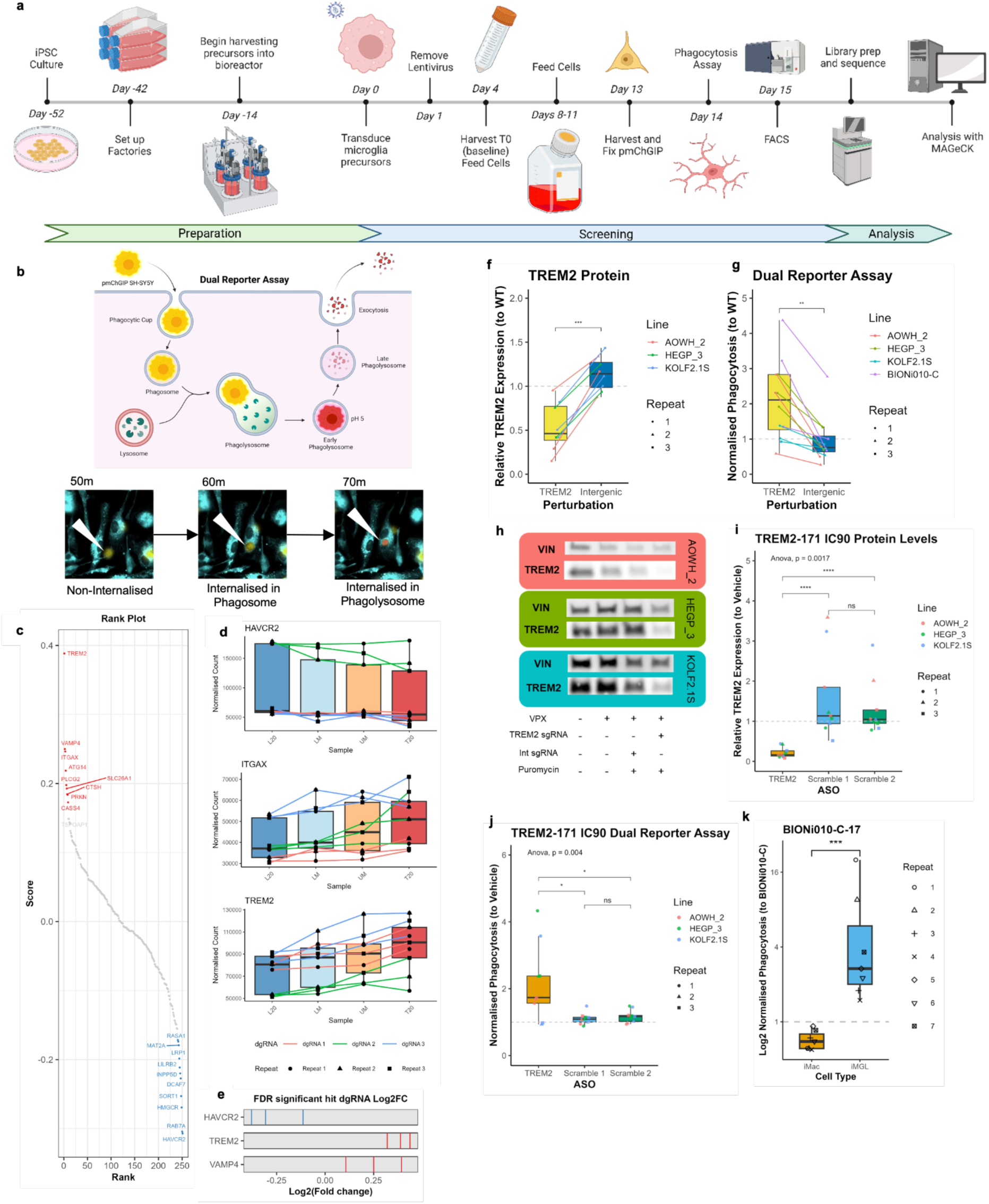
KO Screening in iMGL to identify regulators of phagocytosis of dead neurons. a) Outline of the screening protocol, a full method available at Washer *et al.,* (2025)^70^ b) Representative images of iMGL (turquoise) phagocytosing the dual reporter cargo (yellow), only once the dual reporter SH-SY5Y is internalised to the phagolysosome does the signal change from yellow to single red within 10 minutes of internalisation c) Rank plot of the genetic targets where KO results in increased phagocytosis (red), or decreased phagocytosis (blue). d) Boxplots showing the distribution of the FDR significant hits (*HAVCR2, TREM2, ITGAX*) dgRNA across the four sorted populations in the three repeated screens confirm enrichment. e) The log2FC of the three dgRNA in the three FDR significant hits between the highest and lowest phagocytosing populations, aggregated from the three screens using MAGeCK. f) *TREM2* KO was validated using arrayed CRISPR screening in four different genetic backgrounds with three replicates. *TREM2* expression was significantly reduced and g) phagocytosis was significantly increased in iMGL transduced with *TREM2* sgRNA and not Intergenic sgRNA. Asterisks indicate p-values from Wilcoxon test (* p<0.05, ** p<0.01, *** p<0.001). h) Representative western blots for WT iMGL, VPX only transduced iMGL, *TREM2* sgRNA transduced iMGL, Intergenic sgRNA transduced iMGL. The *TREM2* and Intergenic were selected with puromycin to reduce the background. i) *TREM2* targeting ASO resulted in significant KD of *TREM2* at IC90 and j) a significant increase in phagocytosis in iMGL, compared to cells treated with matched scramble controls. Asterisks indicate p-values from Anova (* p<0.05, ** p<0.01, *** p<0.001). k) The *TREM2* KO line BIONi010-C-17 was differentiated to iMGL or macrophages (iMac) and showed opposite effects on phagocytosis in the different cell types. p-values from Wilcoxon test, *** p<0.001.

We identified three genes at FDR p<0.05 with *HAVCR2* KO decreasing phagocytosis, and *ITGAX* and *TREM2* KO increasing phagocytic signal. Additionally, we observed a suggestive signal for several biologically relevant genes reducing phagocytosis (*RAB7A, HMGCR, SORT1, INPP5D* and *LRP1;* Figure 6c), and increasing phagocytosis (*VAMP4, ATG14, PLCG2, CTSH,* and *PRKN*; Figure 6c). Examining the normalised dgRNA counts across the four sorted bins showed clear directional trends for enrichment, indicating this is not a sampling bias (Figure 6d/e). We further examined the correlation of genes shared between the NeuroKO screen and an independent targeted screen using a single sgRNA approach with 3x sgRNAs per gene and these showed a strong correlation and direction of effect (Pearson’s R=0.86, p=9.1e^-5^), despite being different guides, vectors, and independent repeats (Supplementary figure 20), highlighting detection of robust phenotypic effects.

Interestingly, the identification of *TREM2* KO resulting in an increased mCherry signal (log2FC = 0.389, FDR p-value 0.00125) was consistent in direction with the eQTL-phagocytosis effect. This result was somewhat unexpected given previous reports that *TREM2* knockout reduces phagocytosis in macrophages and microglia^67–69^. To confirm the validity of our finding, we evaluated its consistency across multiple orthogonal approaches, including arrayed CRISPR, antisense oligonucleotide treatment (ASO), and iPSC knockout lines.

### Divergent phagocytic activity in *TREM2* knockout in iMGL and iMacrophage lineages

To confirm the *TREM2* finding we undertook an arrayed CRISPR approach utilising four different iPSC genetic backgrounds^71–74^. We transduced them with lentiviral pools of three sgRNA all-in-one lentivectors targeting either *TREM2* or intergenic regions (as cutting controls). We harvested protein for TREM2 western blotting and performed the phagocytosis assay as previously described. We identified approximately 50% knockout of TREM2 at the protein level in cells treated with *TREM2* targeting guides compared to those targeted with intergenic controls (Wilcoxon, p-value <0.01, N=9 (3 lines, 3 repeats); Figure 6f, h). We identified an increase in the proportion of mCherry positive iMGL in cells treated with *TREM2* targeting sgRNA across all four genetic backgrounds, confirming that *TREM2* knockout increases phagocytosis in iMGL (Wilcoxon p-value=0.0029, N=12 (4 lines, 3 repeats); Figure 6g).

To confirm that the observed phenotype was not a result of lentiviral KO of *TREM2,* we used *TREM2-*targeting ASO^73^. At IC90 we observed a strong decrease in TREM2 protein expression with *TREM2* targeting ASO but not in the scrambled controls (ANOVA, p-value=0.0017) (Figure 6i) and a significant increase in phagocytosis (ANOVA, p-value=0.004) (Figure 6j). A second ASO targeting *TREM2* showed the same phenotype at IC90 (Supplementary Figure 21).

We further validated this finding using the clonal *TREM2* KO iPSC line (BIONi010- C-17) and isogenic control (BIONi010-C) differentiated to both iMGL and macrophage-like cells (iMacs) in parallel. When differentiated to iMGL, there was an increase in phagocytic signal with *TREM2* KO, confirming our results from lentiviral or ASO treatments. Interestingly, when the same precursors were differentiated into iMacs there was reduced phagocytosis with *TREM2* KO (Wilcoxon p-value=0.00058, N=7 (Figure 6k)), consistent with previous literature^67–69^. These results demonstrate that modulation of *TREM2* leads to different cellular phenotype outcomes, depending on the cell state and cell type.

Taken together, our results demonstrate that the impact of *TREM2* modulation on phagocytosis is highly cell state–dependent, with robust knockdown and knockout increasing phagocytosis in iMGL but reducing it in iMacs, and that only strong reductions in *TREM2* expression are sufficient to elicit a phenotypic response. This highlights the importance of cellular context in interpreting the role of *TREM2* in neurodegenerative disease, where microglia are thought to mediate key aspects of pathology, including impaired debris clearance and chronic inflammation.

## Discussion

Establishing causal links between genetic variants identified by GWAS, their effector genes and the cellular mechanisms they perturb remains a central challenge in complex traits. For neurodegenerative diseases, an additional challenge resides in inaccessibility of the primary tissue, such as microglia. Context-specificity of gene regulation further complicates interpretation, as non-coding regulatory variant effects often manifest only under defined cellular states, stimuli, or developmental stages. Our study addresses these challenges and further emphasizes the critical role of context-specific effects, apparent in our eQTL and CRISPR results.

We demonstrate that pooled iPSC-derived iMGL provide a powerful and scalable model for dissecting genotype–phenotype relationships. Despite inherent variability in differentiation efficiencies, we demonstrate we can control for such effects using a careful experimental design including shared and repeat donors, and pooling of genetically matched donors. Combined with our low-depth WGS-based statistical method for donor deconvolution (poodleR), and quantitative modeling of donor effects we can robustly infer genetic associations across molecular and cellular traits. We benchmarked the performance of poodleR against single-cell inferred donor deconvolution and demonstrated its robust performance, contributing a new modelling tool to this expanding community.

In line with previous studies of similar sample size, we were underpowered to detect significant effects of common variants on cellular phenotypes (differentiation efficiency, phagocytosis, and migration). Nevertheless, we detected effects at a suggestive p-value < 10^-7^, indicating that these variants could contribute to shaping cellular phenotypes, but larger sample sizes are necessary to reach sufficient statistical power. However, stronger effects driven by rare deleterious variant analyses identified multiple gene burden hits associated with differentiation, phagocytosis, and migration. We identified 22 significant genes linked to cell expansion at various stages of differentiation, and observed that the cumulative burden of deleterious variants in these genes can partially predict disproportionate donor expansion. This highlights the potential of using it as a pre-screening tool to identify pools of donor lines with matched differentiation outcomes, which will improve the efficiency of this approach in future. Furthermore, we identified enrichment of deleterious variants in 14 genes associated with phagocytosis and with migration, capturing established phenotypic gene effects as well as highlighting novel genes in relevant biological processes. These results demonstrate that *in vitro* pooled iPSC models are effective for identifying genetic regulators of cellular phenotypes. As these approaches scale to larger numbers of donors, they will gain power to resolve both rare, high-impact loss-of-function variant effects, and more subtle contributions from common variants, enabling systematic dissection of the genetic architecture underlying cell biology.

Integrating transcriptomics into our approach added a critical molecular layer that enhanced variant interpretation at multiple levels, linking genetic variation to gene expression, identifying mediators of cellular phenotypes, and strengthening connections to neurodegenerative disease associations. First, differential gene expression analysis across the spectrum of AD polygenic risk and different conditions (resting, LPS and IFNγ) revealed that high PRS is consistently associated with oxidative phosphorylation gene expression across conditions, aligning with established links between AD and mitochondrial dysfunction^56^. In parallel, inflammatory response pathways were upregulated in both IFNγ- and LPS-treated microglia with high PRS. These findings support a model in which elevated genetic risk for AD drives exaggerated microglial inflammatory responses^57^ and suggest that disease-associated alleles converge on gene programs that impair key cellular functions, emphasising the need of further functional phenotyping to elucidate the mechanisms linking genetic risk to microglial dysfunction.

Second, we leveraged transcriptomic data to identify genes that mediate microglial cell phenotypes (phagocytosis and migration), and to inform the interpretation of disease associations. By incorporating transcriptome-wide Mendelian randomization (TWMR) and our matched eQTL results into our analysis of cell phenotype GWAS, we identified 34 significant gene–trait associations linked to phagocytosis or migration. These analyses revealed several biologically relevant genes, including known AD-associated loci such as *LRRK2*, which we now connect to phagocytosis, and *SORL1*, which we link to migration, providing functional context to genetic risk loci through direct effects on microglial behaviour.

Third, our study provided insights into the regulation of gene expression in both resting and stimulated microglial states. We identified thousands of eGenes, with approximately ∼75% shared across all treatment conditions. These eGenes showed significant colocalizations with neurodegenerative disease risk loci. For AD, 29% of tested loci showed evidence of colocalization, while colocalization rates were lower for ALS, MS, and particularly PD. Notably, *LRRK2* and *TREM2*, canonical PD and AD risk genes, respectively, showed condition-specific colocalization patterns, underscoring the context-dependent architecture of gene regulation at these loci. In *LRRK2*, we observed allelic heterogeneity across conditions, with distinct regulatory variants contributing to phagocytosis and disease risk. Furthermore, nearly half of the colocalizations identified here were not previously reported in the Open Targets *locus2gene* analysis, reinforcing the importance of using both cell-type– and cell-state–relevant models to detect regulatory variants mediating disease risk. These findings align with previous work showing that genes with dynamic, temporal regulation are more likely to colocalize with GWAS loci than those with static, baseline eQTLs^75^.

To functionally validate the role of candidate genes in regulating microglial phenotypes, we performed pooled CRISPR knockout screens to quantify phagocytosis and autophagic flux across putative AD-relevant genes identified by GWAS. Several known regulators emerged as significant hits (*TREM2*, *PLCG2*, *VAMP4*), alongside previously uncharacterized candidates such as *HAVCR2*, a gene within a recently identified AD GWAS locus. Notably, *TREM2* knockout consistently increased phagocytic signal in iMicroglia, in contrast to prior reports in macrophage or microglial cells. *TREM2* is a well-established phagocytic receptor, yet studies have reported conflicting effects upon its downregulation. A review by Jay *et al.*^66^ suggested that *TREM2*’s role in phagocytosis is AD-stage dependent, where reduced expression impairs phagocytosis in early pathology but may enhance it in later stages. In advanced AD, microglia often adopt a disease-associated microglia (DAM) transcriptional state and accumulate around amyloid plaques. *TREM2* knockout in this context has been shown to decrease DAM marker expression and promote a return to a more homeostatic phenotype^15^. Our transcriptomic profiling revealed that untreated iMicroglia in our system closely resemble DAM-like states, which may explain the observed increase in phagocytic activity following *TREM2* knockout and the colocalization of TREM2 eQTLs with AD risk loci. Conversely, in iPSC-derived macrophages (iMacs), *TREM2* knockout showed a trend toward reduced phagocytosis, consistent with previous studies. These findings suggest that basal cell state profoundly influences the phenotypic consequences of gene perturbation and must be carefully considered in functional genomic studies of disease-relevant traits.

In summary, this multifaceted approach enabled us to connect genetic variants, both common and rare, to effector genes and gene programs, downstream cellular phenotypes, and potential disease mechanisms. Transcriptomic and phenotypic analyses in iMGL offer a powerful framework for interrogating microglial dysregulation in neurodegenerative disease. The stimulation-specific eQTLs and colocalizations we identified highlight the importance of immune context: placing cells under relevant inflammatory conditions revealed regulatory effects that would otherwise remain undetected, providing insight into how genetically driven disease risk is mediated through dynamic gene regulation. More broadly, our study demonstrates the feasibility and value of integrating genetic association data, single-cell transcriptomics, and pooled functional genomics to investigate genotype–phenotype relationships at scale. While we focused on iPSC-derived microglia, the experimental and computational framework is readily applicable to other iPSC-derived or primary cell types across a range of diseases. This includes applications in inflammatory, metabolic, cardiovascular, and psychiatric disorders, where context-specific regulation and cellular function are critical to pathogenesis. Future efforts will benefit from increased donor diversity, expanded perturbation conditions, and broader phenotype panels. Together, these approaches will be essential for systematically resolving the molecular mechanisms underlying complex traits and for translating genetic risk into mechanistic insight across diverse cellular systems.

## Methods

### Human iPSC maintenance

261 human iPSCs lines from 254 European donors were obtained from the HipSci project (http://www.hipsci.org, Cambridgeshire 1 NRES REC Reference 09/H0304/77, HMDMC 14/013, for a list of lines used, see Supplementary Table 1), IPMAR project^13^ or commercially from Bioneer: TREM2 knockout (BIONi010- C-17) and isogenic control (BIONi010-C). iPSCs were thawed onto tissue culture treated 6-well plates (Corning, 3516), coated with 10 μg/mL vitronectin (VTN-N) (Life Technologies, A14700) or 10 μg/ml Vitronectin-XF (StemCell Technologies, 07180) using complete Essential 8 (E8) medium (Thermo Fisher, A1517001) supplemented with 10% CloneR™ (StemCell Technologies, 05888). After thawing, cells were expanded in E8 medium for at least 2 passages using Gentle Cell Dissociation Reagent (StemCell Technologies, 100-0485) for cell dissociation. Multiple lines were synchronized by adjusting the splitting ratio of each line, aiming for 60-85% of confluence at each passage and on the pooling day.

To enable pooling of more lines, mini-pools of iPSCs lines were generated, cryopreserved and subsequently thawed for final pooling. Briefly, iPSC cultures were dissociated into single cell suspension with Accutase (Millipore, SCR005) and resuspended in E8 medium containing 10 μM Rock inhibitor Y-27632 (StemCell Technologies, 72305). Cell counting was performed using Countess II (Thermo Fisher) and equal amounts of each iPSC line were pooled in E8 medium containing 10 μM Rock inhibitor Y-27632. The cell suspension was seeded at 40,000 cells/cm^2^ on vitronectin and cryopreserved after 3 days in Cell Freezing Medium consisting of 90% Knockout Serum Replacement (Life Technologies, 10828-028) and 10% DMSO (Sigma, D2650). The mini-pools were subsequently thawed and passaged once before final pooling.

### iPSC pooling and microglial differentiation

Microglial differentiation via Embryoid Body (EB) formation was performed according to Washer *et al.* (2022) ^14^. Briefly, iPSC cultures were dissociated into single cell suspensions using Accutase and resuspended in E8 medium containing 10 μM Y-27632. Cell counts were performed using Countess II (Thermo Fisher) and equal proportions of each iPSC line were pooled in E8 medium supplemented with 10 μM Y-27632, 50 ng/mL BMP-4 (Peprotech, 120-05ET), 20 ng/mL SCF (Peprotech, 300-07), and 50 ng/mL VEGF (Peprotech, 100-20A). The pooled suspension was seeded at 10,000 cells in 100 µL per well in Round Bottom Ultra-Low Attachment 96-well plates (Corning, 7007). Plates were centrifuged at 300 x g for 3 min and incubated at 37°C 5% CO2 for 3 days. Each pool contained between 16 to 72 donors.

On EB Day 3, 50 µL spent media was removed from each well and 100 µL fresh media (E8 medium supplemented with 10 μM Y-27632, 50 ng/mL BMP-4, 20 ng/mL SCF and 50 ng/mL VEGF) was added. An extra centrifugation at 300 x g for 3 min was performed if mini-satellite EBs were observed around the main EB. On EB Day 6, every 42 EBs were transferred to one T75 flask (Corning, 430641U) coated with 0.1% gelatin (Sigma, G1890) and cultured in Factory Medium: X- VIVO (Lonza, BE02-060F), supplemented with 2mM Glutamax (Life Technologies, 35050-061), 1:100 Pen/Strep (Life Technologies, 15140-122), 55μM 2-mercaptoethanol (Life Technologies, 31350-010), 100 ng/mL M-CSF (Peprotech, 300-25), and 25 ng/mL IL-3 (Cell Guidance Systems, GFH80). Factory flasks were maintained at 50mL volume with 50%-80% media changes every 3-4 days.

On Factory Day 35 – 57, the floating precursors in spent media were harvested for a 14-day microglial differentiation. Cells were sieved through 40 µm cell strainers (Falcon, 352340), centrifuged and pooled together. Cell counts were performed manually using disposable Neubauer-Improved haemocytometers (NanoEnTek, DHC-N01-50) and cells resuspended in ITMG media: Advanced DMEM/F12 (Life Technologies, 12634-010) supplemented with 2mM Glutamax, 1:100 Pen/Strep, 100 ng/mL IL-34 (Peprotech, 200-34), 50 ng/mL TGFβ1 (Peprotech, 100-21C), 25 ng/mL M-CSF, 10 ng/mL GM-CSF (Peprotech, 300-03- 20). Cell suspensions were seeded at 500,000 cells per 6-well (Greiner Bio-One, 657160) for single-cell RNAseq, 3,906,250 cells per T75 flask (Greiner Bio-One, 658170) for migration assay, or 1,126,400 cells per 6-well for phagocytosis assay. Media was topped up to double the volume on day 3 or 4, and subsequent half media change was performed on day 7 and day 10 or 11.

### iMacrophage differentiation

The floating precursors were harvested for a 7 day macrophage differentiation (Van Wilgenburg et al 2013). Precursors were resuspended in iMac media consisting of X-VIVO 15 (Lonza), supplemented with 2mM GlutaMAX, 100 ng/mL M-CSF. Precursors were incubated at 37℃ and 5% CO2 and differentiated for 7 days with a half media change on day 3 or 4.

### IFN**γ** and LPS stimulation

On day 11 of the 14-day microglial differentiation, cells were treated for 72 hrs with freshly prepared 20 ng/mL IFNγ (R&D Systems, 285-IF) or 100 ng/mL LPS (Sigma, L5668) diluted in ITMG media.

### Sample preparation for single cell RNAseq

On microglial differentiation day 14, the cells were washed twice with DPBS (-/-) (Life Technologies, 14190-094) dissociated with 5 mM EDTA (Life Technologies, 15575-038) supplemented with 4 mg/mL lidocaine (Sigma L5647). The cells were incubated at 37°C for up to 30 min before adding 0.04% BSA (Sigma A9543 or A8806) in DPBS to stop the dissociation. The cells were dissociated using a P1000 or 5mL stripette and collected into a centrifuge tube. After centrifugation, the cells were resuspended in 0.04% BSA and sieved through a 30 μm cell strainer (Miltenyi Biotec, 130-098-458) twice. After centrifugation, the cells were resuspended in 0.04% BSA and manually counted using Neubauer-Improved C- Chip disposable haemocytometer. The concentrations were adjusted to 449,612 cells/ml to load 17,400 viable cells per lane on Chromium Next GEM Single Cell 5ʹ Kit v2 (10x Genomics, PN-1000263) or 1,491,021 cells/ml to load 115,405 viable cells per lane on Chromium Next GEM Single Cell 5’ HT Kit v2 (10x Genomics, PN- 1000356).

Subsequent GEM handling and library preparations were performed according to the manufacturer’s instructions. Libraries were QC’ed using Qubit dsDNA HS assay (Life Technologies, Q32854), BioAnalyser (Agilent, 2100) or TapeStation (Agilent, 4150) and/or KAPA library Quant Kit for Illumina (Roche, KK4824). Multiplexed libraries were sequenced using HiSeq or NovaSeq 6000 (Illumina) with read length 28-10-10-90, 26-10-10-90, 100 paired-end or 150 paired-end, aiming for a depth of at least 20,000 reads per cell.

### Migration assay

Microglia were seeded from precursors of Factory Day 39 – 43 at a density of 3,906,250 cells per T75 flask (Greiner Bio-One, 658175). On microglial day 14, the cells were dissociated with 5 mM EDTA (Life Technologies, 15575-038) supplemented with 4 mg/mL lidocaine (Sigma L5647). AdvDMEM/F12 was added to stop the dissociation and the cell suspensions were sieved through 30 µm cell strainers (Miltenyi Biotec, 130-098-458). The centrifuged cell pellets were resuspended in ITMG media and cell count were performed manually using Neubauer-Improved C-Chip disposable haemocytometer. For each 6-well insert (Sarstedt, 83.3930.500), 2.7mL suspension containing 825,000 cells was seeded to the inner well. After cells were settled for 15 min at room temperature, 3.168 mL ITMG media with or without 3 nM C5a (R&D Systems, 2037-C5-025/CF) was added underneath each 6-well. After incubation at 37°C for 6 hrs, genomic DNA was harvested and sister wells were fixed in 4% PFA for imaging.

Genomic DNA from upper and lower sides of the transwells were harvested using FLOQSwabs (COPAN, 520CS01) and cell pellets were collected from the washes of each side of the transwells. Corresponding swabs and pellets were combined for genomic DNA extraction using Puregene Cell Kit (QIAGEN, 158043) according to manufacturer’s instructions.

To quantify the migration rate, whole well imaging was performed on EVOS Fl Auto (Life Technologies) after fixation in 4% PFA and staining with NucBlue™ Live ReadyProbes™ Reagent (Life Technologies, R37605). After imaging all cells, the unmigrated portion on top of the transwells was removed with swab and the migrated portion at the bottom of the transwells was imaged. The number of nuclei in *all-cells* images was quantified using Cellpose (version 2.0.5) with -- diameter 5 --flow_threshold 0 --cellprob_threshold -8. The number of nuclei in *migrated-cells* images was quantified using Cellprofiler (version 4.2.1) with the threshold strategy set as Global, threshold method set as Robust background, lower bounds on threshold set to 0.015, and upper bounds on threshold set to 1.0. The migration rate was calculated as the number of nuclei in *migrated-cells* divided by the number of nuclei in *all-cells*. The percentage of chemotaxis was calculated by subtracting the cell migration rate in -C5a condition from the cell migration rate in +C5a condition.

Generation of mCherry-eGFP pmChGIP SH-SH-SY5Y, maintenance & fixation mCherry-eGFP fusion SH-SY5Y were generated as previously described in Washer *et al.* 2022, and are henceforth referred to as pmChGIP SH-SY5Y, clone F4 was used for all experiments. pmChGIP SH-SY5Y were maintained in DMEM/F12 (ThermoFisher, 11320033) supplemented with 2mM GlutaMAX (ThermoFisher, 35050061) and 10% FBS (Gibco, 10500-064) at 37°C 5% CO2.

Cells were maintained until reaching confluency before washing with DPBS (-/-) to remove dead cells and lifted with TryPLE Express (Life Technologies, 12604- 013) and harvested into 50mL tubes. Flasks were washed with suspension containing harvested pmChGIP SH-SY5Y. Cells were centrifuged at 400g for 5 min at room temperature before removing supernatant and resuspending with 1 mL DPBS (-/-). Cells were then adjusted to 8x10^6^ cells/mL in DPBS (-/-) before adding an equal volume of 4% PFA (1:1 dilution, final concentration 2%) mixed vigorously with a stripette and incubated at room temperature for 10 min, with gentle vortexing every 2 min. The 2% PFA/cell mix was diluted to 1% by adding an equal volume of DPBS (-/-) before centrifuging at 1200g for 5 min at room temperature. The supernatant was removed and the pellet resuspended 5mL DPBS (-/-) before repeating the centrifugation and resuspension a further two times (total three). The fixed pmChGIP-SHSY5Y were then counted and stored at 4°C overnight. Prior to addition to iMGL, the pmChGIP-SH-SY5Y were resuspended in ITMG media.

### Dual colour phagocytosis assay

Microglia were seeded from precursors of Factory Day 46 – 57 at a density of 1,126,400 cells per 6-well (Greiner Bio-One, 657160). On microglial day 14, fixed SH-SY5Y cells were added to microglia in ITMG media at a ratio of 2:1 SH- SY5Y:microglia. After incubation at 37°C for 2 hrs, wells were washed with HBSS (+/+, Life Technologies, 14025-050) to remove cargos and with DPBS (-/-) before lifted with TryPLE express (Life Technologies, 12604-013) and collected into 0.04% BSA. The cell suspension was fixed in 4% PFA and washed with DPBS twice before resuspended in 0.04% BSA for sorting on Mo-Flo XDP (Beckman Coulter).

Genomic DNA extraction was extracted from fixed cells using Puregene Cell Kit according to manufacturer’s instructions and quantified using Qubit dsDNA HS assay.

### Genomic DNA library prep for phenotypic assays

Genomic DNA extracted from migration and phenotypic assays was quantified using Qubit dsDNA HS assay. Library prep was performed using NEBNext® Ultra™ II FS DNA Library Prep Kit for Illumina (NEB, E7805) with NEBNext® Multiplex Oligos for Illumina® (Dual Index Primers Set 1) (NEB, E7600) following manufacturer’s instructions. Libraries were QC’ed using Qubit dsDNA HS assay, BioAnalyser or TapeStation and/or KAPA quantification qPCR. Multiplexed libraries were sequenced using NovaSeq 6000 (Illumina).

### Cloning of Cas9-guide all-in-one vector

Three guide pairs per target (plus intergenic controls) were cloned into pLentiCRISPRv2 (Addgene https://www.addgene.org/52961) with a modified guide scaffold (Addgene https://www.addgene.org/50946/) in an arrayed format. Briefly, forward primers containing the 5’ guide, homology arms to plasmid backbone and dual guide cassette^76^ and reverse primers containing the 3’ guide, homology arms to plasmid backbone and dual guide cassette were synthesized by IDT in an arrayed format.

*Sequence of forward primers:*

tatcttgtggaaaggacgaaacaccGNNNNNNNNNNNNNNNNNNNgtttcagagctagaaatagcaagttg

*Sequence of reverse primers:*

gctgtttccagcatagctcttaaacNNNNNNNNNNNNNNNNNNNCTGCATTGGCCGGGAATTGAAC

The dual cassette flanked by dual guides and homology arms was PCR amplified with the above primers and a template plasmid containing the dual guide cassette, using Q5 High-Fidelity 2X Master Mix (NEB, M0492) with an annealing temperature of 64°C.

*Sequence of dual guide cassette:*

gtttcagagctagaaatagcaagttgaaataaggctagtccgttatcaacttgaaaaagtggcaccgagtcggtgcGCAGAGGCATTGGTGGTTCAGTGGTAGAATTCTCGCCTCCCACGCGGGAGaCCCGGGTTCAATTCCCGGCCAATGCAg

The gel extracted (NEB, T1020) or column purified (NEB, T1030) PCR was cloned into Bsmb1-V2 (NEB, R0739) digested pLentiCRISPR v3.0 backbone using Gibson Assembly® Master Mix (NEB, E2611) at a ratio of 3:1 insert:backbone. The Gibson product was transformed into DH5-α cells (NEB, 2987U) and clones were miniprep using Qiaprep 96 plus miniprep kit (QIAGEN, 27291). The sequence of each clone was verified by sequencing with universal U6 primer.

Post arrayed cloning, library was pooled and the representation was quantified by MiSeq (Illumina) before expansion using electrocompetent cells (Lucigen, 60242-1) to ensure sufficient coverage and maxiprep using ZymoPURE™ II Plasmid Maxiprep Kit (Zymo, D4203). For Miseq verification, a nested PCR was performed using the following primers and KAPA HiFi HotStart ReadyMix (Roche, KK2602).

PCR1 forward primer: ACACTCTTTCCCTACACGACGCTCTTCCGATCTCTTGTGGAAAGGACGAAACA

PCR1 reverse primer: TCGGCATTCCTGCTGAACCGCTCTTCCGATCTttcccactcctttcaagacc 12 cycles of 98°C 20s, 60°C 15s and 72°C 15s

PCR1 was column purified (NEB, T1030) and 1/100 was used as template for PCR2.

*PCR2 forward primer:*

AATGATACGGCGACCACCGAGATCTACACTATAGCCTACACTCTTTCCCTACACGAC GCTCTTCCGATCT

*PCR2 reverse primer:*

AATGATACGGCGACCACCGAGATCTACACTATAGCCTACACTCTTTCCCTACACGAC GCTCTTCCGATCT

14 cycles of 98°C 20s, 70°C 15s, 72°C 15s

### Production of lentiviral particles

Lenti-X HEK293T (Takarabio, 632180) were maintained in HEK media (DMEM (Corning, 15-013-cv), 10% FBS, 2mM GlutaMAX) at 37°C 5% CO2 and split when at 70-80% confluency using TrypLE Express following standard practice.

Prior to seeding for lentiviral production, plates were coated with 0.1% Gelatin/dH2O (Sigma, G1393) for 1 hour at 37°C. For large scale production Lenti- X HEK293T were seeded in 15cm dishes at 14x10^6^ cells/dish in 30mL HEK media, for small scale production in 6wp. 18-24 hours post seeding the confluency was checked (approx 70%) before transfecting with 7.5μg pSIV3+ or lentiviral library, 18.5μg psPAX2 (Addgene, 12260), and 4μg pMD2.G (Addgene, 8454) in 7.5mL OptiMEM (Invitrogen, 51985026) with lipofectamine LTX (Invitrogen, 15338100) per 15cm plate, topped up to 15mL with OptiMEM. 24 hours post transfection, the transfection mix was replaced with HEK media.

At 48 hours post transfection viron containing media was harvested into 50mL tubes and replaced with fresh HEK media. Viron media was centrifuged at 500g for 5 min to pellet any HEK cells, before filtration through 0.45μm filters. Filtered viral supernatant was split between 50mL tubes and concentrated through centrifugation at 20000g for 3 hours at 4°C. Supernatant was removed and the viral pellets resuspended on ice for 1 hour in 1.5% BSA/DPBS. Resuspended virons were combined and stored at 4°C. The process was repeated at 72 hours post transduction before combining both the 48 hour harvest and 72 hour harvest and split into single use aliquots and stored at -80°C. A full method can be found in Washer *et al.* 2025.

### Total protein extraction and western blotting

At d14 ITMG media was removed and either 50μL (96wp) or 100μL (48wp) RIPA (Thermofisher, 89900) containing protease inhibitor (Roche, 11836170001) was added to each well and the plate incubated on ice for 10 min. Lysates were transferred to a 96 well PCR plate, sealed, and vortexed for 20 seconds. The plates were centrifuged at 5000 rpm for 5 min to pellet any cell debris and lysates transferred to a clean 96 well plate and stored at -80°C.

Total protein was calculated by BCA assay following manufacturers instructions (Thermo Scientific, 23225). 10-40μg of protein was then diluted in water, NuPAGE reducing agent (Invitrogen, NP0009) and NuPAGE LDS sample buffer (Invitrogen NP0007), boiled at 95°C for 5 min and loaded onto a 4-15% Mini- PROTEAN TGX precast gel (BioRad, 4561086) with 4μL Spectra Multicolor High Range Protein Ladder (Thermo-scientific, 26625). Samples were run at 100V for 75 min. Protein was transferred to a PVDF membrane (activated in methanol for 2 min) at 100V for 75 min. Membranes were then blocked for 1 hour at RT with 5% Milk PBS-Tween (PBST). Membranes were washed with PBST before staining with primary antibody made up in 1% Milk PBST overnight at 4°C. Primary antibody was removed and membranes washed with PBST before incubating with the HRP conjugated secondary antibody (prepared in 1% Milk PBST) for 1 hour at room temperature. Membranes were then washed and stained with enhanced chemiluminescence substrate for 5m before imaging.

For data analysis images underwent densitometry in ImageJ. Target protein levels were normalised to respective housekeeping controls before being normalized to the untreated control to give “Relative Protein Expression (to Control)”. All data analysis was performed in R.

### CRISPR KO screening for phagocytosis

KOLF2.1S hiPSC were differentiated to iMGL as described above. The library contains 783 dual-guide all-in-one CRISPR vectors targeting 251 AD GWAS hits and intergenic controls, henceforth known as NeuroKO. Genes were chosen from prioritisation in Schwartzentruber *et al*.^5^, and OpenTargets Locus2Gene score^58^ for the additional loci discovered in Bellenguez *et al.*^8^ Guide RNA pairs were designed using anchor guides chosen using the prioritisation scheme from MinLib^77^ and paired with guides targeting the same exon using a tRNA-based dual guide system^78^. In order to maintain high coverage of the library, between 1.1 x 10^8^ and 1.23 x 10^8^ precursors were transduced at a multiplicity of infection between 1.2 - 1.5 along with a viral accessory protein (VPX), which has been shown to improve transduction efficiency in this model^76^. We performed 3 replicates of the screen, defined as 3 separate transductions and harvests of the precursors.

After differentiation to iMGL for 14 days (14d) in ITMG we then incubated the iMGL with pmChGIP SH-SY5Y at a ratio of 2 pmChGIP SH-SY5Y to 1 iMGL and phagocytosis was undertaken for 6 hours. Non-phagocytosed cargo were washed, iMGL lifted, and fixed with 2% PFA, before sorting on the mCherry signal in four bins: the top 20% (top T20), 55-75% (upper middle UM), 25-45% (lower middle LM) and bottom 20% (lowest L20). In total between 0.92 x 10^6^ and 1.7 x 10^6^ cells were sorted into each bin and a total of between 4 x 10^6^ - 6 x 10^6^ total cells sorted across the four bins in the three screens. Following DNA extraction and assuming 6pg of DNA per cell, the coverage of screen across the three repeats were estimated to be between 263 and 454 per bin.

dgRNA sequences were amplified and indexed by PCR and sequenced by Illumuna NovaSeq. Following deconvolution, dgRNA sequences were quantified by MAGeCK^79^. Mapping efficiency and guide representation by gini index was examined to confirm the quality of the data using MAGeCKFlute^80^ and are shown in (Supplementary Fig. 22). Following this, the guide overrepresentation was calculated between the L20 and T20 populations, representing lowest phagocytosing and highest phagocytosing iMGL to provide a list of hits. Full methods are available at Washer *et al.* (2025)^70^.

### Arrayed Lentiviral Screening

AOWH_2, HEGP_3, and KOLF2.1S lines from the HipSci repository (www.hipsci.org) were used for all CRISPR screening validation.

3 sgRNA lentiviruses to TREM2 and Intergenic regions were cloned into pLentiCRISPR v3.0 as described above and individual viruses were generated as previously described. Individual virus titres were calculated by resazurin survival following puromycin selection on KOLF2.1S iMGL. Following successful titring the 3 TREM2 sgRNA were pooled, and the 3 INT sgRNA were pooled and stored at - 80°C.

AOWH_2, HEGP_3, and KOLF2.1S were transduced as follows. PreMacs were harvested from respective factories (age 32d, 36d, 42d) and adjusted to a final concentration of 4.84x10^5^ cells/mL in ITMG. Polybrene was added to a final concentration of 4μg/mL and inverted to mix. 200μL of cells were aliquoted into 48wp (Greiner, 677180) to give 9.68x10^4^ cells/well (WT). VPX was added to the remaining cell suspension at the determined concentration for full KD of SAMHD1, cells were inverted and 200μL of cells were aliquoted to give a VPX control (VPX). The remaining cell suspensions (containing polybrene and VPX) were split between two 15mL tubes and either TREM2 lentiviral pools (TREM2) or INT lentiviral pools (INT) were added, cells were inverted and 200μL of cells were aliquoted. Cells were incubated at 37°C 5% CO2. 24hr post transduction, all media was removed and replaced with 200μL fresh ITMG. On d4/5 post-transduction 200μL fresh ITMG was added to each well, a subset of wells were selected with puromycin (at final concentration 1μg/mL) to enrich for transduced cells and to check transduction efficiency (TREM2 - Puro, INT - Puro). Two further 50% media changes were undertaken on d7 and d10 post transduction with assays completed on d14.

Protein was harvested as previously described and underwent western blotting for TREM2 and Vinculin to confirm levels of TREM2 KD. The remaining wells were fed the dual fluorescent reporter pmChGIP SH-SY5Y line for 6 hours before washing, lifting, fixing with 2% PFA, and undergoing flow cytometry as previously described.

### TREM2 ASO Knockdown

An overview of the ASO experiments is provided in Supplementary Fig. 23. TREM2-171, TREM2-192, Scramble 1, and Scramble 2 ASO sequences from Vandermeulen *et al.*^81^ were ordered with modifications from IDT and resuspended to 100μM in DPBS (-/-) and stored at -20°C.

To obtain IC50 and IC90 curves, KOLF2.1S preMics were seeded in 96 well plates (Greiner 655180) at 3x10^4^ cells/well in 96 well plates in 100μL ITMG. 72hr post seeding 50μL of ITMG was replaced with 50μL fresh ITMG. 7 days post seeding 50μL ITMG was removed and individual TREM2 ASO were added serially in triplicate to give final concentrations (20μM, 5μM, 1.25μM, 312nM, 78.1nM, 19.5nM, 4.88nM, 1.22nM, 305pM, 76.3pM), in 100μL ITMG. Another 50% media change occurred on d10 before harvesting total protein on d14. IC50 and IC90 were calculated in R using the *drm()* function from the *drc* package (version 3.0-1) (Supplementary Fig. 21)^82^.

For validation experiments the above protocol was repeated at the respective IC50 and IC90 doses for TREM2-171 and TREM2-192, with the scramble controls matching the respective IC doses for each TREM2 targeting ASO.

AOWH_2, HEGP_3, and KOLF2.1S were used as follows. PreMics were harvested from respective factories (age 32d, 36d, 42d) and adjusted to a final concentration of 4.84x10^5^ cells/mL in ITMG before plating 200μL in 48wp (Greiner, 677180). At d3-4 post seeding, a 50% media change was undertaken. At d7 ASO were diluted at 2x concentration in ITMG before removing 100μL from the cells and adding 100μL 2x ASO in ITMG (final concentration 1x). At d10 a 50% media change was undertaken and assays completed at d14.

Protein was harvested as previously described and underwent western blotting for TREM2 and Vinculin to confirm levels of TREM2 KD. The remaining wells were fed the dual fluorescent reporter pmChGIP SH-SY5Y line for 6 hours before washing, lifting, fixing with 2% PFA, and undergoing flow cytometry as previously described. TREM2 protein levels and phagocytosis are shown as relative to vehicle control.

### Genotype imputation and quality control

Raw genotyping array files were obtained per line from the HipSci project (http://www.hipsci.org) and from IPMAR project^13^, and imputed using the *<u>genimpute</u>* pipeline from the eQTL Catalogue. Briefly, it converts genome coordinates from GRCh37 to GRCh38 with CrossMap v.0.4.1^83^, aligns them with 1000 Genomes Project High Coverage^84^ reference panel with Genotype Harmonizer v.1.4.20^85^, excludes variants with Hardy-Weinberg p-value < 1e-6, missingness > 0.05 and minor allele frequency < 0.01 with bcftools^86^, performs genotype pre-phasing with Eagle v.2.4.1^86,87^, and imputation with Minimac4^88^. Post-imputation QC excludes variants with imputation r^2^ < 0.4, keeps variants on chromosomes 1-22 and X, retains variants with MAF > 0.01, and multiplies genotype dosage of male samples on the Non-PAR region of the X chromosome by two. Sex was imputed for donors that were missing this information using bcftools *+guess-ploidy* on region X:2781480-155701381.

### Donor proportion deconvolution by whole genome sequencing

We used whole genome sequencing (WGS) to measure the proportion of the different donors within each sample. This method (named *poodleR*) requires that donors are genotyped, and then variants subset to those that present non- identical genotypes across all donors per individual pool. WGS files had duplicated reads removed with Picard’s *MarkDuplicates*^89^, and alleles were counted at the non-identical variants detailed before using Bam-readcount^90^. Alternative allele frequencies were estimated (only for the most abundant minor allele in the case of multiallelic variants, and only for point mutations) for all predefined variants covered by at least one WGS read.

Finally, at each variant *i* the alternative allele frequency *b* is the sum across the total number of donors in the pool (*k*) of the product of each donor’s (*j*) minor allele dosage *a* and proportion *w*:

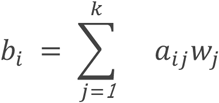

Extending this for all donors and positions requires the genotype matrix A coded as minor allele dosages:

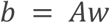

Donor proportions were calculated by least-square regression:

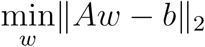

With constraints:

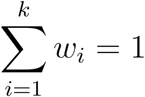 and 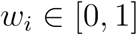 [0,1] for all *i*.

### Accuracy estimation of donor proportion deconvolution

We simulated WGS pools with known donor proportions sequenced at a variety of mean depths to estimate the performance of the method. For that, we first estimated the mean depth and number of reads from individual donor WGS sequencing files, and calculated the total number of reads needed to achieve the target mean depth. Then, we calculated the number of reads needed per donor per mean depth to achieve a target donor proportion in a simulated pool, and sampled those reads randomly from individual donor WGS sequencing files before merging them for all donors into a simulated pool of reads. We finally counted the occurrence of each minor allele over the total number of reads, and estimated the donor proportions.

We did this for a range of proportions per mean depth: equal proportions for ten donors, equal proportions (0.2) for five donors where other five were zero (ten donors in total), and at a range of proportions from 0.0005 to 0.39 from 19 donors to simulate a real-life scenario.

We then estimated the absolute error (Supplementary Figure 1a) comparing the real proportions we sampled to those estimated by the method. The method’s maximum absolute error is 3% from a mean depth of 0.25X onwards, across the range of unequal proportions for a representative pool of 19 donors. The other two scenarios present a lower absolute error.

Relative error was calculated as abs((real proportion - estimated proportion) / real proportion)*100 (Supplementary Figure 1b). Relative accuracy was measured as 100 - relative error. We observed that at higher (over 10%) real proportions the estimated proportions were always underestimated. Under 10% real proportions, these were always overestimated. This was taken into account to adjust the real donor proportions from our pools, as described in the next section.

### Estimation of differentiation efficiency, migration, and phagocytosis cellular phenotypes

WGS data from 14–15 pooled experiments, comprising 243 donor iPSC lines (19– 24 per pool), was generated for the cellular phenotypes (differentiation efficiency, migration or phagocytosis). Donor proportions within pools were estimated using *poodleR* as described above. Then the cellular phenotypes were calculated as follows. First, the raw donor proportions were adjusted upwards or downwards, by adding or subtracting the absolute error estimated at the closest simulated proportion (based on the simulations described in the previous section).

For the differentiation efficiency phenotype, log-fractions were calculated per line, pool and replicate as log(proportion at later stage /proportion at earlier stage) for macrophage precursors vs iPSC, old vs young macrophage precursors, and microglia vs macrophage precursors, the latter while matching the ages of the macrophage precursors from which they originated (see Supplementary Table 2). Then log-fractions were scaled per replicate.

For the phagocytosis phenotype log-fractions were calculated per line, pool and replicate as log(proportion mCherry+ /proportion mCherry-), then scaling per replicate. Of note, these donor proportions were deconvoluted including the genotype of the SH5Y5Y line that is added to the pools during the phagocytosis assay. For the migration phenotype log-fractions were calculated as log(proportion bottom side of transwell / proportion top side of transwell) per line, pool and replicate in the C5a+ and C5a- conditions separately, then the C5a+ fractions were normalised to C5a- fractions per replicate, and finally they were scaled per replicate.

### Functional variant effect annotation of WES

Whole exome sequencing (WES) files for 226 available lines present in our study were filtered as in Rouhani *et al.*^24^ and only high-quality variants (with PASS filter) were retained. Functional variant effects were annotated using the variant effect predictor^91^ (VEP, v. 111) for each line. We defined gene coordinates as those present in the Ensembl gene annotation (GRCh37, v. 102). After variant annotation, variant files were merged with bcftools v. 1.19 and coordinates lifted over with Picard v. 2.26.2. Variants were then classified as deleterious (69,223 variants across 16,348 genes) if they were annotated as frameshift, stop gained, transcript ablation, splice acceptor or splice donor; or if they were annotated as missense, start lost and protein altering and had CADD Phred scores (v. 1.7) over 15. All other missense variants were considered missense non-deleterious (simply “missense” – 62,022 variants across 15,523 genes).

### Test of deleterious and missense variant effects on cellular phenotypes

Gene-level effects of missense and deleterious variants were tested for each phenotype (averaging the scaled phenotype per line across pools and replicates) using the sequence kernel association optimal test (SKAT-O)^92^, combining SKAT and burden tests. Empirical p-values were calculated by resampling the residuals 1000 times from the null model (using bootstrapping) while adjusting for covariates: sex, average of the minimum line proportion per replicate, and the first two genotype principal components (and the scaled differentiation from macrophage precursor to microglia in the case of the phagocytosis and migration phenotypes, to control for possible effects of differentiation on these two).

Since SKAT-O does not provide a beta estimate on the phenotype, we calculated it through burden tests only for SKAT-O significant genes, by performing linear regression aggregating the variants per gene as done in Puigdevall *et al.*^3^ and fitting a linear mixed model with the following formula using *lmerTest* in R:

~~~
Scaled phenotype (phagocytosis or migration) ∼ deleterious burden + sex + minimum line proportion in replicate + scaled differentiation + genotype PC1 + genotype PC2 + (1|pool)
~~~

For the scaled differentiation efficiency phenotype (which reflects the proliferation, survival and differentiation capacity of each line), we fitted the same formula without including scaled differentiation as a covariate, for the effects on iPSC to macrophage precursors, and young to aging macrophage precursors:

~~~
Scaled differentiation efficiency ∼ deleterious burden + sex + minimum line proportion in replicate + genotype PC1 + genotype PC2 + (1|pool)
~~~

For the effects on macrophage precursors to microglia the test was performed in the same way including an additional `(1|treatment)` term. P-values were corrected by Bonferroni adjustment for multiple testing. Significant genes after Bonferroni correction for every comparison had their deleterious burden scores aggregated together, separated by the directionality (positive or negative) of the beta, and tested using the same model above.

AUC was calculated by building a binary classification of “dropouts” for lines with scaled differentiation efficiency more than 2 standard deviations below the mean, or of “takeovers” in the opposite direction, and then testing the aggregated burden of deleterious variants in significant genes in the relevant direction as predictors using *roc()* and *auc()* in the *pROC* package^93^.

Significant differentiation genes as tested by SKAT-O were tested for enrichment in DepMap essential genes and macrophage survival genes from Covarrubias *et al*.^27^ by Fisher’s exact test (Supplementary Table 14).

### Single cell RNA-seq processing and line identity deconvolution

FASTQ files were processed with 10x Genomics Cell Ranger v.6.0.1^94^ using default parameters.

In preparation for line identity deconvolution in scRNA-seq, the reference genotype files were subset to exons using vcftools v.0.1.16^95^. Then variants were subset to those that had non-identical genotypes across all lines per individual pool. Single cell RNA-seq deconvolution was performed from the cellranger BAM outputs with Vireo v.0.5.8^96^ and cellSNP-lite v.1.2.2^97^ using default parameters.

3,404,020 cells were processed in Seurat v.5^98^ (Supplementary Table 15), filtering out low quality cells (those with more than 10% mitochondrial genes, those that presented a number of UMI counts under the 5th percentile, doublets and cells with unassigned line identity; Supplementary Figure 1a). This left in total 2,341,497 cells remaining across all pools and treatments. Each pool per treatment was log normalised using *NormalizeData()* in Seurat, and the 2,000 most variable genes were selected for integration, which was performed on a “sketched” subset of 5000 cells, sampling cells without replacement while incorporating rare cell types with *SketchData().* For the subset of cells, variable genes were selected again and their expression scaled, before dimensionality reduction and clustering using PCA, finding shared nearest neighbors (SNN), and performing modularity optimization based clustering with the Louvain algorithm. Finally, we performed dimensionality reduction by Uniform Manifold Approximation Projection (UMAP) using default parameters, providing the foundation for the integration across all pools using Harmony with *IntegrateLayers()*. The remaining cells were then projected onto the pre- integrated transcriptomic manifold using *ProjectIntegration()* and *ProjectData()*. The integrated UMAPs (per treatment and merged) were used for cell subtype annotation with individual marker genes and differential expression gene sets from Mancuso *et al.* (2024)^15^ by using Seurat’s *AddModuleScore()*. Expression heatmaps and markers’ expression dotplots and UMAPs were plotted with SCpubr^99^ and scCustomize^100^

### Polygenic risk score calculations

Polygenic risk scores (PRS) were calculated with PRSice2. First, we excluded the APOE regions from the genotype file from the HipSci and IPMAR lines. Then PRS was calculated using the AD GWAS summary statistics (from Bellenguez *et al.*^8^) and the LD information from the European ancestry (EUR) subset from 1000 Genomes, using a p-value threshold of 0.1 and leaving other options as default. We then calculated the PRS for the EUR subset from 1000 Genomes, using the same variants from the HipSci and IPMAR lines genotype file. We adjusted the raw scores for 5 genotype PCs for both datasets, and scaled the HipSci and IPMAR lines’ scores to those of 1000 Genomes EUR. We then added the scores for the APOE alleles with values B(APOE e2)= -0.47, B(APOE e3)=0, B(APOE e4)=1.12 as in Leonenko *et al.*^101^ and scaled them to those of 1000G EUR. Finally we added the scaled polygenic and APOE scores to obtain the full PRS. The distribution of PRS in the HipSci, IPMAR, and 1000G EUR lines are shown in Supplementary Figure 12a-c.

For microglia-specific PRS, regions 250Kb around the lead colocalizing loci between eQTLs and AD GWAS were taken (per treatment), as used as an input for PRSice2, both for data from our 261 donors and for that of 1000 Genomes EUR used in the scaling. All other steps to build the scores were identical. The distribution of microglial-specific PRS is shown in Supplementary Figure 12e.

### Differential expression analysis across treatments

Raw UMI counts were aggregated by line, pool, and treatment replicate for the non-proliferative clusters. We then removed samples with < 100 cells (leaving 194 lines for analysis) and retained genes that were present in at least 30% of the donors with at least 1 count per million (CPM) (Supplementary figure 2a). We then transformed the data to log2-counts per million and calculated sample-level weights from the mean-variance relationship using *voom()* from *limma*^102^, while adjusting for pool random effects with *duplicateCorrelation()*. Finally a linear model was fit for the full dataset including donor effects and log10(number of cells) as covariates. The final fit was as follows:

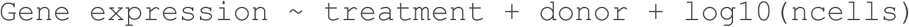

Sex and age effects were not fit as they were confounded with donor, and age was not confidently recorded in several cases. We then tested every treatment contrast (LPS vs untreated, IFN𝛄 vs untreated, and IFN𝛄 vs LPS) before moderated t-statistics were calculated with *eBayes()*. False discovery rate (FDR)-adjusted p-values were calculated per gene using the Benjamini- Hochberg procedure. Log2 fold changes in expression take the second element in the contrast as the baseline. The final numbers of differentially expressed genes are:

| | IFNγ vs Untreated | LPS vs Untreated | IFNγ vs LPS |
| --- | --- | --- | --- |
| Down | 926 | 1144 | 975 |
| Not sig. | 11046 | 10736 | 10719 |
| Up | 800 | 892 | 1078 |

### Differential gene expression across PRS

Raw UMI counts were aggregated by line, pool, and treatment replicate as in the differential expression analysis across treatments. Genes and lines were QC’d, counts were transformed and pool was accounted for in the same way. We fitted the following linear mixed model per treatment:

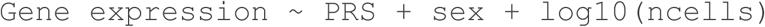

We did not fit age for the reasons mentioned before, and because it is confounded with high PRS for those IPMAR donors with AD diagnosis. We then tested PRS effects, for which each gene’s coefficients are interpreted as log2 fold changes per unit increase in PRS. This resulted in the following number of significant (FDR-adjusted p-value < 0.05 & abs(log2FC)>0) DE genes:

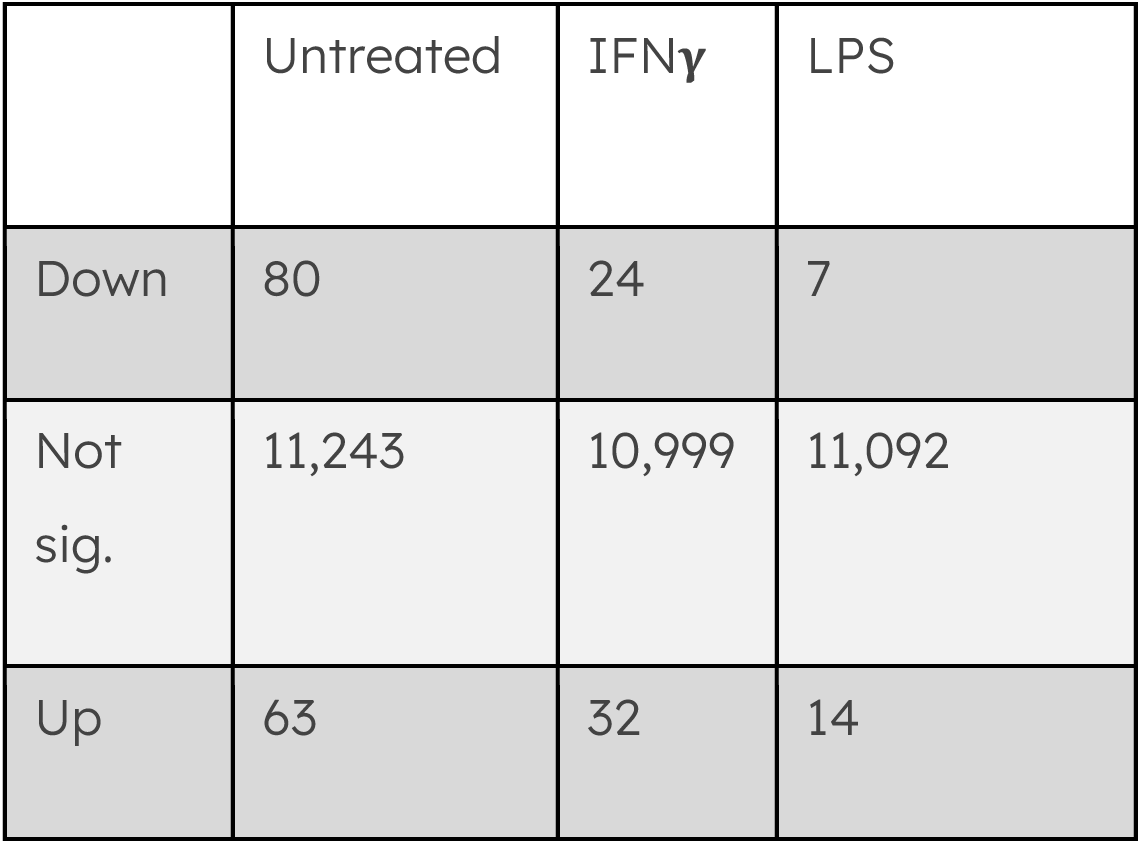

Empirical p-values were calculated for each gene per treatment by shuffling each line’s PRS 100 times, sampling the t-statistics per gene and calculating p- values by estimating how often the t-statistic from the real set of PRS was more extreme than that of the permuted set. Log2 fold changes in expression are referred to each unit change in PRS.

### Differential gene expression across phenotypes

UMI counts were aggregated and lines filtered as in the previous section. Afterwards, we averaged across replicates the scaled phenotypes (migration and phagocytosis) and the minimum line proportion of the fraction to account for higher variance of the fractions from lines that present in smaller proportions in the pool. Finally the differential expression analysis was performed per treatment with *limma*, accounting for pool effects as in the previous section and fitting the following regression model:

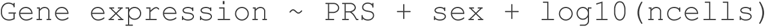

Genes with FDR-adjusted p-value < 0.05 and abs(log2FC)>0 were considered significantly differentially expressed. Log2 fold changes in expression are referred to each unit change in the scaled phenotype.

### Enrichment analysis in differential expression and eQTL results

Signed t-statistics from differential expression results were centered to ensure a normal distribution before enrichment analysis. For transcription factor activity inference, an univariate linear model was fitted per treatment using the decoupleR package and the <u>CollecTRI</u> network of weighted TF-target relationships. <u>PROGENy</u> pathways were fitted in the same way for the differentially expressed genes between treatments. Reactome hallmark pathways and candidate gene sets were tested with Gene Set Enrichment Analysis using the *msigdbr* and *fgsea* packages in R, and BH-adjusted p-values are reported. AD and PD candidate genes were defined by the prioritised gene sets from Schwartzentruber *et al.* ^7^ for AD, and from the top 2 genes from the Open Targets locus2gene score per associated locus for PD.

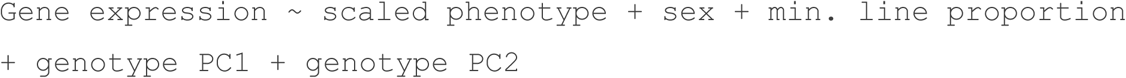

### Filtering genotypes and lines for eQTL and phenotype association analysis

Genotype quality control included filtering out variants with imputation INFO < 0.7, HWE<10^-6^, missing in > 10% of samples, and MAF < 5% in our samples. We also excluded lines with over 10% missing genotypes, which resulted in 6,405,518 variants used for eQTL and association analyses. We then filtered the 1000G genotype file in the same way and performed joint principal component analysis. The second line in each pair of clones (letw_5, lizq_3, zaie_1, romx_2, seru_7, qonc_2, sebn_4) and those lines that did not map close to European populations (boqx_2, garx_2, sojd_3, yoch_6) were excluded from the analysis (Supplementary Figure 12d). This left 250 donors for downstream analysis.

### eQTL mapping

Single cell RNA-seq counts were aggregated into pseudobulks per donor, and grouping together all non-proliferative clusters. This decision was driven by two factors: first, the lack of clearly defined subtypes within each treatment condition; and second, the observation that the number of cells comprising a given cluster strongly correlates with the number of detectable differentially expressed genes and eQTL-associated genes (eGenes)^75^. We then performed several quality control filters for the pseudobulks: at the gene level, retaining genes that were present in at least 30% of the donors with at least 1 count per million (CPM), and log-normalising and scaling gene counts; and at the line level, removing those lines with < 100 cells per pseudobulk and removing the second individual in a pair of clones, and those lines for which we didn’t have genotype information or that mapped outside of European populations when projected within the PCA of 1000 Genomes data. This left 188 (IFN𝛄 and LPS) and 189 (untreated) lines for analysis in the Non-proliferative group.

We included two genotype PCs and 5-120 expression PCs as covariates in the linear regression model. eQTL analysis was performed with *tensorQTL* v.1.0.9^104^ including variants in the *cis* neighbourhood of each expressed gene (250kb up and down the transcription start site, 500kb total). To correct for the number of tests performed per gene we used tensorQTL in the *map_cis* mode, which permutes gene expression values a thousand times to obtain empirical false discovery rates (FDR) per gene-variant tested. We then used q-value correction on the genes tested to obtain a final set of significant (FDR<0.05) eGenes. The final set of expression PCs used maximized the number of significant eGenes: 63 for the untreated sample, 73 for IFN𝛄, and 84 for LPS.

The degree of eQTL sharing was calculated by multivariate adaptive shrinkage with *mashR*, subsetting the results per treatment to the lead eQTL-eGene pairs, gathering these pairs from other treatments if not present, and estimating the lfsr and posterior means. eQTL were deemed shared if they had lfsr < 0.05 and if they had posterior means (betas) within 0.5 and the same sign.

### Genome-wide association analysis of cellular phenotypes

Genotypes and lines were processed as detailed for eQTL analyses. To avoid inflation of false positives due to repeated line measurements and to limit the effect of the more variable phenotype fractions of less abundant donors, samples per line, pool and replicate, were filtered to those in which the smallest proportion of the pair in the fraction reached at least 1% abundance, and then the scaled fractions were averaged across replicates and pools. This left 133, 137, and 140 lines left for testing in the untreated, IFN𝞬, and LPS treatments, respectively, in the phagocytosis phenotype. In the migration phenotype, there were 147, 92, and 147 lines left for testing in the untreated, IFN𝞬, and LPS treatments, respectively. In the differentiation phenotype, there were 209, 229, and 230 lines left for testing in the untreated, IFN𝞬, and LPS treatments, respectively. Association tests were performed by fitting a linear regression model with the following formula:

~~~
Mean phenotype across pools ∼ Alternative allele dosage + sex + number of pool replicates + genotypePC1-5
~~~

### Colocalization analysis of eQTL and GWAS

External disease GWAS and eQTL datasets were subset to common significant loci, those where a significant eQTL variant and a significant GWAS variant were within 500kb of each other. After subsetting to shared variants between the two datasets, colocalization was performed using *coloc* under the single causal variant assumption for regions with at least 50 shared variants. Locus plots were made using *locuszoomr*, taking LD information from 1000 Genomes GRCh38 high coverage via *LDlink*. Colocalizations involving eQTLs and association results from our iMGL phenotypes were performed in the same way but subsetting to common significant eQTL variants.

### Fine mapping of TREM2 AD GWAS locus

We used *susie_rss()* from *susieR* for fine mapping of selected GWAS loci, taking LD information 1000 Genomes GRCh38 high coverage via *LDlink* and using the harmonised GWAS summary statistics. Non-default arguments for *susie_rss()* were: n=aggregate number of cases and controls in GWAS; L=1; coverage = 0.60; min_abs_corr = 0.0.

### Testing for enrichment in disease heritability near gene sets

The enrichment of expressed gene sets for common variants associated with disease (GWAS variants) was tested using stratified LDSC^105^ as implemented with CELLECT^106^. Because CELLECT uses as input a continuous representation of cell type expression between zero and one, we re-scaled the t-statistics of differential expression analyses min/max method from 0 (least significant) to 1 (most significant and largest effects), after sorting by either positive or negative sets, and taking those genes with opposite sign as zero. eGene lists were created in the same way by sorting and scaling their combined measure of local false sign rate (lfsr) and posterior mean (pm) estimates (-log10lsfr x pm) from *mashR*.

~~~
Finally, CELLECT - LDSC leverages GWAS variants’ betas and LD structure and the top ranked genes to give a disease enrichment p-value for each treatment.
~~~

### Transcriptome-wide Mendelian Randomization analysis

To estimate the causal effect of gene expression on microglial phenotypes we employed the TWMR method^35^, which estimates the effects of multiple exposures (eGenes) on each outcome (phenotype GWAS result) employing multiple instruments (independent eQTL variants). For each eGene (“focus gene”), per treatment, we selected the most significant eQTL variant and the top colocalizing variant (if present). We then selected eGenes for which any of those variants is an eQTL, and expanded the search for each eGene to other significant eQTL variants (nominal p-value < 10^-8^). We then pruned the variants to only those in low LD (R^2^<0.2, prioritizing those from the focus gene), and eGenes to those that were not highly correlated (R^2^<=0.4 across all variants). We then extracted the betas from the phenotype GWAS and calculated the causal effect of the gene expression on the phenotype solving the following:

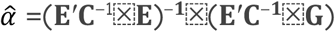

Where E is the *n* × *k* matrix of betas of *n* SNPs on *k* eGenes (as a single cis-eQTL often affects several nearby eGenes in a correlated way). G is the vector of phenotype GWAS beta values (of length *n,* for all variants tested). C is the pairwise LD matrix between the *n* variants, where LD is the R^2^ taken from the EUR cohort from TOPMed^107^. Despite using overlapping samples for exposures and outcomes, strong instrument selection mitigates the inflation of TWMR estimates^108^, and thus we selected genes with F parameter values > 10.

### Association of PRS with microglial phenotype

Full PRS and microglia-specific PRS were related to microglial phenotypes by linear regression. Phenotypes were processed as for the genome-wide association, and a regression model was fitted for all remaining donors with the following formula:

Mean phenotype across pools ∼ PRS + sex + number of pool replicates + mean of minimum donor proportion across pools

Where PRS can be the full PRS including the polygenic component and the APOE component, or the last two on their own. Genotype PCs were not included as covariates as PRS were already adjusted for these.

## Acknowledgements

DRICU iPSC lines were generated pursuant to funding received by Cardiff from the UK Dementia Research Institute (award number UK DRI-3201) through UK DRI Ltd, principally funded by the Medical Research Council, and The Moondance Foundation), and are provided by Cardiff to the Recipient Organisation with the approval of the UK DRI IPMAR Research Group. The authors wish to acknowledge the support of the Cytometry Core Facility and Scientific Operations sequencing facilities at Wellcome Sanger Institute and members of the Bassett and Trynka labs for helpful discussions.

## Code and data availability

The raw flow cytometry, western blotting, processed sequencing data and code to replicate the analyses done in this publication is available on Zenodo (10.5281/zenodo.16684932). The donor deconvolution R package *poodleR* is available on GitHub.

All raw sequencing data is available under study EGAS00001004854 (scRNA- seq) and EGAS00001005143 (DNA-seq), comprising scRNA-seq and WGS data from all pools. Donor and line genotypes are available at www.hipsci.org and the genotype of SH5Y5Y is available at https://systemsbiology.uni.lu/shsy5y/.

## Contributions

A.B., G.T. and S.A.C. conceived the study. S.W., Y.C., J.S. and J.M. performed the laboratory experiments. M.P-A., D.G-P. and P.K. performed the bioinformatics data analysis. S.W. performed and analysed the CRISPR screen. G.T., A.B., S.A.C., K.A. N.P. and D.W. supervised the analysis. M.P-A., S.W., G.T., A.B., D.G-P. and J.C. wrote the manuscript. All authors interpreted the results and provided critical comments on the manuscript.

