## Supplemental Figures for "Integrated QTL mapping and CRISPR screening in pooled iPSC-derived microglia reveals genetic drivers of neurodegenerative risk"

Consisting of Supplementary figures 1-23

#### Supplementary Figure 1

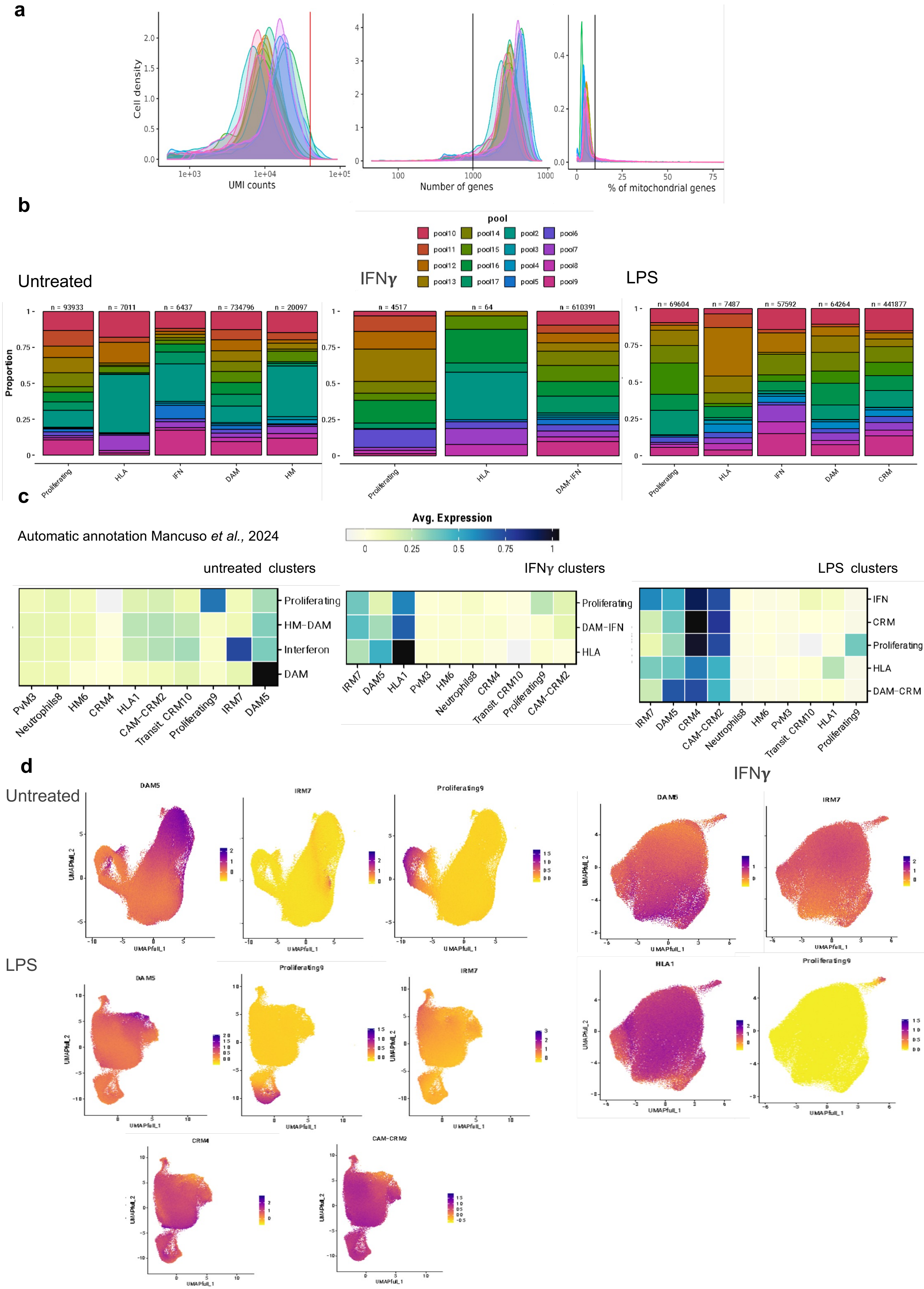

**Supplementary Figure 1:** a) Histograms of cell density of UMI counts, number of genes and % of mitochondrial genes per pool. b) Cell proportions (y axis) per pool (color) and named cluster (x axis), per treatment. Total number of cells is shown over each cluster. c) Results of automatic cell annotation per named cluster, using as reference the annotated cells from Mancuso *et al.* 2024. d) Results of automatic cell annotation per cell (from Mancuso *et al.* 2024) projected within each treatment's UMAP.

#### Supplementary Figure 2

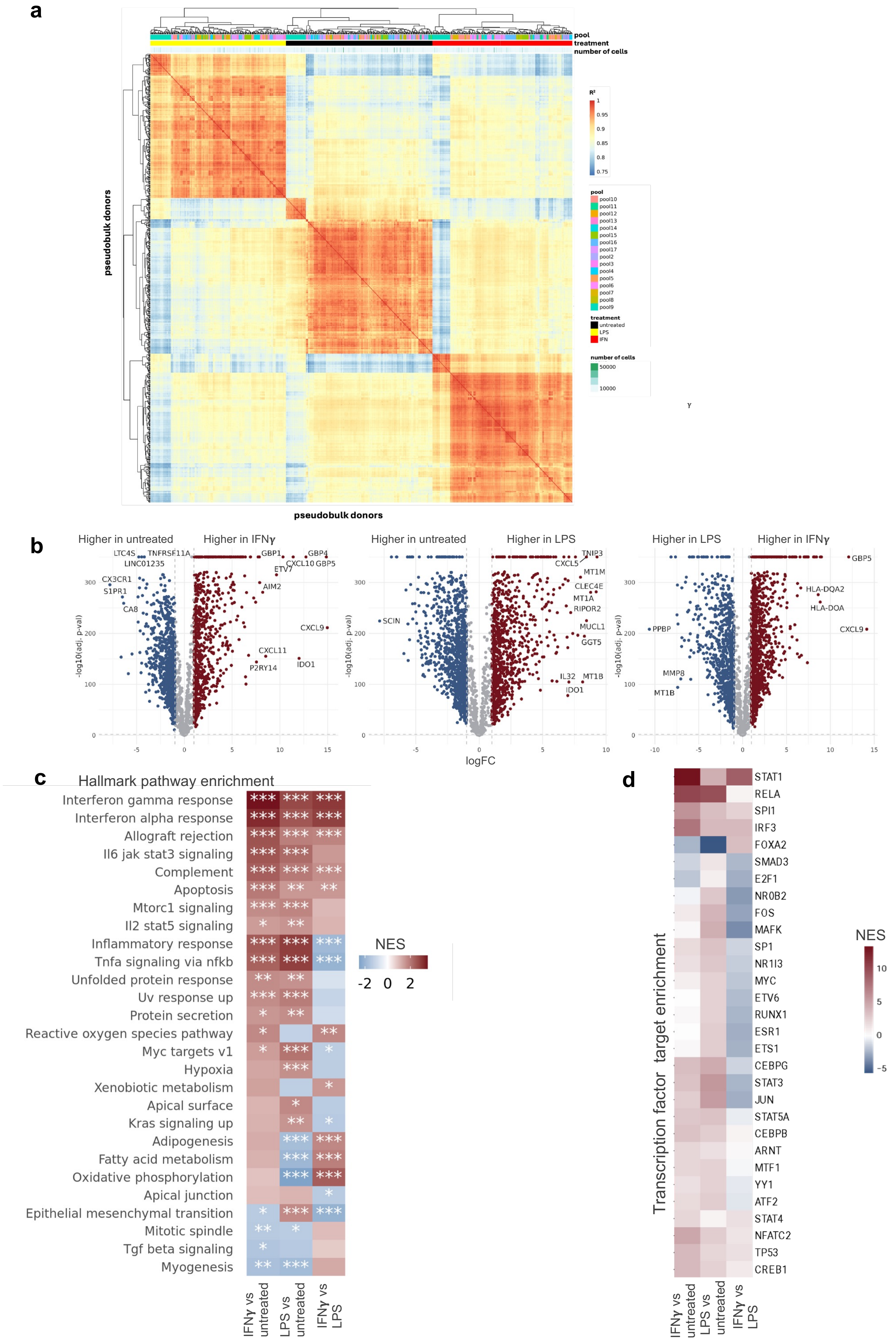

**Supplementary Figure 2:** a) Heatmap of the Pearson $R^2$ correlation of pseudobulked RNA-seq UMI counts, aggregated per donor, treatment and pool. Results have been clustered row and column-wise, with distance shown as a dendrogram on left and upper side, with coloured bars showing metadata for pool, treatment, and number of cells in each aggregated sample. b) Volcano plots of the pairwise differential expression comparisons, with false discovery rate ($-\log10_{FDR}$) on the y axis and fold change (log2FC) on the x axis. Dots with FDR < 0.05 and |log2FC| > 1 are colored by direction of effect. c) Heatmap of Hallmark pathway enrichment (using Gene Set Enrichment Analysis [GSEA]) results for signed and centered t-statistics of the pairwise differential expression comparisons, colored by normalised enrichment score (NES). BH-adjusted p-values are reported: * < 0.05, ** < 0.01, *** < 0.001. d) Heatmap of the top 30 enriched transcription factor (TF) targets and significant Progeny pathways of the pairwise differential expression comparisons results ranked as in c).

#### Supplementary Figure 3

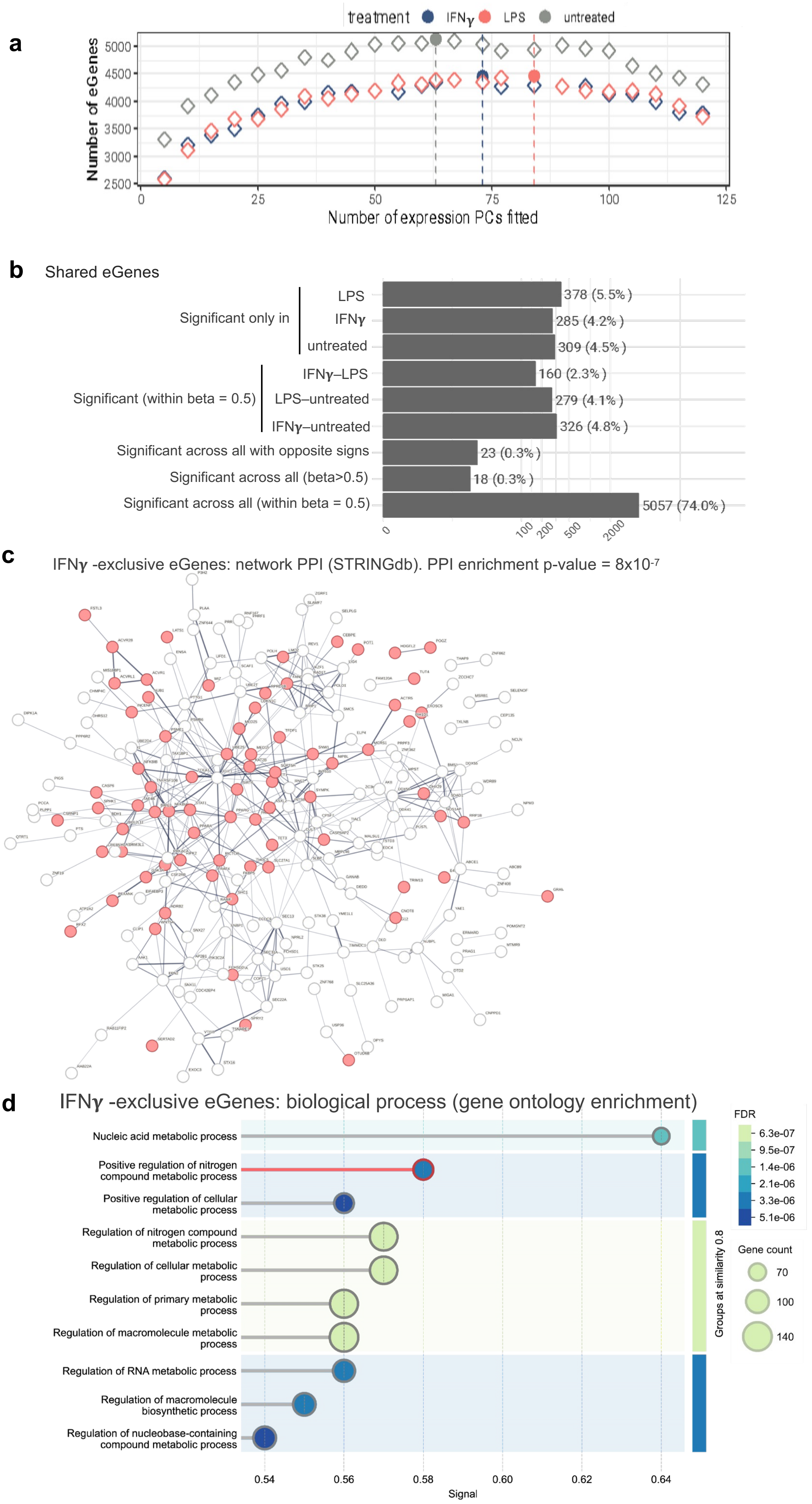

**Supplementary Figure 3:** a) Number of eGenes (y axis) per number of expression Principal Components (PCs) fitted, coloured by treatment. Filled dots represent final number if PCs fitted for that treatment. b) Barplots of percentage (in parentheses) and number of eGenes per specificity category, as measured by *mashR*. c) Network of Protein-Protein Interactions (PPI) in STRINGdb for the IFN$\boldsymbol{\gamma}$ -exclusive eGenes. In red: positive regulators of nitrogen compound metabolic process. d) Lollipop plot of gene ontology enrichment of biological process terms for IFN$\boldsymbol{\gamma}$ -exclusive eGenes. Highlighted in red: the term with network shown in c). Terms are grouped by similarity. Size of dots represents the gene set size, and the color the enrichment FDR.

**Supplementary Figure 4**

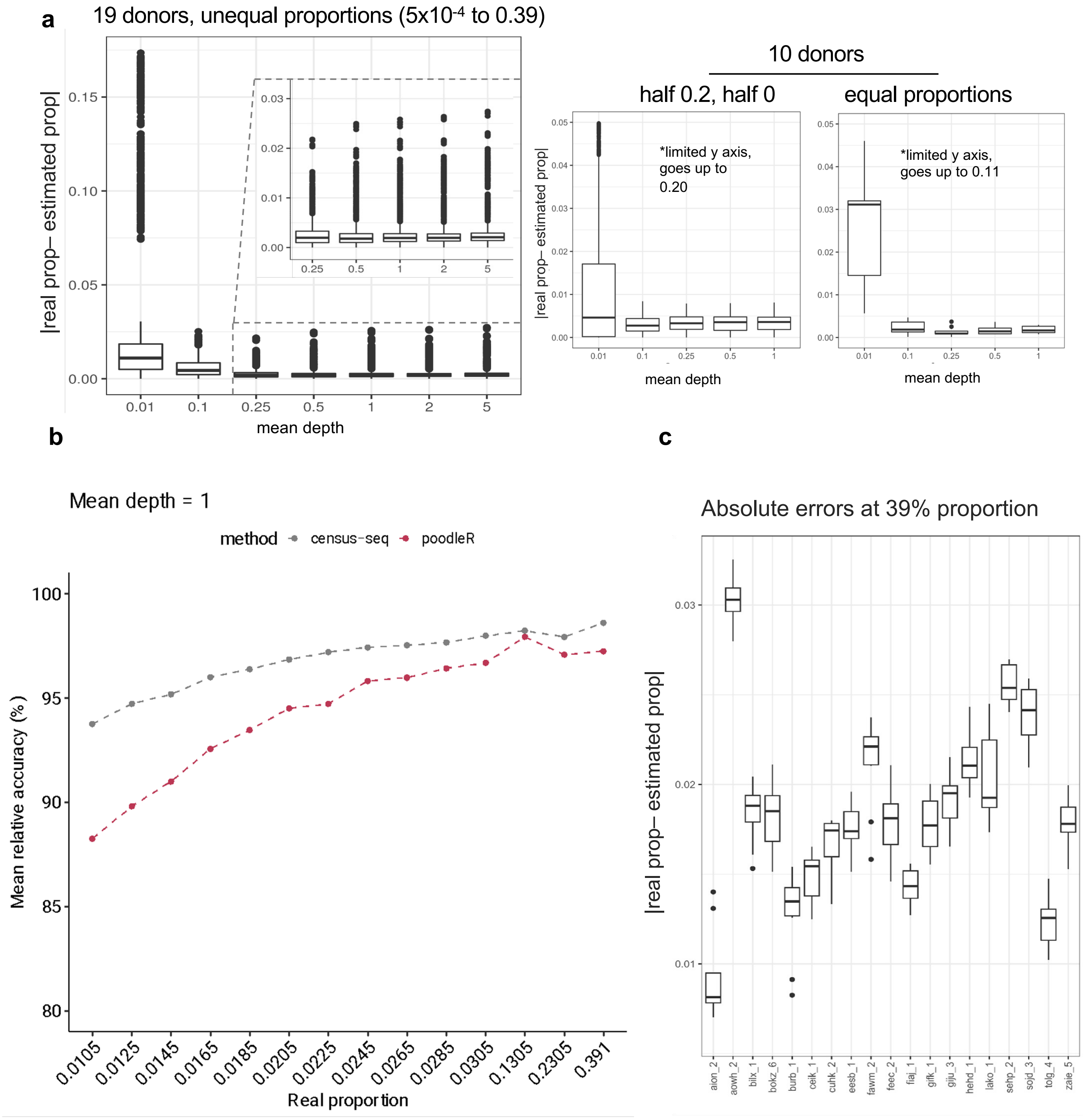

**Supplementary Figure 4:** PoodleR accuracy results on simulated WGS pools a) Absolute difference between real and estimated donor proportions (y axis) per mean depth (x axis). Synthetic pools comprise 19 donors at unequal donor proportions (left), or 10 donors where half are at equal proportions and half are absent (middle), or 10 donors at equal proportions (right). b) Mean relative accuracy (y axis) comparison for a pool of 19 donors at a range of proportions (x axis) between poodleR and census-seq. c) In a pool of 19 donor at unequal proportions, the absolute difference between real and estimated proportions (y axis) is shown for a range of donors (x axis) in repeats where they present 39% donor proportions. Aggregated over 100 synthetic pool replicates.

### Supplementary Figure 5

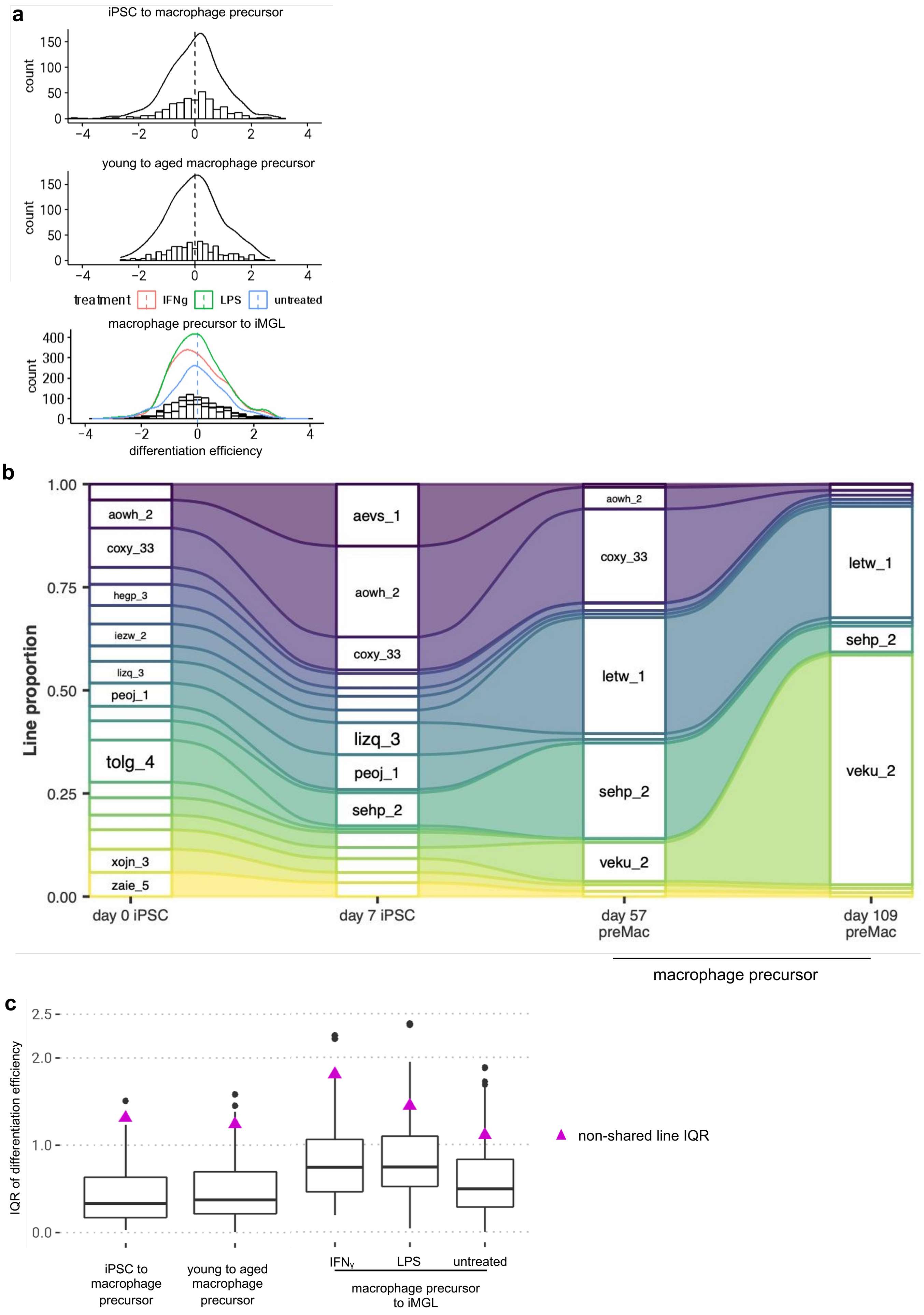

**Supplementary Figure 5:** a) Frequency distribution of scaled differentiation efficiency at the three stages measured (from top to bottom): iPSC to macrophage precursors, young to aging macrophage precursors, and macrophage precursors to iMGL (the latter per treatment). b) Alluvial plot representing changing donor proportions with longer iPSC culture and aging precursors. c) Distribution of interquartile ranges (IQR) of differentiation efficiency between shared lines (boxplots) and non-shared lines (average represented by magenta triangle), per comparison.

### Supplementary Figure 6

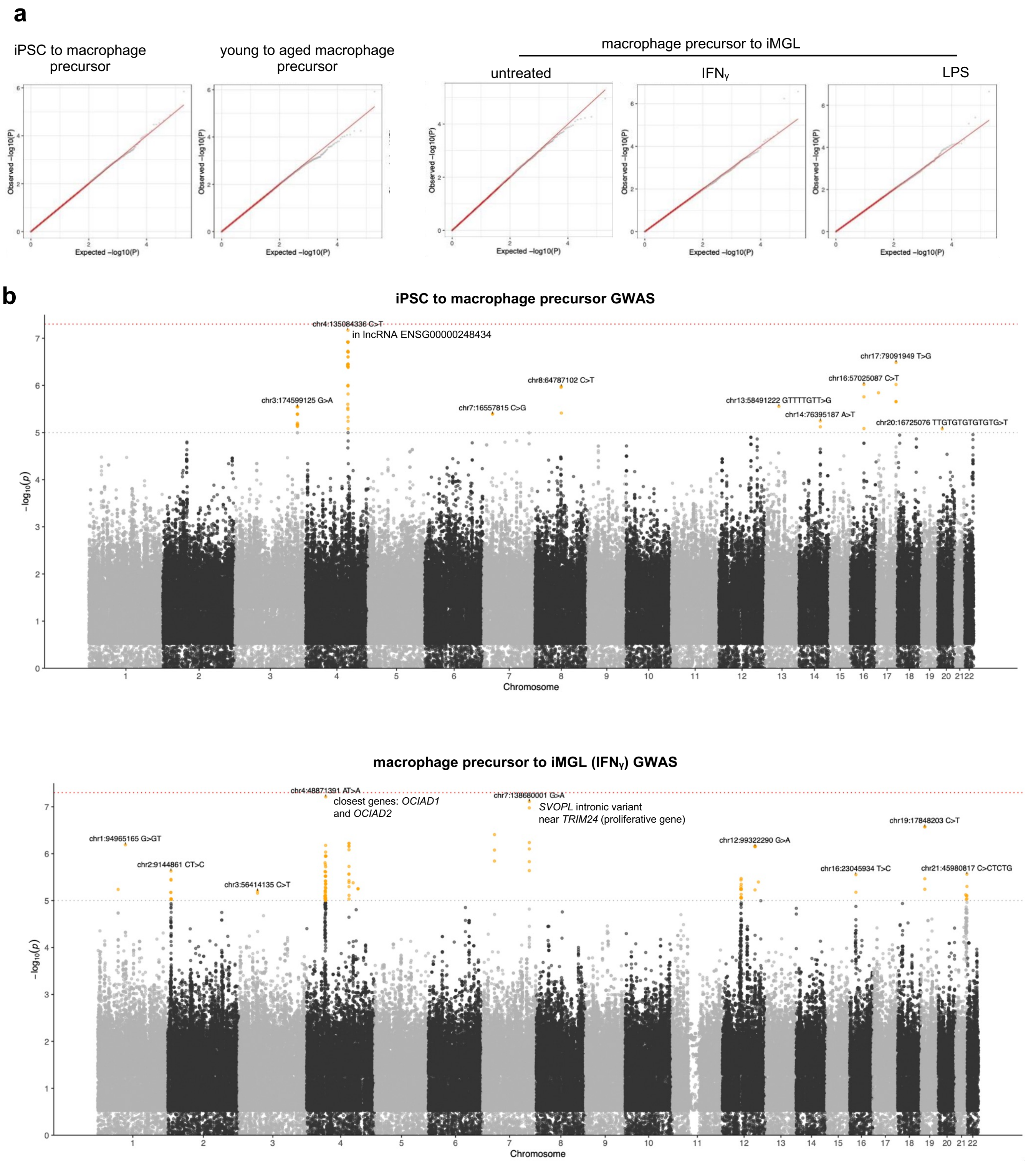

**Supplementary Figure 6:** a) Quantile-quantile plot of differentiation efficiency GWAS, per comparison. b) Manhattan plots of results from GWAS of differentiation efficiency from iPSC to macrophage precursors, and macrophage precursors to iMGL (IFNɣ treatment). Lead variants and nearby genes are annotated.

### Supplementary Figure 7

**a**

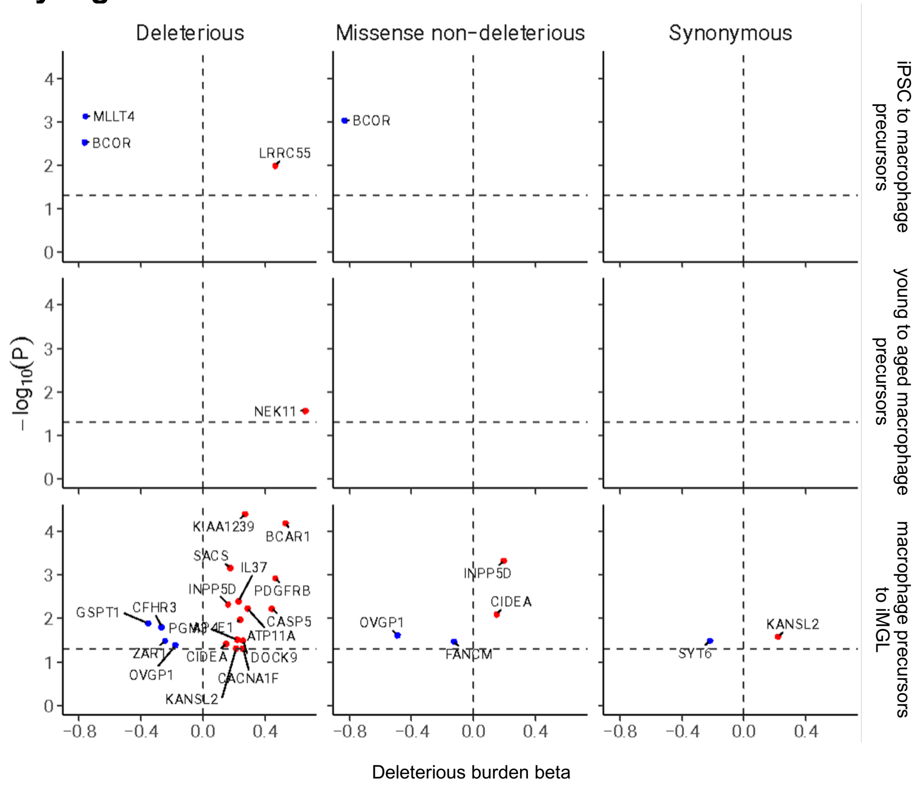


**b**

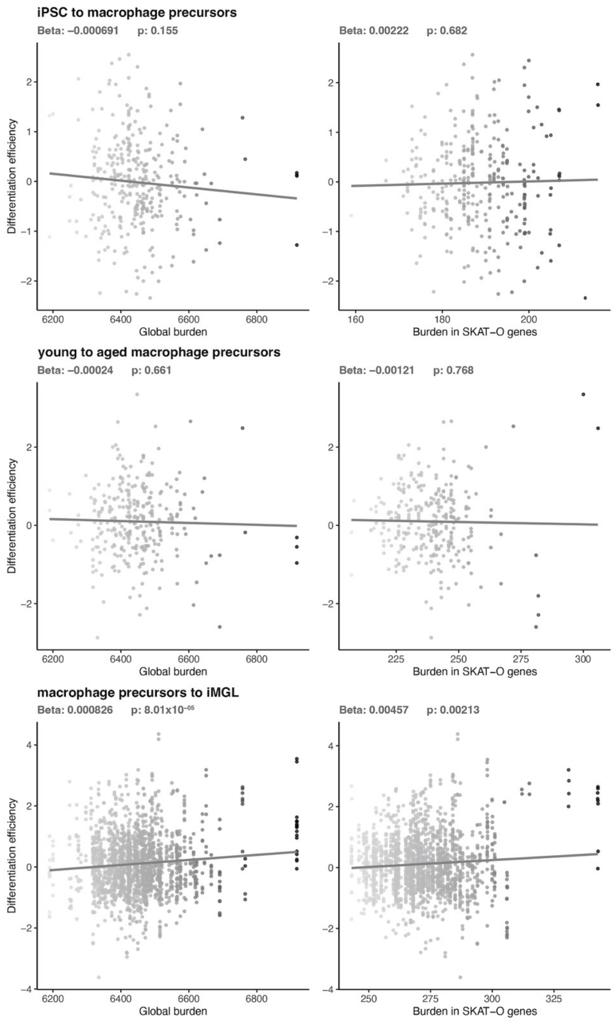


**c**

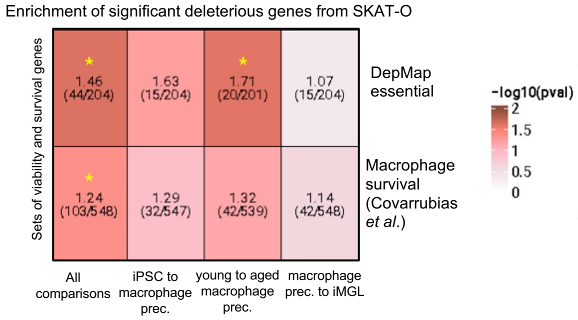

**Supplementary Figure 7:** a) Significant results for gene burden tests of deleterious, missense non-deleterious and synonymous variants on differentiation efficiency, per stage change. –log10(p-value) on the y axis, and burden test beta estimates on the x axis. b) Scatter plot of burden of deleterious variants (x axis) vs differentiation efficiency (y axis), per stage change (top to bottom), transcriptome-wide and for significant genes as tested by SKAT-O (left to right). Represented is the slope line, the beta and the p-value of the linear regression. c) Enrichment results of significant SKAT-O results for deleterious variants per all and each stage comparison (left to right), for two sets of viability and survival genes (top to bottom). Color denotes –log10(p-value) of GSEA, the top value in each box denotes the normalized enrichment score, and within parenthesis is shown the found genes / total tested genes per group. * denotes p-value < 0.05. In all subfigures macrophage precursor to iMGL results are reported for the three treatments.

### Supplementary Figure 8

**a**

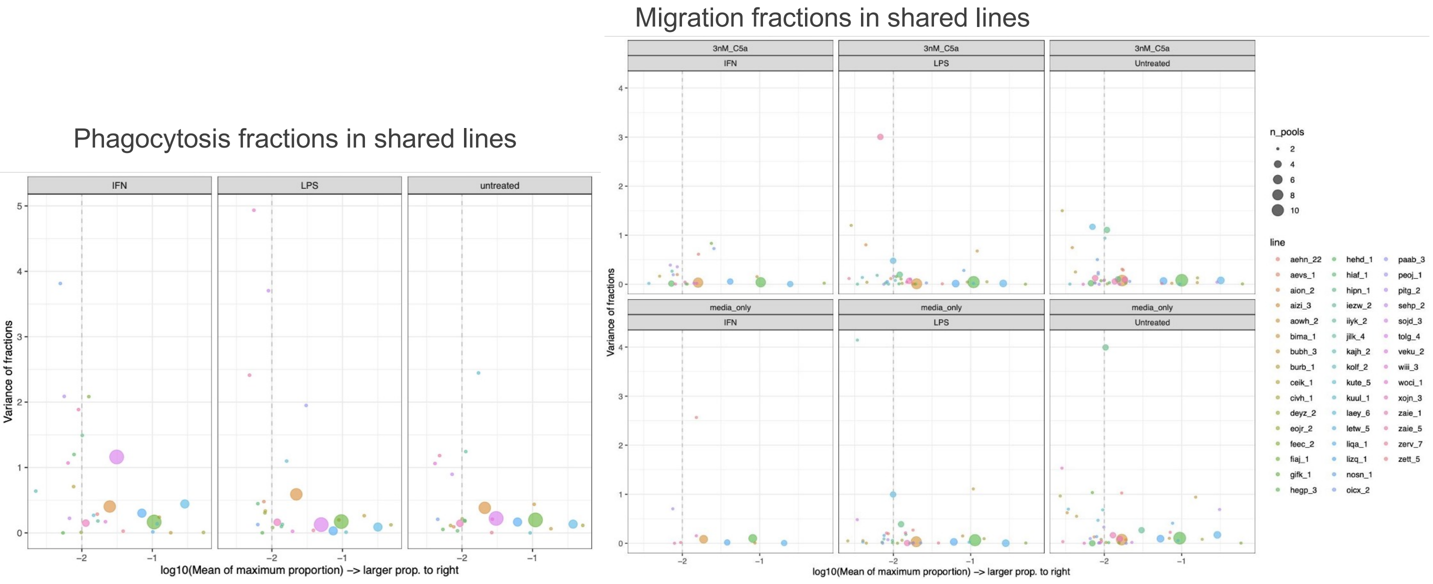


**b**

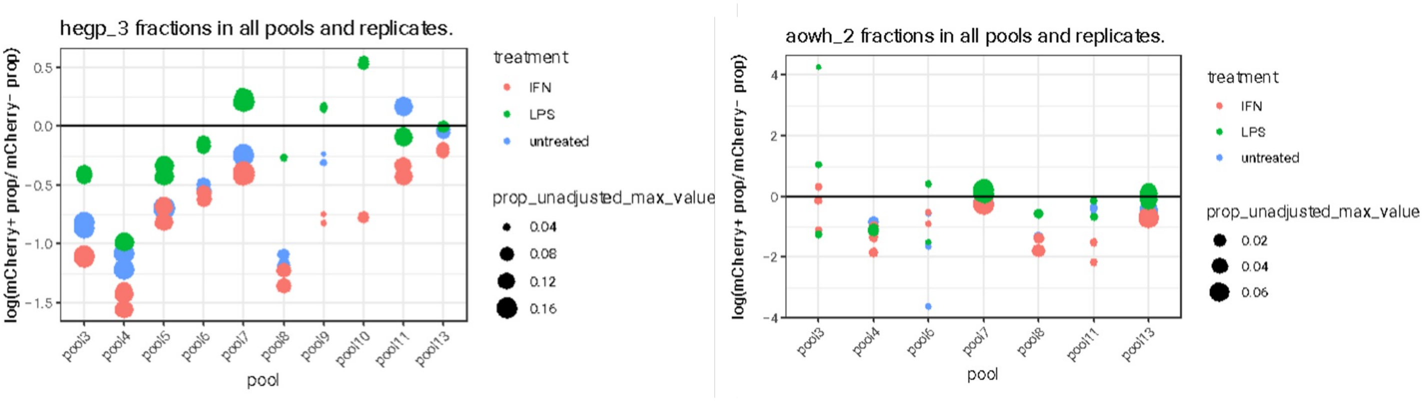

**Supplementary Figure 8:** a) Variability of line proportions across replicates, treatments and pools for lines shared in at least 2 pools in phagocytosis (left) and migration (right) phenotypes. y axis represents the variance of the scaled phenotype. x axis represents the $\log_{10}$-adjusted mean of the largest proportion of the two measured in the fraction. Area of the circle represents in how many pools the line is repeated. Vertical dashed line marks the 1% line proportion threshold. b) Estimate of variability of two lines: hegp_3 (left) and aowh_2 (right). y axis represents the variance of the scaled phenotype. x axis represents the pool. The area represents the maximum proportion of the two measured in the fraction.

#### Supplementary Figure 9

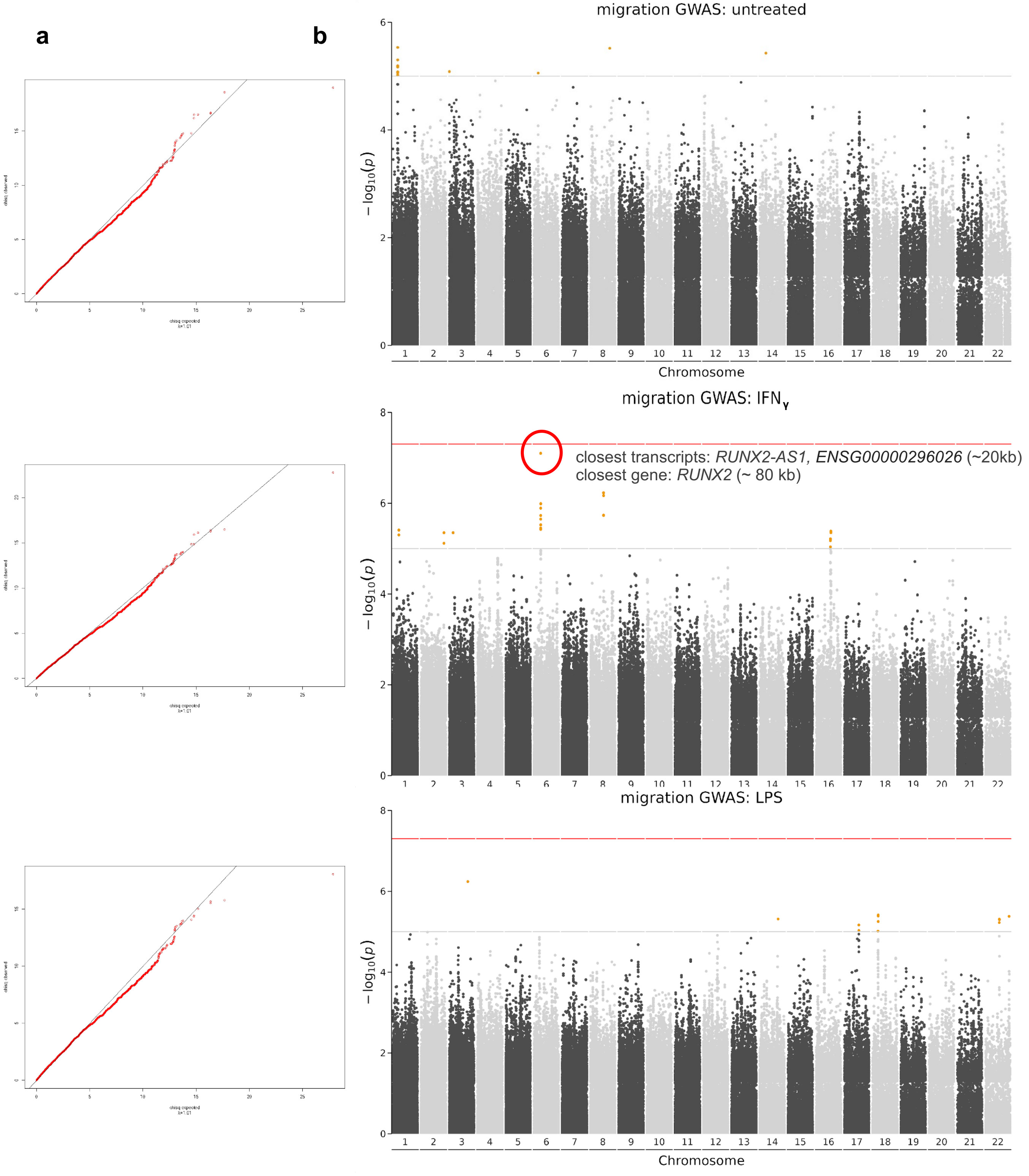

**Supplementary Figure 9:** a) Quantile-quantile plot of migration GWAS, per treatment. b) Manhattan plots of results. Red horizontal line indicates genome-wide significant threshold. Grey horizontal line indicates suggestive significance threshold.

#### Supplementary Figure 10

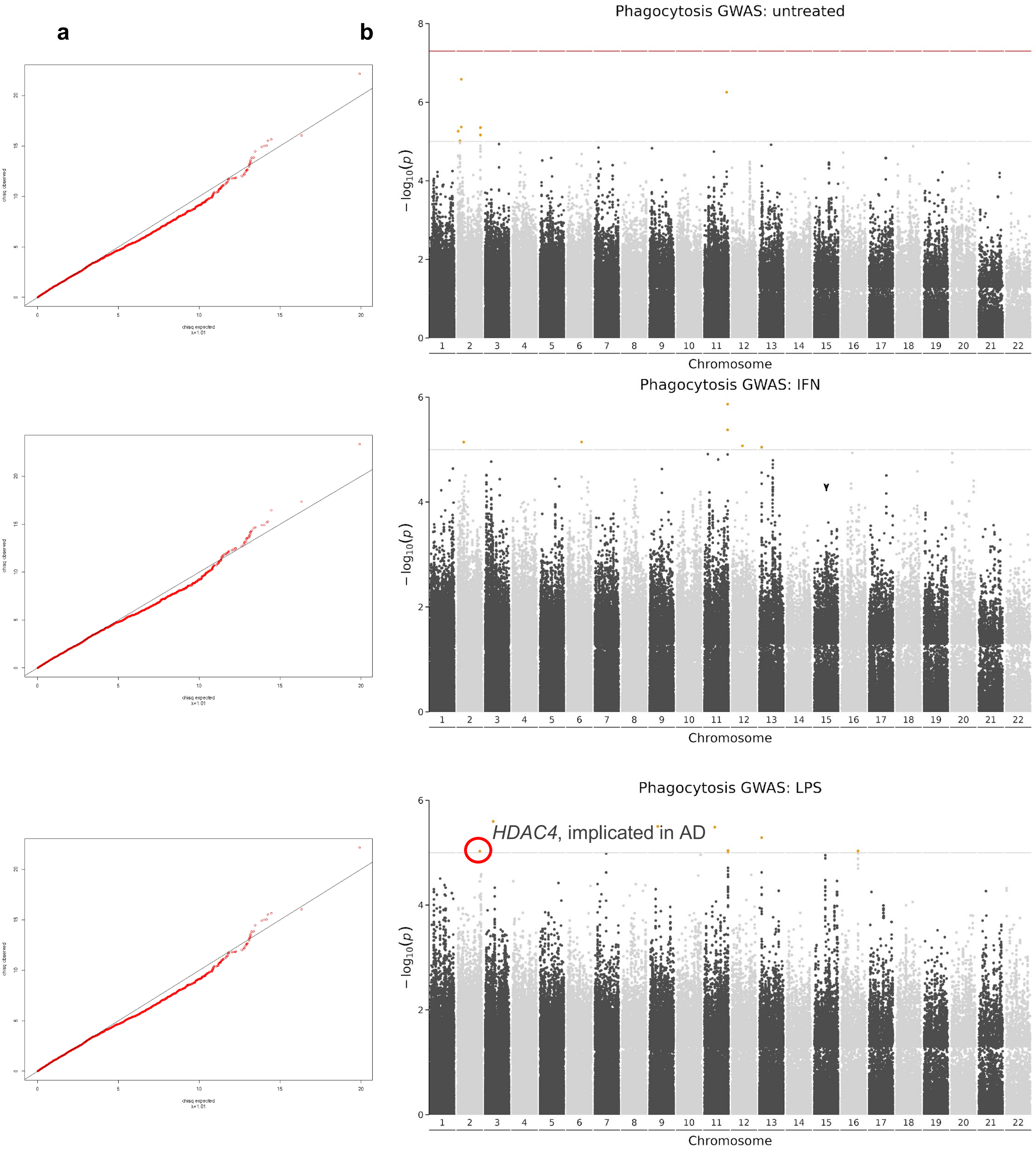

**Supplementary Figure 10:** a) Quantile-quantile plot of phagocytosis GWAS, per treatment. b) Manhattan plots of results. Red horizontal line indicates genome-wide significant threshold. Grey horizontal line indicates suggestive significance threshold.

#### Supplementary Figure 11

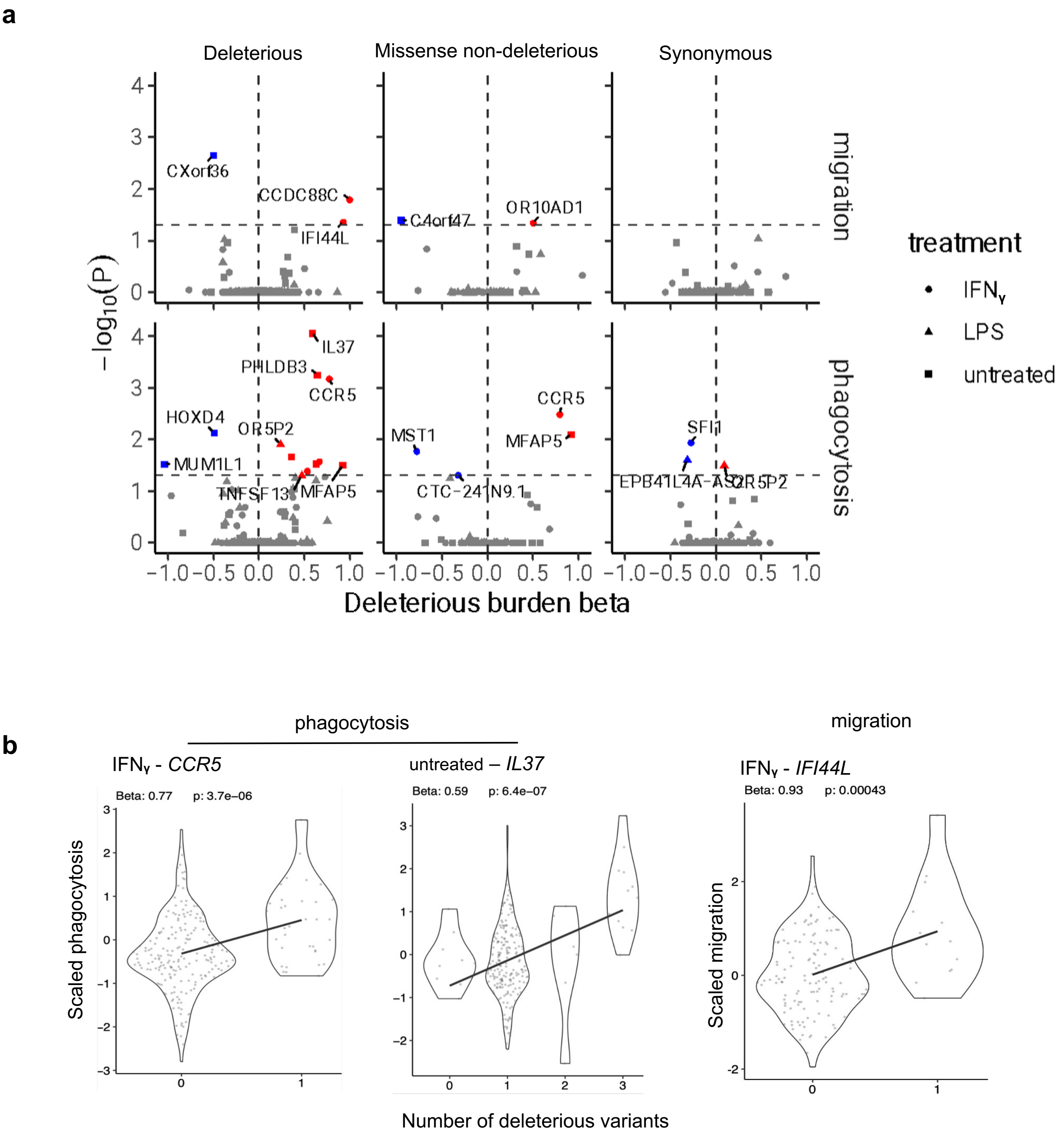

**Supplementary Figure 11:** a) Significant results for gene burden tests of deleterious, missense non-deleterious and synonymous variants on phagocytosis and migration, per stage change. –log10(p-value) on the y axis, and burden test beta estimates on the x axis. Shapes represent each treatment. b) Burden tests regression estimates of highlighted genes, with scatter and violin plot of phenotype estimates (y axis) vs number of deleterious variants (x axis). Depicting regression line, regression estimate (beta) and its p-value.

#### Supplementary Figure 12

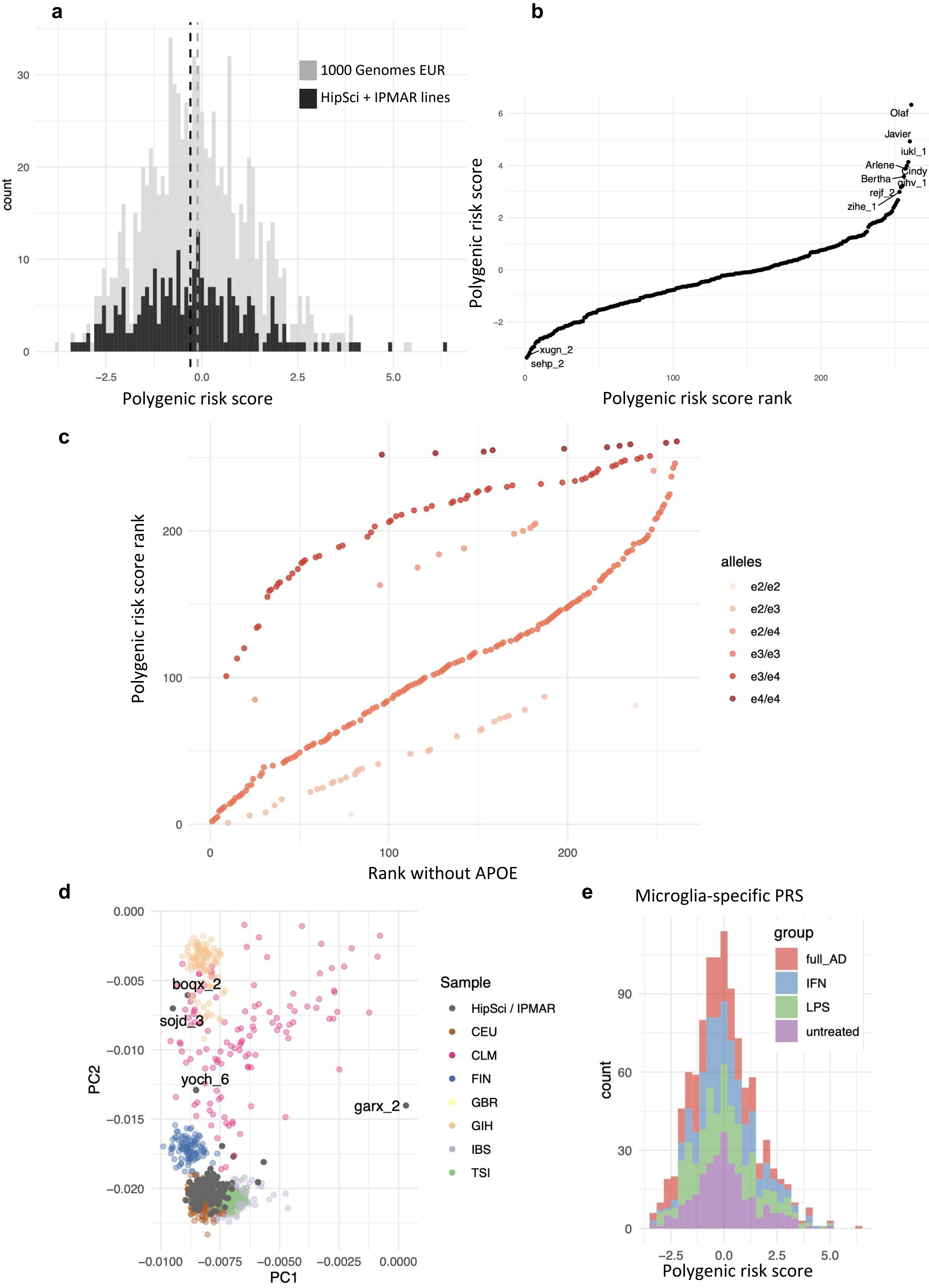

**Supplementary Figure 12:** a) Distribution of polygenic risk scores (x axis) from 1000G EUR subset and the HipSci and IPMAR donors used in this study. b) Ranks (x axis) of polygenic risk scores (y axis) from each HipSci and IPMAR donor. c) Ranks of polygenic risk scores with (y axis) and without (x axis) the APOE component. Different combinations of APOE alleles were colored. d) PCA plot of HipSci and IPMAR lines (in dark grey) overlaid on top of 1000G population subsets. Outlier donors (clustering near non-European populations) were labelled. CEU, Utah residents with Northern and Western European ancestry ; CLM, Colombian in Medellin ; FIN, Finnish in Finland ; GBR, British in England and Scotland ; GIH, Gujarati Indian in Houston, TX; IBS, Iberian populations in Spain; TSI, Toscani in Italy. e) Histogram of microglia-specific PRS distribution, calculated from variants near colocalizing genes per treatment (full_AD) indicates the regular PRS using the full AD GWAS results.

#### Supplementary Figure 13

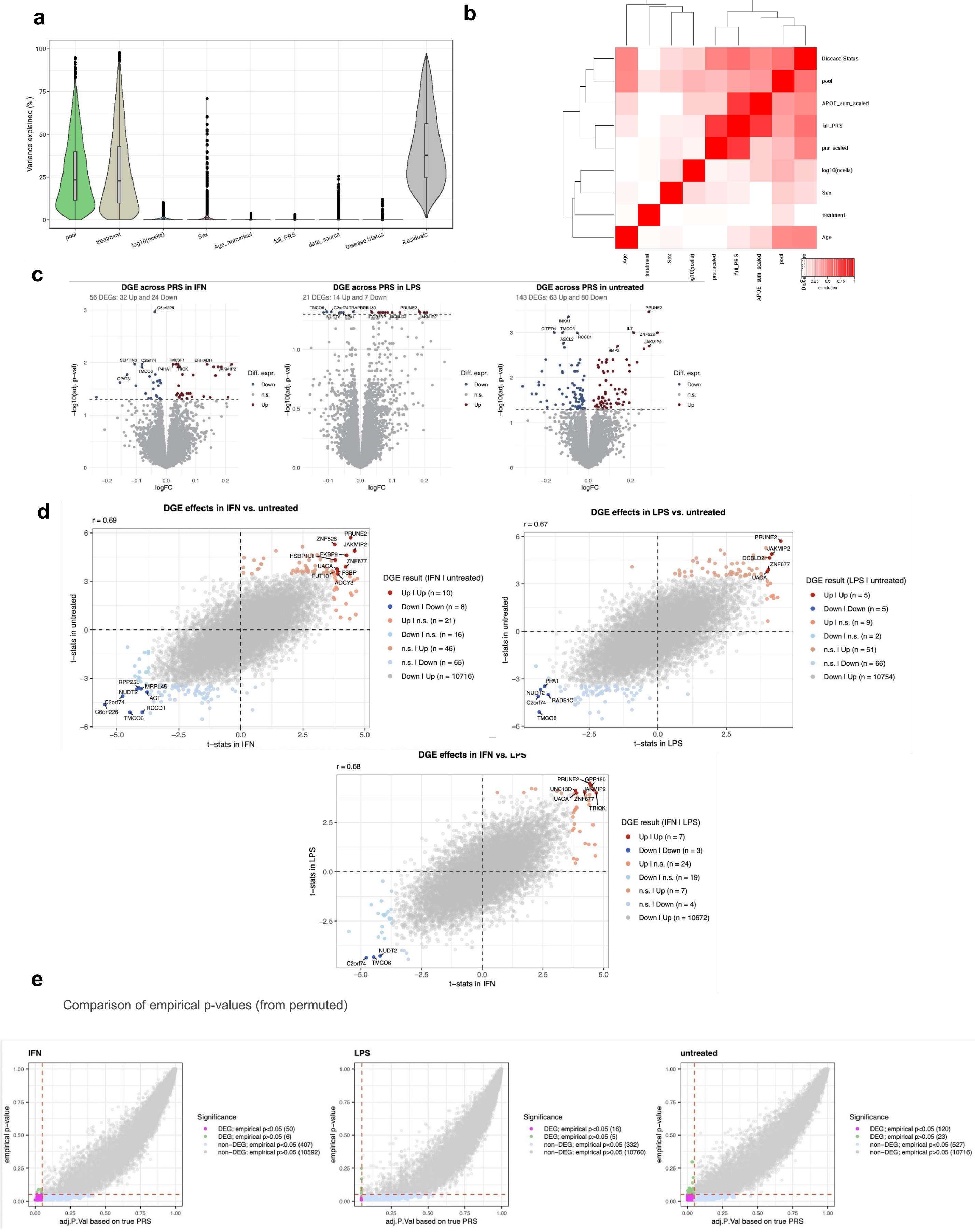

**Supplementary Figure 13:** a) Variance partition analysis of the pseudobulked expression (by donor, pool and treatment) showing the percentage of variance explained (y axis) per metadata category. b) Canonical correlation analysis results of the metadata. c) Volcano plots of differential gene expression across AD polygenic risk score (PRS) values: coloured dots represent genes with FDR < 5% and abs(log2FC) > 1. d) Scatterplot of differential expression t-statistic across PRS, for the LPS and IFN**γ** treatments and for the IFN**γ** - treated and untreated. e) Comparison between regular p-values from limma (x axis) and empirical p-values (y axis).

#### Supplementary Figure 14

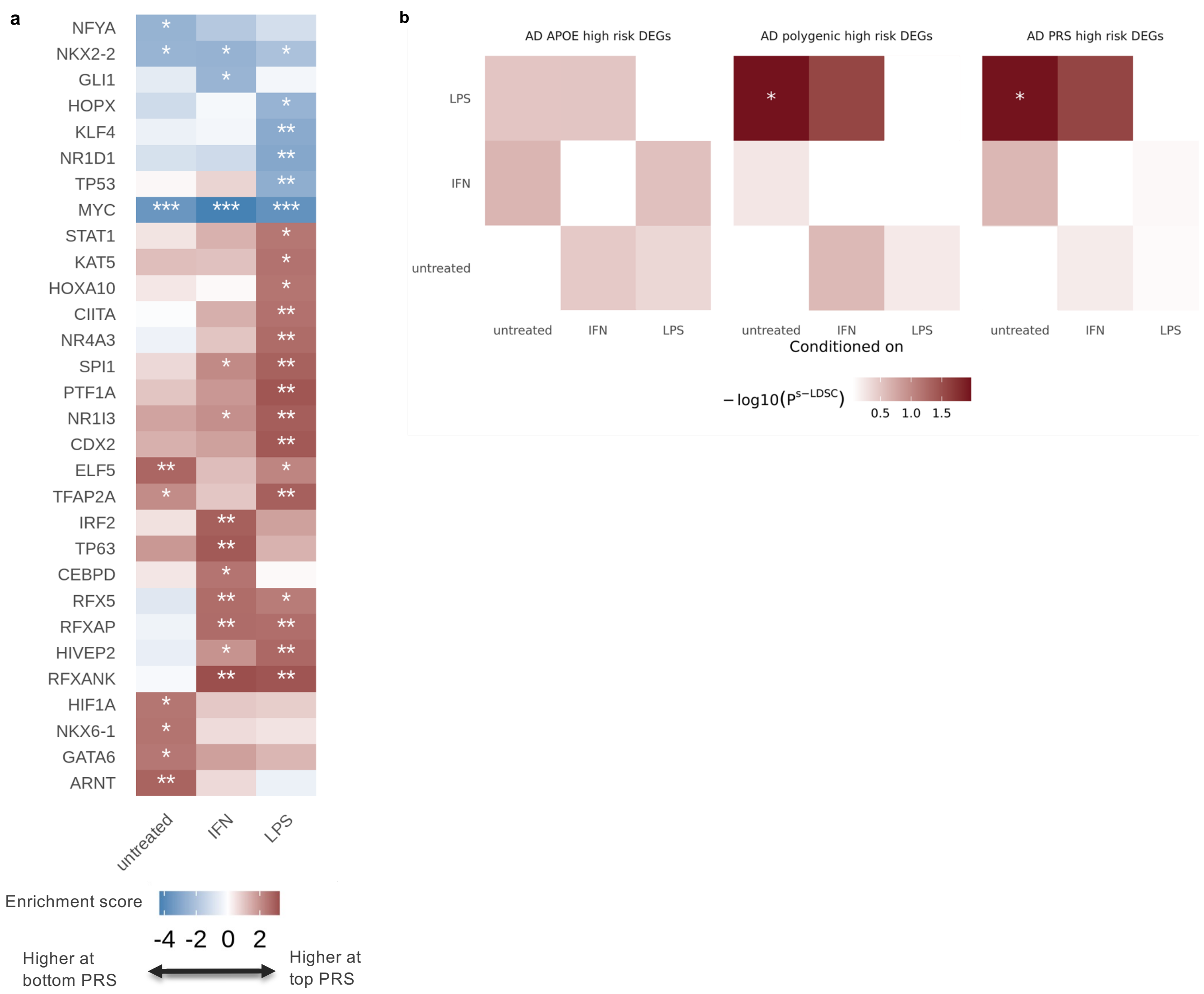

**Supplementary Figure 14:** a) Activity inference scores for PRS DEA results, showing the top 30 enriched transcription factor (TF) targets. Positive scores indicate upregulated TF targets at high PRS. b) Stratified LDSC heritability enrichments for AD GWAS signals within high PRS risk genes (for APOE-only PRS, polygenic PRS or full PRS) for various treatments (y axis), conditioned on the treatments indicated on x-axis. Significance level for the Bonferroni adjusted p-value * < 0.05.

### Supplementary Figure 15

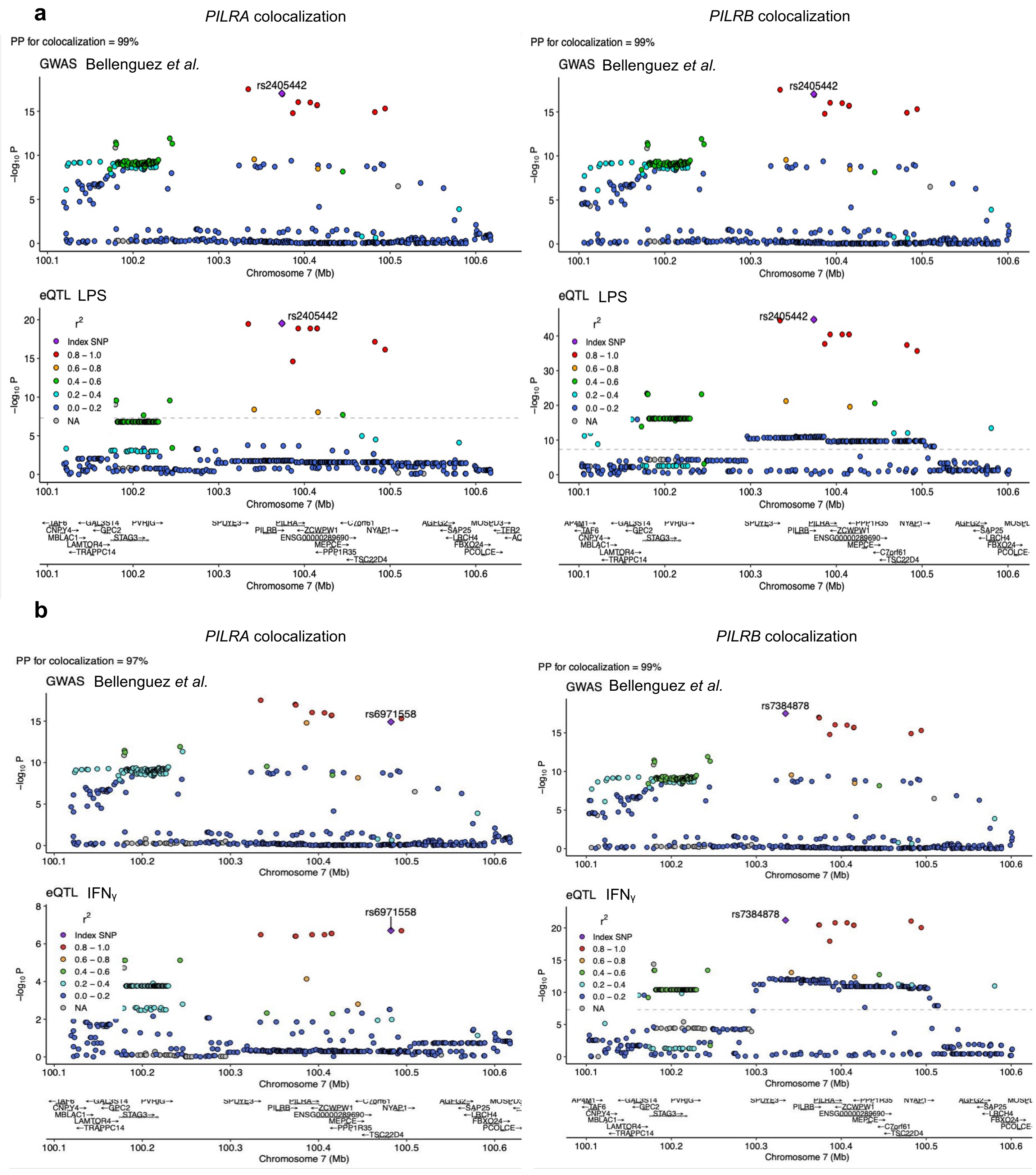

**Supplementary Figure 15:** a) Colocalization plots for *PILRA* (left) and *PILRB* (right) loci between the AD GWAS results from Bellenguez *et al.* and our eQTL in LPS-treated iMGL. b) Colocalization plots for *PILRA* (left) and *PILRB* (right) loci between the AD GWAS results from Bellenguez *et al.* and our eQTL in IFNɣ-treated iMGL.

### Supplementary Figure 16

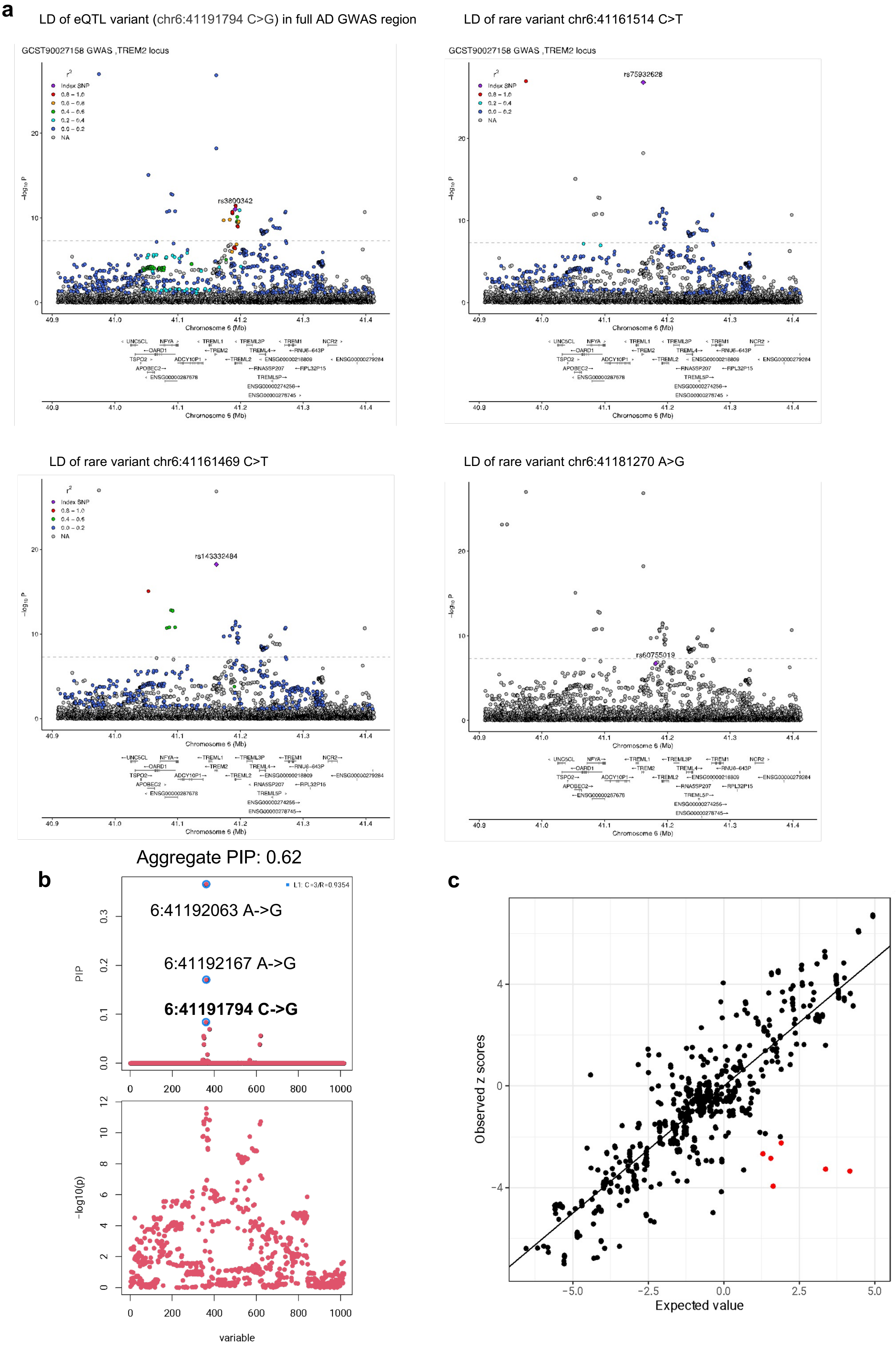

**Supplementary Figure 16:** a) LD (1000 Genomes high coverage) in the full set of tested variants from Bellenguez *et al.* AD GWAS for the *TREM2* locus. Index SNPs are: top left, our lead eQTL variant; the rest are the three independent variants identified by conditional analysis in the GWAS. b) Top: Credible set of AD GWAS variants (subset to shared with eQTL variants) at *TREM2* locus. Highlighted the eQTL lead variant. x-axis is SNP position, y-axis is the marginal posterior inclusion probability. Legend shows the average linkage disequilibrium R between the three variants. Bottom: AD GWAS association p-values for the variants depicted above. c) Correlations of observed vs expected z scores: significant deviations shown in red.

### Supplementary Figure 17

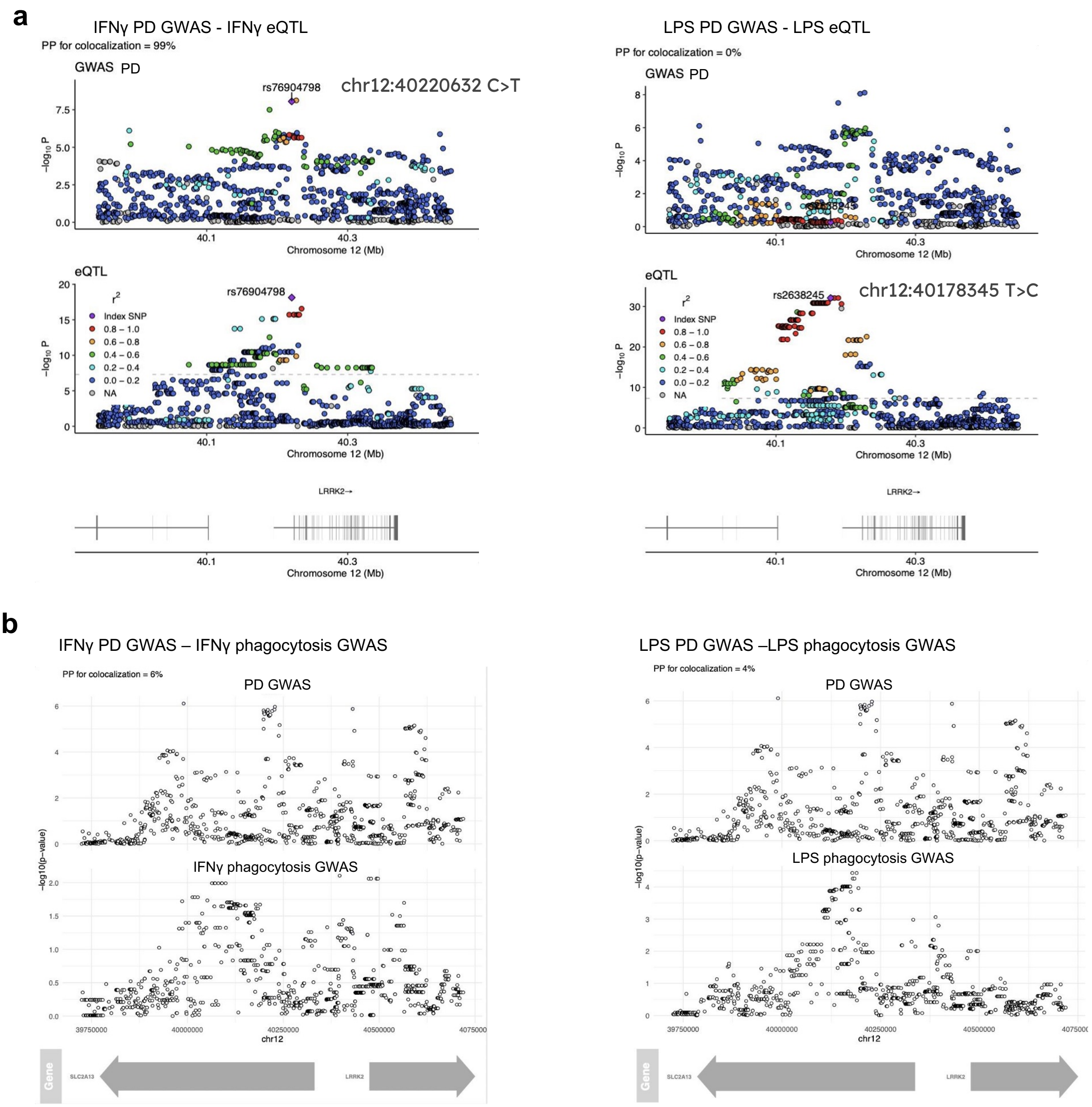

**Supplementary Figure 17:** *LRRK2* locus colocalization figures. a) Colocalization between PD GWAS from Nalls *et al.,* and iMGL eQTL in IFNγ-treated (left) and LPS-treated cells (right). b) Colocalization between PD GWAS and iMGL phagocytosis GWAS in IFNγ-treated (left) and LPS-treated cells (right).

### Supplementary Figure 18

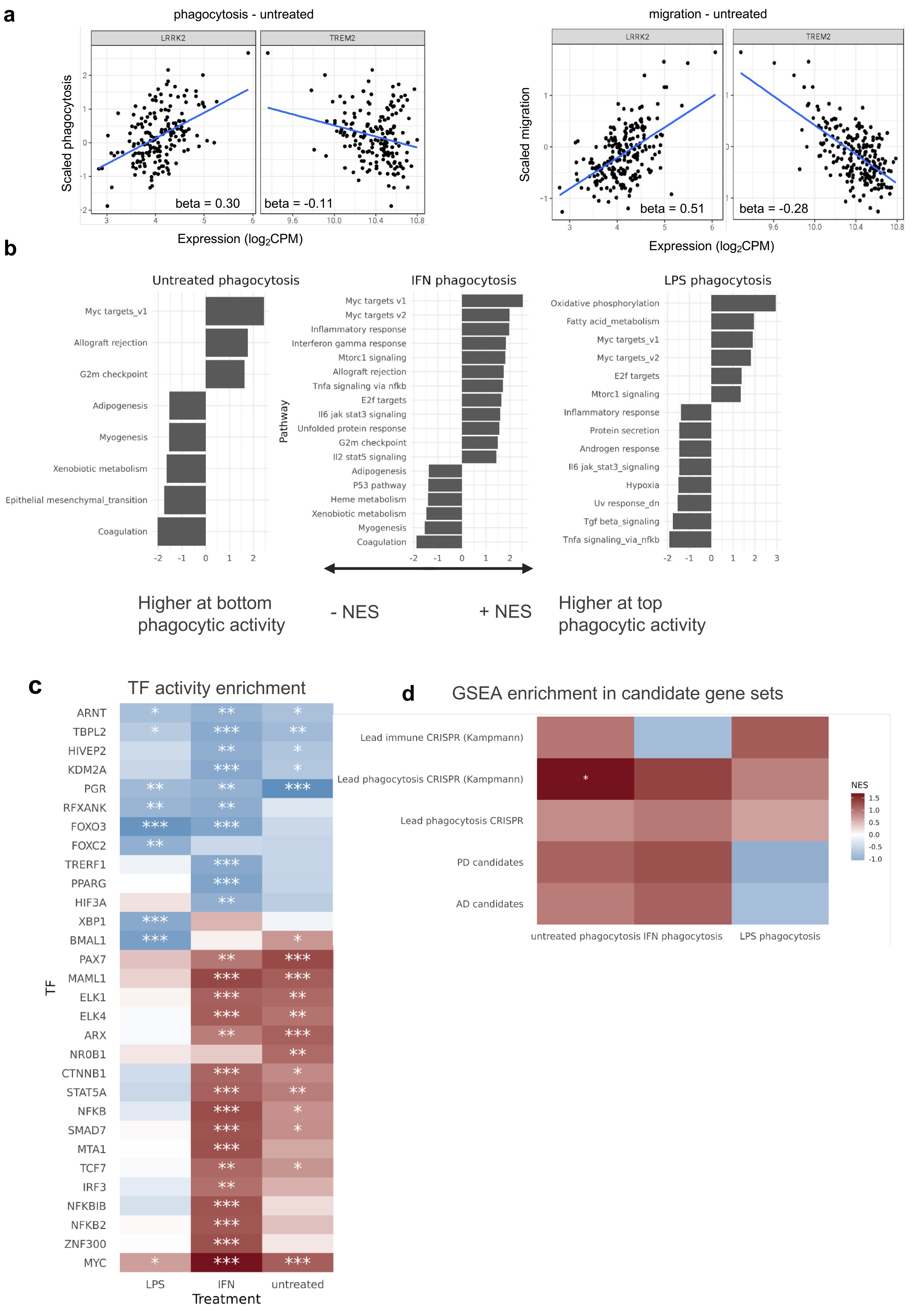

**Supplementary Figure 18:** a) Relationship between gene expression and phenotypes for *LRRK2* and *TREM2*, in the phagocytosis (left) and migration (right) phenotypes for untreated iMGL. b) Reactome hallmark pathway enrichments using GSEA for differential expression results across phagocytic activity. c) Activity inference scores for differential expression results across phagocytic activity, showing the top 30 enriched transcription factor (TF) targets. Positive scores indicate upregulated TF targets at high phagocytosis. d) Enrichments of candidate genes using GSEA for differential expression results across phagocytic activity. All phagocytosis DEA results have been ranked by the signed t-statistic for GSEA.

**Supplementary Figure 19**

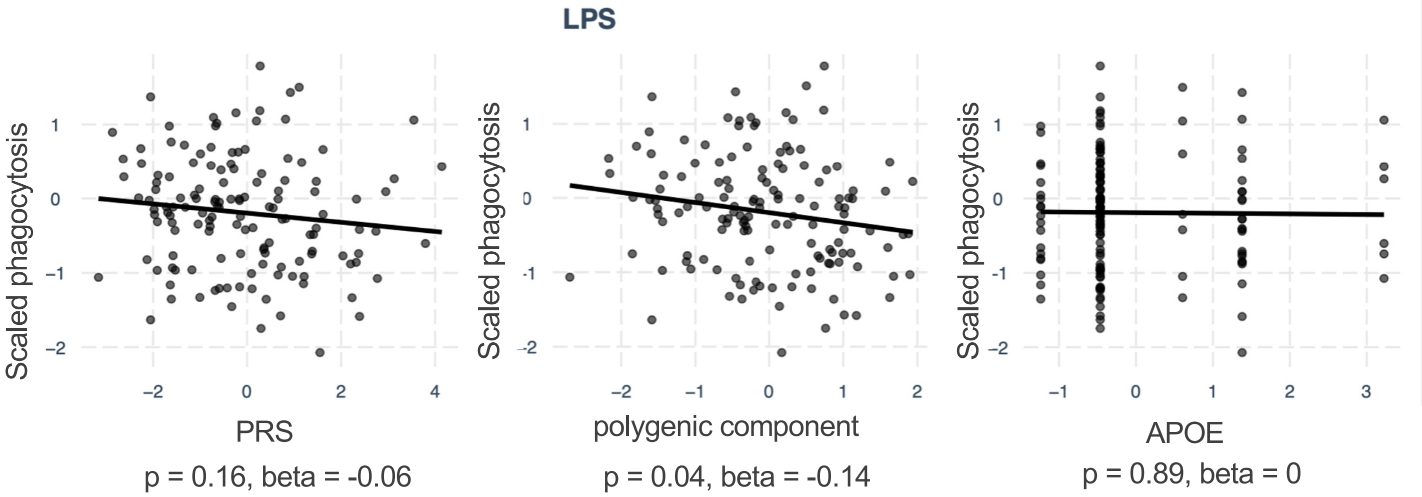

**Supplementary Figure 19:** Scatterplots with regression line and estimate betas and p-values of linear regression between AD eQTL-PRS (x axis) and phagocytosis (y axis) in LPS-treated iMGL. Left to right indicate the full eQTL-PRS, the polygenic component of the eQTL-PRS only, and the APOE component.

**Supplementary Figure 20**

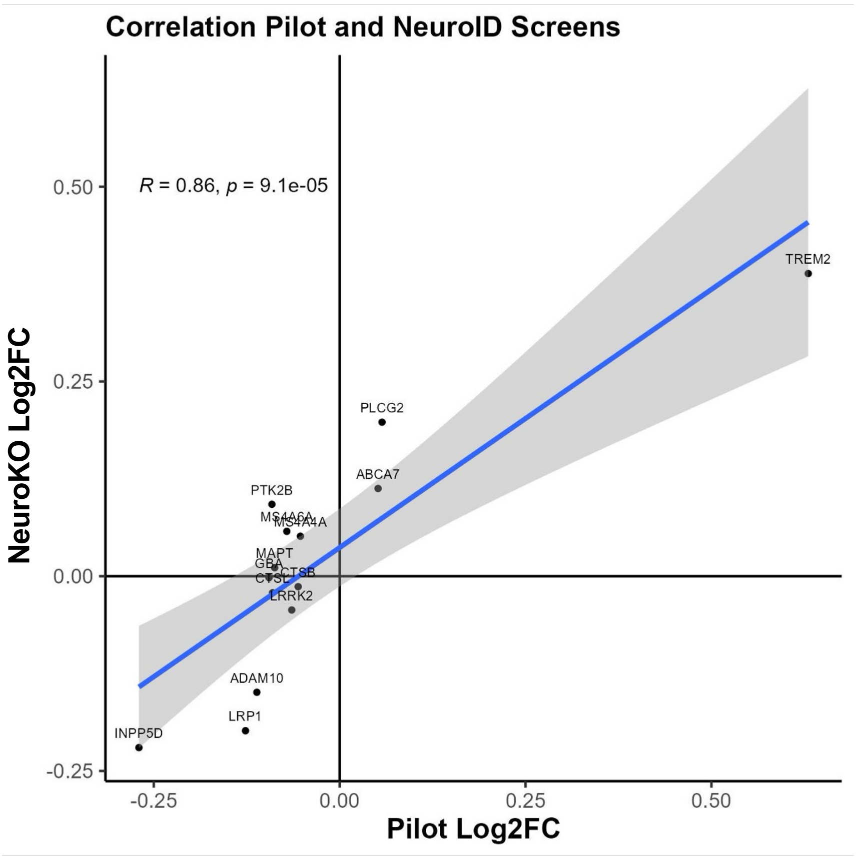

**Supplementary Figure 20:** Correlation of the Log2FC between the overlapping genetic targets of the NeuroKO screen (y axis) and the pilot screen (x axis). Dark grey shaded area indicates 95% confidence interval.

#### Supplementary Figure 21

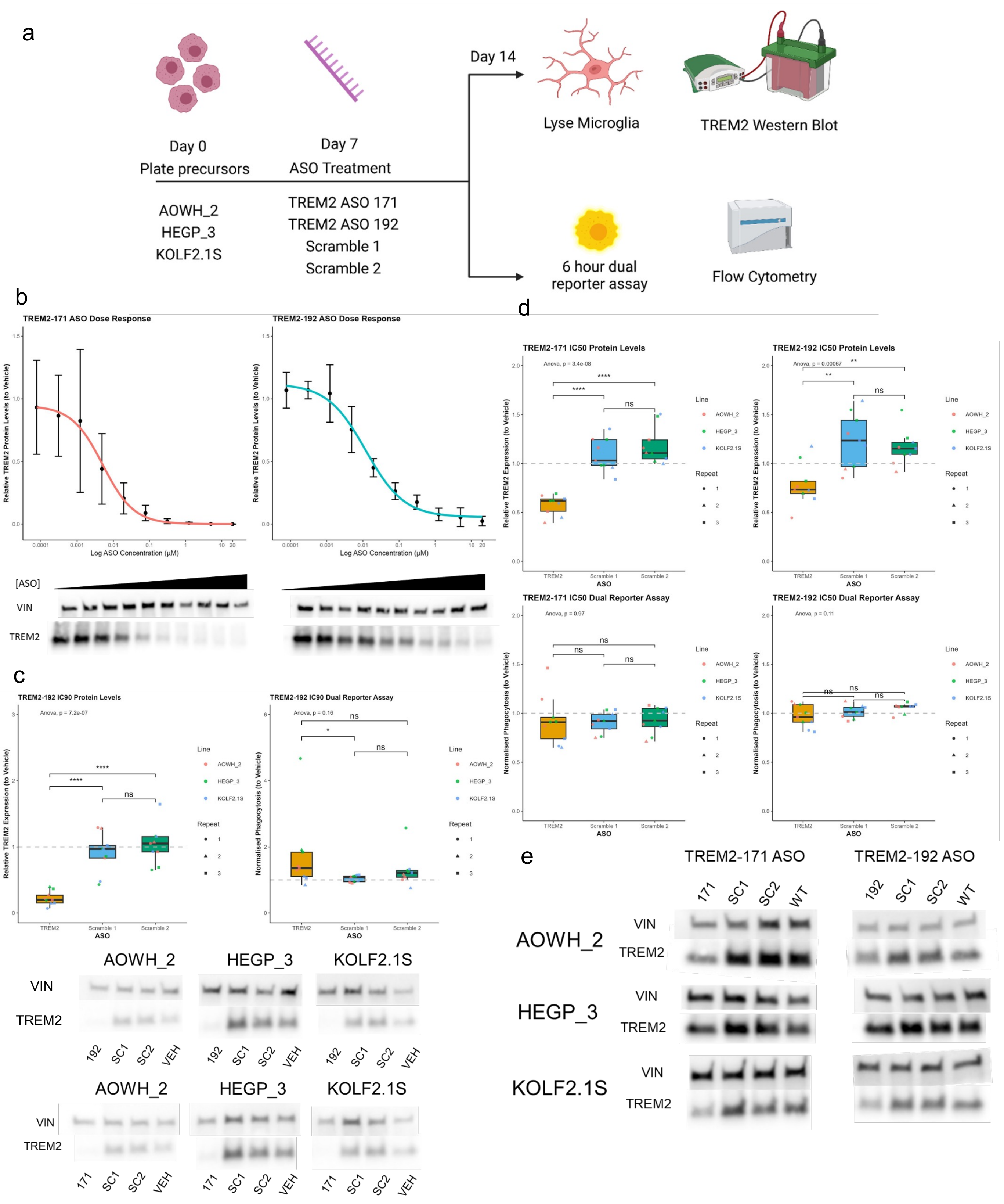

**Supplementary Figure 21:** Validation of TREM2 phenotype with ASO. a) Overview of the validation methodology. b) Inhibitory concentration curves for the two ASO TREM2-171 and TREM2-192 and representative western blots for TREM2 and Vinculin (used as a loading control). c) The same phenotypes are observed using TREM2-192 as shown using TREM2-171. A reduction of TREM2 protein at IC90 and increased phagocytic activity. d) IC50 values were also tested, while significant decreases in TREM2 expression are observed with both targeting ASO, there was no change in the phagocytosis. e) Relative western blots of TREM2 at IC50 for both ASO in the three cell lines, VIN = vinculin loading control.

#### Supplementary Figure 22

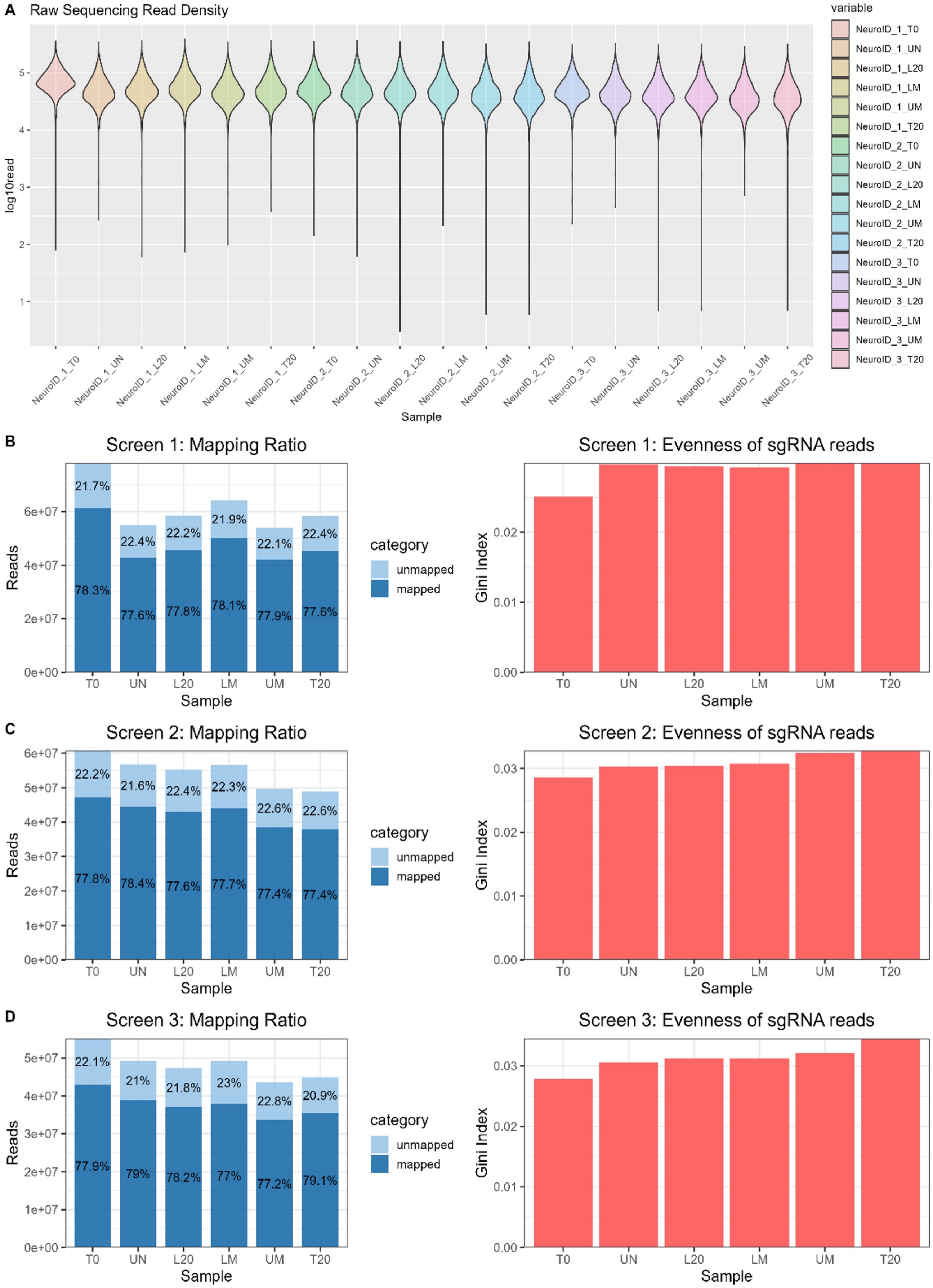

**Supplementary Figure 22:** Quality control of the sequencing data. a) Violin plots of the mapped reads to the sgRNA indicates strong coverage of the library across all populations. b-d) Over 75% of all sequence reads mapped to a sgRNA within the library with very few drop-outs according to the Gini indexes for the individual repeats of the screen, indicative of high coverage of the library.

### Supplementary Figure 23

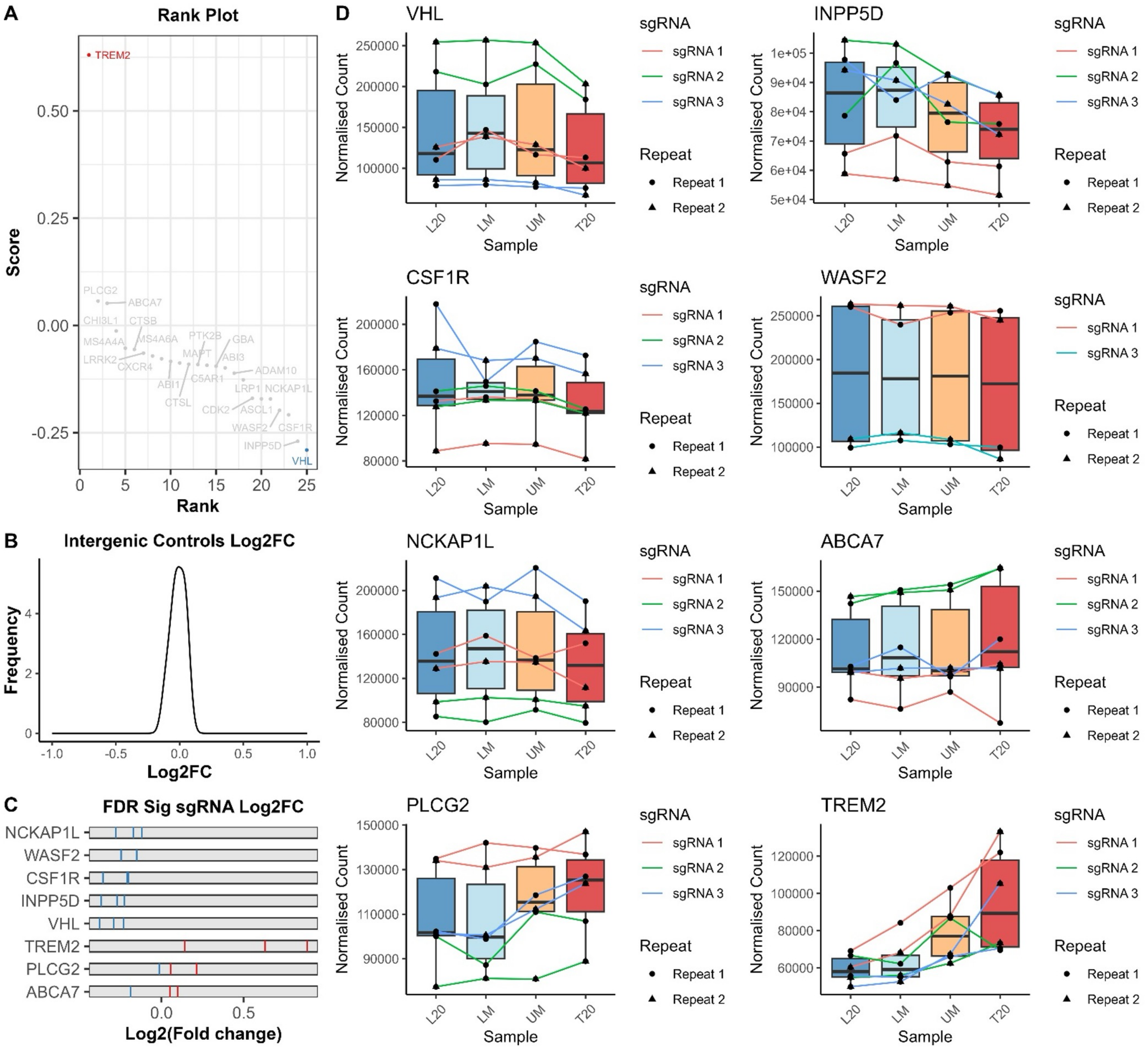

**Supplementary Figure 23:** A targeted pilot screen confirms the feasibility of the screening strategy. 25 genes were taken from the literature with known strong effects on phagocytosis from previous screens for a proof of concept screen. Three single guides were designed and cloned into the lenti-all-in-one CRISPR backbone and transduced into precursor cells and processed as outlined in Washer et al 2025. a) A rank plot of the genes identified that the selected controls moved in the expected direction – such as WASF2, NCKAP1L – known knockouts reduce phagocytosis. b) Normal distribution of the intergenic cutting control guides c) The directionality of the three sgRNA vectors across the FDR significant hits. d) Normalised sgRNA count data from MAGeCK across the four sorted bins confirms directionality and that the hit are not due to sampling biases
